# Pathogenic microbiota disrupts the intact structure of cerebral organoids by altering energy metabolism

**DOI:** 10.1101/2024.09.11.612577

**Authors:** Melis Isik, Cemil Can Eylem, Kubra Erdogan-Gover, Pinar Aytar-Celik, Blaise Manga Enuh, Emel Emregul, Ahmet Cabuk, Yalin Yildirim, Emirhan Nemutlu, Alysson Renato Muotri, Burak Derkus

## Abstract

This study investigated the impact of different bacterial populations on the biomolecular structures of cerebral organoids (COs) at various levels. COs were co-cultured with non-pathogenic (NM) and pathogenic (PM) bacterial populations. PM reduced the number of TUJ1+ neurons and disrupted the intact structure of COs. In addition, PM was found to induce changes in the transcript profile of COs, including a decrease in the activity of the glycolysis pathway and an increase in the pentose phosphate pathway, leading to deterioration in cellular energy metabolism, which is linked to neurodegenerative diseases. Proteomic analysis revealed a unique cluster of proteins in COs. PM exposure upregulated proteins related to neurological diseases, consistent with RNA-seq data. Communication between bacteria and neural cells was demonstrated using ^18^O-stable isotope labeling (SIL)-based metabolic flux analysis. COs showed higher ^18^O-enrichment of TCA cycle intermediates when co-cultured with NM and PM, indicating increased oxidative phosphorylation activity upon exposure to bacteria. This study provides a useful platform to monitor metabolic signals and communication between microbiotas and human brain cells. The findings suggest that pathogenic bacteria release metabolites that alter biomolecular structures in brain organoids, potentially contributing to neurodegenerative diseases.

## 1. Introduction

Recent studies have consistently reported that gut microbiota can modulate behavior and the function of the nervous system in rodents^1^. Although the mechanism remains unclear, findings indicate that the gut-brain axis (GBA) is responsible for this effect. The GBA consists of the central nervous system, gastrointestinal system, vagal nerves, and enteric microbiota. External factors such as stress can also affect the composition of microbiota, leading to diseases such as irritable bowel syndrome (IBS)^2,3^. The gut flora produces a range of neuroactive molecules that can directly affect brain function. Changes in the gut microbiota composition due to diet, drugs, or disease affect the secretion of these metabolites^4–6^. Therefore, there is a need to establish a realistic human *in vitro* model to explore further the effects of altering microbial populations on the neurochemistry of the human brain.

Most studies investigating the mechanisms of neurodegenerative diseases are conducted on animal models. Researchers have investigated the alterations in the proteome and metabolome in the hippocampus in animal models^7–9^. However, human studies are limited due to the lack of an advanced model. Recent advances in 3D cell culture technology have allowed pluripotent and primary cells to exhibit their self-organizing potential, resulting in organoids that reflect unique morphological and functional properties of organs. Researchers recognized that organoids have the potential to create gut microbiota models, including *Helicobacter pylori* infection model using human gastric primary cell-based organoids^10^; induced pluripotent stem cells (iPSCs)-based intestinal organoid models for the assessment of interactions with *Salmonella enterica*^11^, rotaviruses^12^, and commensal and Shiga toxin-producing *Escherichia coli*^13^; *Clostridium difficile* infection model for the investigation of paracellular barrier function^14^; modeling *Cryptosporidium* infection in human small intestinal and lung organoids^15^. However, there is no published work on establishing an *in vitro* co-culture model using cerebral organoids (COs) that help understand the biomolecular alteration in the brain in response to altering microbial populations in the gut. Thus, functional and complex physiological models are still needed to combine organoids to mimic the *in vivo* GBA process and assess how microbial population alters the biomolecular profiles of brain tissue. Developing a comprehensive *in vitro* model that engages a miniature brain tissue and defined microbiota is thought to contribute to elucidating the mechanisms and the question: “How does the change in microbiota composition affect the neurochemical structure of the brain?”.

Multi-omics approach provides a deep and integrative understanding of the effects of microbiota and its secretion on the brain^16,17^. For example, metabolomic signatures of microbiota regulate brain pathways involving aromatic amino acids and neurotransmitters^18^. A link between depressive-like behavior and microbiota has also been established by harnessing multi-omics tools^19^. However, no published work aimed to decipher the mechanisms of microbiota-mediated pathogenesis in the brain *in vitro*. In this work, multi-omics tools have been harnessed to comprehensively understand the pathogenic effects of defined bacterial populations on COs. The selection of aerobic or facultative strains was made considering their reported neurodegenerative potentials (**Table S1**). In addition to widely utilized transcriptome, proteome, and metabolome analyses, the ^18^O-stable isotope labeling (SIL) strategy has been employed to trace metabolic flux from bacterial populations to COs, modulating biomolecular plasticity in COs. ^18^O-SIL approaches have been previously implemented in the labeling of cancer cells^20^, normal tissues, and living animals^21^. To our knowledge, the utilization of ^18^O-SIL to investigate metabolic flux from microbiota to organoids was never performed. Such an approach complements the results obtained with transcriptome, proteome, and metabolome analyses, and enables a collective understanding of microbiota-mediated pathogenesis in the brain.

## 2. Materials and Methods

### 2.1 Maintenance of iPSCs culture

Two iPSC cell lines were obtained for this study. One was purchased from the Coriell Institute (GM25256, USA), while the other was kindly provided by Dr. Yalin Yildirim from the Department of Cardiovascular Surgery at the University Heart & Vascular Center Hamburg in Germany. The cells were expanded on Vitronectin (Thermo Scientific, USA)-coated 6-well plates in Essential 8^TM^ Medium (Thermo Scientific, USA). Rock inhibitor (10 μM, Y-27632, STEMCELL Technologies, Canada) was included in the medium in the first 24 hours of culture. Once the cells reached confluence, they were washed with DPBS (Thermo Scientific, USA) and detached using Versene (Gibco, USA), a non-enzymatic cell dissociation reagent, to expand the cells or induce CO formation. Pluripotency (SOX2+SSEA4+) was confirmed using a fluorescent microscope (Leica DMIL, Germany).

### 2.2 Construction and characterization of cerebral organoids

COs were obtained from hiPSCs using the STEMdiff Cerebral Organoid Kit (STEMCELL Technologies, Canada) and following a well-established protocol^22^. Briefly, hiPSCs (9.000 cells/well) were seeded into ultra-low adherent plates (ULAP) in embryoid body (EB) formation medium, supplemented with Y-27632 for the first 24 hours. When the EB size reached 500-600 µm, the medium was changed to a neural induction medium. After 2 days of induction, the neuroepithelial tissues, characterized by 3D polarized neuroectodermal tissue with a dense core structure with outermost buddings, were embedded in Matrigel, and the culture was maintained in neural expansion medium for 5 days. The medium was then switched to a maturation medium for another 5 days, and the culture was maintained on an orbital shaker. *Reverse transcriptase-quantitative polymerase chain reaction (RT-qPCR):* The gene expression levels of the developed COs were characterized using RT-qPCR. Neural stem/progenitor cell markers (Nestin, PAX6), neuronal markers (TUJ1 and MAP2), and glial markers (GFAP) were examined before and after neural induction of EBs. The detailed protocol, normalization of data, and the list of primers can be found in the Supporting Information (SI).

#### Immunofluorescent microscopy

COs were fixed in 4% paraformaldehyde for 20 min at 4°C followed by washing in PBS three times. COs were then treated in 30% (wt.) sucrose overnight and embedded in a mixture of gelatin (10% wt.) and sucrose (7.5% wt.). Cryosectioning was performed with a cryostate (Leica, Germany) at 10 μm thickness. Sections were blocked and permeabilized in BSA (1% wt.) and 0.25% Triton-X, respectively. Sections were then incubated with primary antibodies Nestin (Invitrogen, MA1-110), TUJ1 (Santa Cruz, sc-51670), MAP2 (Abcam, ab32454), GFAP (Invitrogen, A-21282). Secondary antibodies used were goat anti-mouse IgG (Alexa Fluor 488, Thermo Fisher, A-11008) and goat anti-rabbit IgG (Alexa Fluor 568, Thermo Fisher A-11011). Nuclei were stained with DAPI. Microscopic observation was performed under an epifluorescent microscope (Leica DMIL model, Leica, Germany). ImageJ software was used for determining Nestin+ and TUJ1+ cells by measuring the areas of green and red fluorescence per image (approximately 1mm^2^). The quantification was expressed as the mean of three independent biological replicates.

### 2.3 Preparation of non-pathogenic and pathogenic microbial populations

Four pathogen species including *Pseudomonas aeruginosa* (ATCC 27583), *Staphylococcus aureus* (ATCC 6538), *Salmonella typhimurium* (ATCC 13311), *Enterococcus faecalis* (ATCC 29212), as well as four probiotic species, including *Lactobacillus plantarum* 299v (obtained from the Pharmaceutical Microbiology Laboratory, Hacettepe University, Turkey), *Enterococcus durans* (obtained from Microbiota Biotechnology Inc., Turkey), *Hafnia paralvei* and *Lactobacillus lactis* (previously isolated by our group), were stored at −80°C in glycerol (20%, v/v). Two days before the co-culture experiments, non-pathogenic microorganisms were pooled, and the population was denoted as NM and grown in MRS medium (Merck, Germany). Similarly, the pathogenic microorganism population was denoted as PM and grown in Nutrient broth medium (Merck, Germany) at 37 °C with shaking under aerobic conditions. All bacteria were cultured until the stationary growth phase (6-10 hours), and the concentration of bacteria was determined by OD600. The final cell density per microorganism was set to approximately 10^7^ CFU/mL in both NM and PM.

### 2.4 Exometabolome analysis

To anticipate alterations in biomolecular profiles in COs and understand possible pathogenic mechanisms thereof, exometabolome analysis was conducted to reveal possible metabolic differences in NM and PM. First, the expansion kinetics of the strains were examined as explained in Section 2.3. Then, the supernatants obtained from NM and PM cultures were analyzed using gas chromatography-mass spectrometry (GC-MS, GC-MS-QP-2010 Ultra system, Shimadzu, Japan)^23,24^ to obtain exometabolomic profiles. Additional notes for the GC-MS analysis can be found in the Supporting Information.

### 2.5 Establishing the organoid-microbiota co-culture system

To investigate the potential impact of microbial secretome on the biomolecular profiles of COs, a transwell system with two compartments separated by a 0.4 μm membrane was utilized. This system allowed for molecular diffusion and vesicle trafficking. COs (n=30) grown in Matrigel in 6-ULAPs were transferred into the top chamber with maturation medium, while NM or PM was inoculated into the bottom chamber with MRS or Nutrient broth medium. The co-culture was maintained for up to 24 hours at 37 °C, 5% CO_2_, and 95% humidity. COs co-cultured with NM (NM_COs) and PM (PM_COs), as well as non-co-cultured COs (negative control, Cntrl_COs), were removed from the insert and prepared for downstream analyses. Each group, namely Cntrl_COs, NM_COs, and PM_COs consisted of 30 organoids, and three biological replicates were used for each group.

### 2.6 Transcriptome analysis

To uncover the transcriptomic similarities and differences in NM_COs and PM_COs in comparison to Cntrl_COs, we conducted an RNA-seq study using NovaSeq 6000 (Illumina, USA). RNA was extracted using the RNeasy mini kit (Qiagen, USA), and messenger RNA was purified from total RNA using poly-T oligo-attached magnetic beads. Raw data of FASTQ format were initially processed through in-house Perl scripts to obtain clean data by removing reads containing adapter, poly-N, and low-quality reads.

### 2.7 Proteome analysis

To explore the impact of altering microbial populations on the proteomic makeup of COs, we removed COs co-cultured with NM and PM from the transwell systems. We then treated them with a mixture of methanol and water (9:1, v/v), dissolving protein pellets with ammonium bicarbonate (100 mM) that contained 20% methanol. After a rigorous vortexing, we reduced the samples with dithiothreitol (DTT, 10 mM) for 30 minutes at 56°C. The samples were then cooled to room temperature and treated with iodoacetamide (IAA, 55 mM) in the dark at room temperature for 30 minutes. The reduced and alkylated protein samples were then digested with trypsin (1:30 w/w) for 8 hours at 37°C. Finally, we evaporated the samples to dryness and diluted them in a 200 µL of water/acetonitrile mixture (1:1, v/v) containing formic acid (FA, 0.1%).

Proteomic analyses were performed by LC-qTOF-MS on a C18 column (150 × 1 mm, 3.5 μm, 300 Å, Agilent, USA) with buffer A (0.1% formic acid in water) and eluted with a 145 min gradient time, with 2% to 90% buffer B (0.1% FA in acetonitrile) at a flow rate of 0.15 mL/min. Agilent MassHunter Workstation Software Q-TOF B.08.00 was used to run the Agilent 1260 Infinity series LC and 6530 accurate-mass LC**–**qTOF-MS instrument in auto MS/MS positive ionization mode. The run time was 180 min in gradient elution mode. The MS/MS scan data were collected in the *m***/***z* range of 300**–**1700. Peptide fragmentation was performed via the following collision energy formula; slope: 3.6 and offset: −4.8. Reference mass correction was achieved using a tuning mix (Agilent 85000 ESI-TOF Tuning Mix, Agilent, USA).

### 2.8 Untargeted metabolome analyses

To uncover the changes in metabolomic profiles in response to microbial population alterations, we subjected Cntrl_COs, NM_COs, and PM_COs to GC-MS analysis, as outlined in Section 2.4 and detailed in the Supporting Information. A total of 90 organoids per group were analyzed, with 30 organoids for each co-culture experiment and three replicates in each group. At the end of the 24-hour co-culture period, 30 organoids were carefully removed from the transwell system, and metabolites were extracted using a methanol-water mixture (1 mL, 9:1 v/v). The samples were then rapidly frozen in liquid nitrogen to halt cellular metabolism. The frozen tissues were scraped off into Eppendorf tubes, and the extracts were centrifuged at 15,000 rpm for 10 minutes at 4 °C. Next, 400 µL of each supernatant were transferred to two Eppendorf tubes and completely dried by a vacuum centrifuge for GC-MS analysis. Additional notes for the GC-MS analyses can be found in the Supporting Information.

### 2.9 Metabolic flux analysis

Before co-culture, we investigated the effects of ^18^O-SIL on microbial expansion. This involved incubating the strains with H_2_[^18^O] (30% wt.) for up to 24 hours and measuring cell density using a UV-spectrometer (Multiskan Microplate Spectrometer, Thermo Fisher, US). We also delved into the time-dependent labeling efficiency of NM and PM (final cell density ∼10^7^ CFU/mL per organism) with H_2_[^18^O] by incubating them for varying time points (0.5, 4, and 24 hours) and quantifying the labeling efficiency for each time point using GC-MS. It is important to note that the labeling optimization study was peculiarly performed on a 24-well plate to assess the suitability of microbial cultures for the co-culture system.

To initiate the metabolic flux experiment, we inoculated labeled NM or PM into the bottom chambers of 24-well transwell systems in population-specific media supplemented with 30% H_2_[^18^O], while unlabeled COs (30 organoids per insert) were placed into the upper chamber in organoid maturation medium. COs co-cultured with blank (unlabeled) NM or PM media without microorganisms were used as negative controls. After a 4-hour co-culture period, we carefully removed the COs from the inserts, washed them with DPBS, and flash-frost them in liquid nitrogen to immediately halt metabolic activity. All experiments were performed in triplicate, creating a rigorous study.

The extraction of metabolites and subsequent GC-MS analysis were conducted using the detailed methodology outlined in Sections 2.4 and 2.8, respectively. To calculate the ^18^O-labeled metabolites, we employed the techniques utilized in our previous studies^25,26^. Further details, including sample preparation, data analysis, and calculation of 18O-labeled metabolite ratio, were provided in the Supporting Information.

### 2.10 Statistical analysis and bioinformatics

#### Transcriptome data analysis

Differential expression analysis (DEA) of two-group comparisons (two biological replicates per condition) was performed using the DESeq2 R package (1.20.0). The resulting *p*-values were adjusted using Benjamini and Hochberg’s approach for controlling the false discovery rate. Genes with an adjusted *p*-value <0.05 were assigned as differentially expressed. Gene Ontology (GO) enrichment analysis of differentially expressed genes was implemented by the cluster Profiler R package.

#### Proteome data analysis

MaxQuant (Max Planck Institute of Biochemistry) was used for computational proteomic analysis of MS/MS data. Peptide length was restricted to 7 amino acids, and peptides exceeding 4600 Da were discarded. Raw MS/MS data were compared against the Uni-Prot Human Database (Taxonomy ID: 9606) for identification. Statistical analysis was performed with the Perseus platform. In the Perseus platform, the reverse, potential, only identified by site proteins and missing values between groups were filtered from the data matrix. The LFQ values of the proteins were transformed to Log 2. To verify, if the data is normally distributed, the histogram plots were generated, the intensities were compared and the statistical calculations (mean, standard error, and p-values) were performed.

#### Metabolomics data analysis

MS-DIAL software (ver. 4.48) was utilized to perform deconvolution, peak alignment, and metabolite identification. Retention index libraries employed peak annotation. TIC normalization was performed on the raw data set, and mean scaling was applied to the identified metabolite list in each group. Statistical analysis was conducted in Metaboanalyst 5.0, with 50% of the missing values excluded from the data matrix. PLS-DA, heatmap, one-way ANOVA, volcano, and pathway impact plots were generated, and the statistical calculations (mean, standard error, and p-values) were performed for each group.

## 3. Results and Discussion

### 3.1 Construction and characterization of cerebral organoids

The generation of COs from hiPSCs entails a multi-step process encompassing the formation of EBs, neuroepithelial induction, expansion, and neuronal maturation (depicted in **Figure 1A**). The morphological changes of the COs were monitored throughout induction, spanning approximately 17 days. Following the initiation of iPSCs with EB medium, the formation of EBs (measuring 500-600 μm) was verified by visualizing structures with prominent margins and a clear central region (as shown in **Figure 1B**). Upon exposure to a neural induction medium, the EBs exhibited thicker epithelial tissue and a dense core region, indicative of neuroectodermal differentiation. The formation of COs (measuring 2-3 mm) was confirmed by the presence of outermost convoluted structures and a folded neuroepithelium-like surface (**Figure 1B**), which were macroscopically visible and displayed the expected spherical morphology.

**Figure 1.**
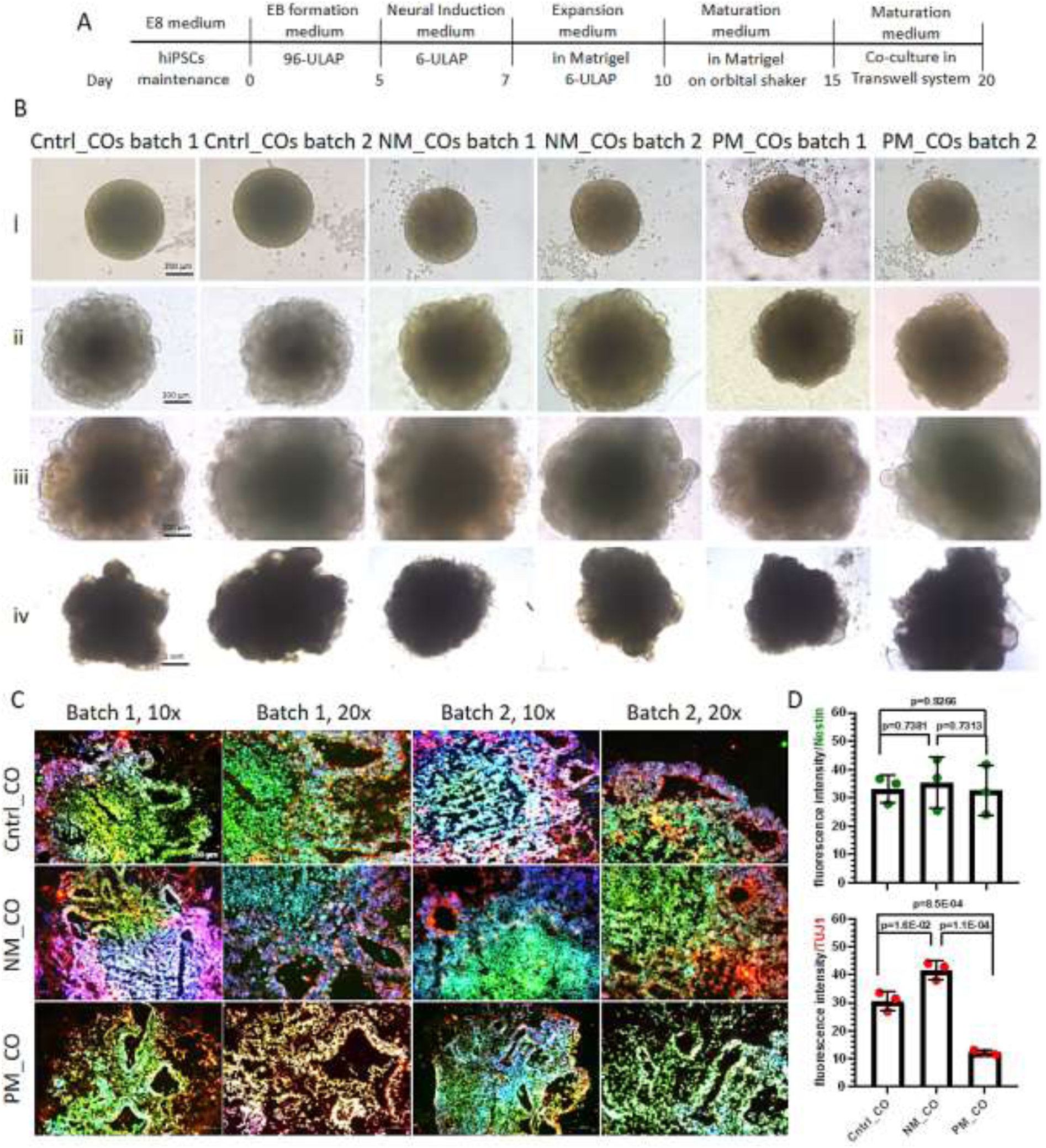
Generation of COs. (A) COs were constructed following the protocol which involves EB formation, neural induction, neural expansion and maturation, and co-cultured with microbiotas in a transwell system after 5 days of maturation and up to 5 days. (B) Inverted microscope images showing each step of Cntrl_COs, NM_COs, and PM_COs formation, provided for two batches of COs. (C) Immunofluorescent images of Cntrl_COs, NM_COs, and PM_COs, stained by Nestin (green), TUJ1 (red) and DAPI (blue). (D) Fluorescence intensity for Nestin and TUJ1, calculated by ImageJ per mm^2^ of images (n=3). Data are shown as mean ± SD of three independent replicates using two iPSCs with different genetic backgrounds. *p* < 0.05 was considered statistically significant.

The levels of expression of various neural markers, such as Nestin, PAX6, MAP2, TUJ1, and GFAP, were quantified at different time points and developmental stages of the brain organoids during culture. The expression of Nestin and PAX6, two markers of neural stem/progenitor cells, underwent a significant increase of 5.5-fold (*p = 0.00012*) and 3.5-fold (*p = 0.0019*), respectively, following the neural induction of EBs (**Figure S1**). The expression of MAP2 and TUJ1, two neuronal markers, increased by 8.1-fold (*p = 0.00014*) and 5.7-fold (*p = 0.0043*), respectively, compared to EBs, signifying the progressive differentiation of neural progenitor cells into neurons (**Figure S1**). Furthermore, it was observed that the levels of MAP2 and TUJ1 continued to increase by 2- and 3-fold, respectively, in a steady manner throughout the culture period, from day 10 to 20. The expression level of GFAP was relatively low (2.3-fold compared to EBs, *p = 0.0054*), and no significant difference in GFAP expression was discernible between neuroectoderm and EBs (**Figure S1**).

The COs before and after co-cultured with NM and PM were further examined at the phenotype level by immunofluorescent staining. To investigate the presence and structure of tissues containing neural stem cells and neurons, the Nestin and TUJ1 staining on two separate batches of COs obtained from two distinct iPSC lines was conducted **(Figure 1C)**. Cntrl_COs displayed Nestin+ and TUJ1+ cells emerging the specific morphology of Nestin+ cell enriched ventricular zone as well as TUJ1+ cells enriched outermost layer, complying with the specific histology of COs^22^. NM_COs contained multiple ventricular regions with a lateral arrangement similar to Cntrl_COs, while the outer region containing TUJ1+ neurons was observed to be broader. In contrast, in PM_COs, the ventricles are not longitudinally laterally arranged like in Cntrl_COs and NM_COs, but rather arranged in a triangular or rectangular geometry. Additionally, the regions containing TUJ1+ neurons were shown to be damaged. Additional images showing single-staining micrographs can be found in the Supporting Information (**Figure S2-4)**. ImageJ software was used to measure the amount of Nestin+ and TUJ1+ cells in each group (**Figure 1D**). There was no significant difference (*p>0.05*) in the amount of Nestin+ cells among the three groups. However, the amount of TUJ1+ cells was found to be highest in NM_CO and lowest in PM_CO (*p<0.05*).

### 3.2 Exometabolomics for non-pathogenic and pathogenic microbial populations

Before initiating co-culture experiments, we conducted untargeted metabolomics analysis using GC-MS to identify the metabolite profiles of NM and PM, to shed light on possible neuropathogenesis in COs when co-cultured with bacterial populations. A comprehensive analysis of the metabolites obtained from the supernatants of NM and PM is presented in **Table S2**. The PLS-DA score plot revealed a clear distinction between the metabolite profiles of NM and PM (**Figure S5A**). This differentiation in the metabolite profile of NM and PM was further visualized by a heatmap with hierarchical clustering analysis, as demonstrated in **Figure S5B**. The most significantly altered 15 metabolites, which were enriched in the PM exometabolome compared to that of NM, were identified in the VIP graph (**Figure S5C**). The t-test and volcano plot (p*<0.05, fold-change>2*) were generated to reveal the significant metabolites between groups (**Figure S5D-5E**). The metabolites that contributed to the separation of the two groups at higher levels included thymine, D-Alanine, L-isoleucine, galactinol, 4-hydroxyphenyl acetic acid, lactulose, fructose, N-ethylglycine, creatine, 3-phosphoglyceric acid, adenine, pyridoxine, uracil, nicotinic acid, and fumaric acid (**Figures S5C-5E**). The potential pathways that might be affected by the altered metabolites mainly involve galactose metabolism (impact 0.12), pentose-phosphate pathway (impact 0.27), pyrimidine metabolism (impact 0.26), β-alanine metabolism (impact 0.40), and ascorbate and aldarate metabolism (impact 0.50).

### 3.3 Distinct microbial populations alter transcript profiles in COs

To elucidate the molecular signatures of pathogenic bacterial population-mediated pathogenesis, we subjected COs to NM and PM in diverse transwell systems. We hypothesized that the pathogenic bacterial population secretome diffuses through the pores of the transwell system, altering the transcriptomic structure of COs and potentially leading to neuropathogenesis. To validate this hypothesis, we first confirmed the expansion kinetics of each strain in 24-well plates (**Figure S6**). We then inoculated NM or PM into the bottom chamber of a transwell system, with COs (30 organoids per well) placed in the top chamber. The COs were co-cultured with NM (referred to as NM_COs) or PM (referred to as PM_COs) for 24 hours. The control group (Cntrl_COs) was utilized to assess differences in gene expression across groups.

Principal components analysis (PCA) obtained from the RNA-seq data (**SI Excel I and II**) showed a clear separation of the groups. PC1, explaining 37.8% of the variance, highlighted the differences between Cntrl_COs and NM/PM_COs (**Figure 2A**). PC2, accounting for 24.9% of the variance, exhibited transcriptional differences between Cntrl_COs and NM/PM_COs, as well as slight differences between NM_COs and PM_COs. The Venn diagram illustrates the number of differentially expressed genes (DEGs) across groups (**Figure 2B**). 47% of transcripts were shared by all groups, while 10.6% (2337 transcripts) were up-regulated in PM_COs and 6.7% (1473 transcripts) were up-regulated in NM_COs. Interestingly, only 2.8% of transcripts (632 transcripts) were shared by NM_COs and PM_COs, contradicting the PCA results. Based on the Venn diagram, it can be concluded that the similarity in the transcriptome between NM_COs and PM_COs observed in the PCA could be attributed to the DEGs with high log2(fold change) values, while a significant portion of the transcriptome varied between these groups. RT-qPCR validation was performed on transcripts significantly upregulated in PM_COs compared to Cntrl_COs, with reported associations with neurodegenerative diseases in the literature (**Table S3**). The RT-qPCR study confirmed the RNA-seq data for the expressions of NNat, BAG3, NOS3, and NRG1 (**Figure S7**) and supported our hypothesis that the pathogenic bacterial population might be linked to neurodegenerative processes in the brain. The DESeq2 method was utilized to analyze the RNA-seq data and identify DEGs (*Log2FC>2, −log10>1.301*), resulting in 3,804 upregulated and 2,591 downregulated genes in NM_COs compared to Cntrl_COs, and 3,834 upregulated and 3,540 downregulated genes in PM_COs compared to Cntrl_COs as depicted in **Figure 2C, D**. Several genes related to the neural system with low p-values were indicated on the plots. A set of transcripts, including DNAJB1, PPAX3, JUN, NFKB2, ZRSR2, GATA3, TET2, and PAX8, were highly upregulated in NM_Cos compared to Cntrl_COs with the highest statistical significance. Similarly, H1F0, NDRG1, CDKN1C, STRA6, HSP90AA1, and RPS21 were some of the transcripts that were highly upregulated in PM_COs compared to Cntrl_COs with the highest statistical significance. A global differential expression analysis of the transcriptomes of COs versus those co-cultured with NM_COs and PM_COs revealed four distinct clusters of genes with different patterns of expression (**Figure 2E**). The patterns in NM_COs and PM_COs diverged sharply from the pattern of Cntrl_COs but only slightly from each other, as mentioned earlier. These clusters contained several genes relevant to human neurological diseases, such as Parkinson’s, Alzheimer’s, Neuropathy, Neurodevelopmental Disorders, Autism Spectrum Disorder (ASD), and Neurodegenerative Diseases (as presented in **Figures S8-15**). Differential expression analysis of our RNA-seq data of PM_COs relative to Cntrl_COs demonstrated the upregulation of various genes in the abovementioned diseases (**Figure 2F**). These genes were selected from those previously linked to the corresponding disease in the literature. The KEGG pathway annotation map illustrated changes in gene expression levels in glycolysis/gluconeogenesis, TCA cycle, and pentose phosphate pathways, complementing previous findings (**Figure S16**).

**Figure 2.**
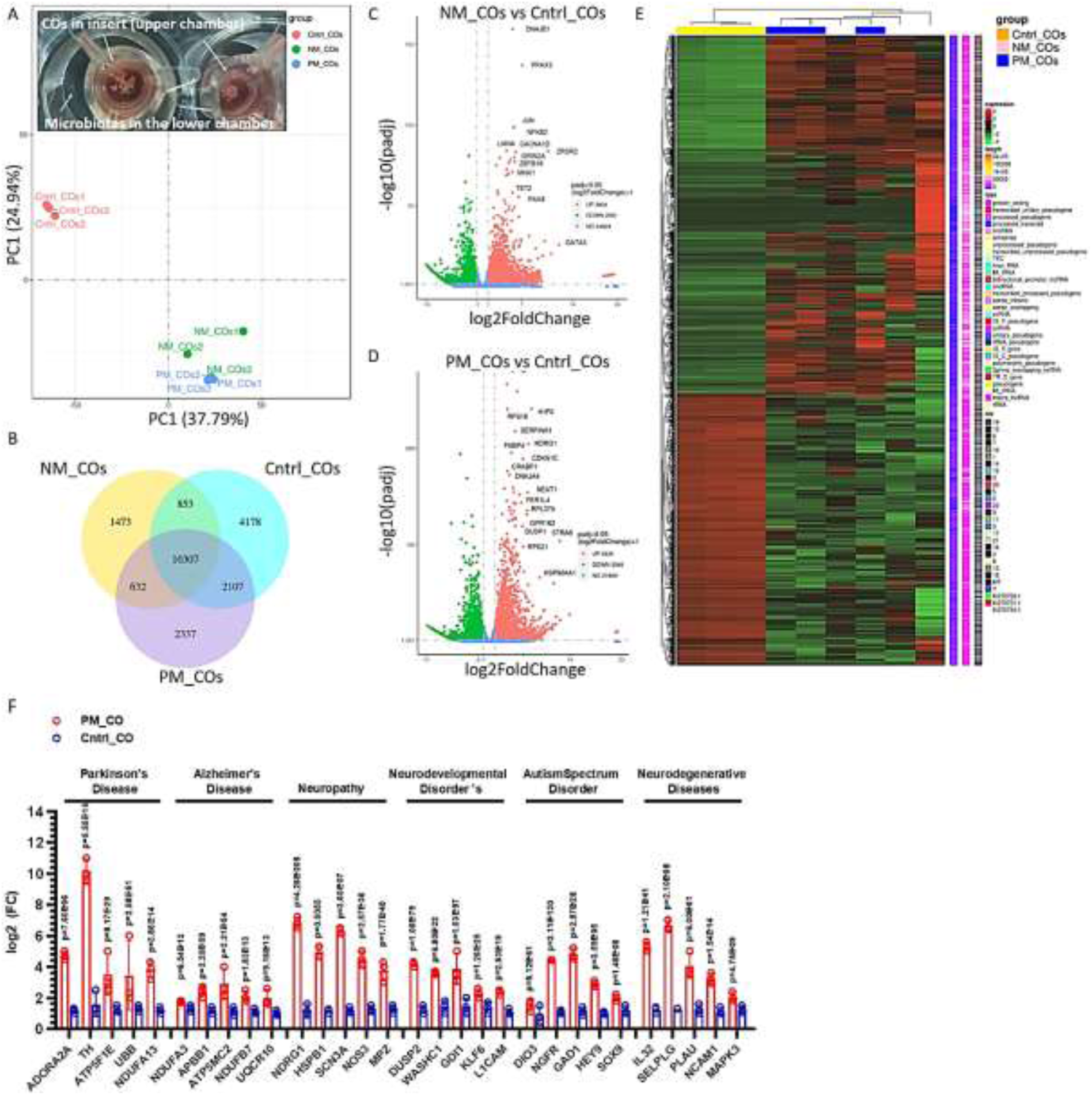
Post-microbial interaction transcript profiles in COs. (A) PCA graph demonstrating the discrimination across groups. (B) Venn scheme that shows the number of shared and differentially expressed transcripts in each group. (C, D) Volcano plot presenting the statistically significant (p*<0.05, fold-change>2*) up- and down-regulated transcripts between NM_COs and Cntrl_COs and PM_COs and Cntrl_COs. (E) Heatmap with hierarchical clustering analysis depicting the transcriptomic similarity and difference between groups. (F) Gene expression graph that shows the top 5 transcripts for each neurodegenerative disease, which most significantly up-regulated in PM_COs compared to Cntrl_COs. Analysis was performed with 30 COs for each sample, and three samples were used for each group. Transcriptome analysis was conducted with organoids obtained with a single iPS cell line.

### 3.4 Proteome analysis

A shotgun proteomics analysis was performed on the samples to complement the transcriptome data. The PLS-DA score plot (**SI Excel III**) revealed distinct clusters between the groups (**Figure 3A**). However, the sample variance was higher in PM_COs. The group variance was more apparent in the clustering analysis for the top 25 proteins and multi-scatter plot with Pearson correlation (**Figures 3B and C)**. The cluster of Cntrl_COs was noticeably distinct from the clusters of NM_COs and PM_COs, and the protein profiles in NM_COs and PM_COs showed differences at the sub-cluster level (**Figure 3B**). These variations were quantified on the multi-scatter plot (**Figure 3C**).

**Figure 3.**
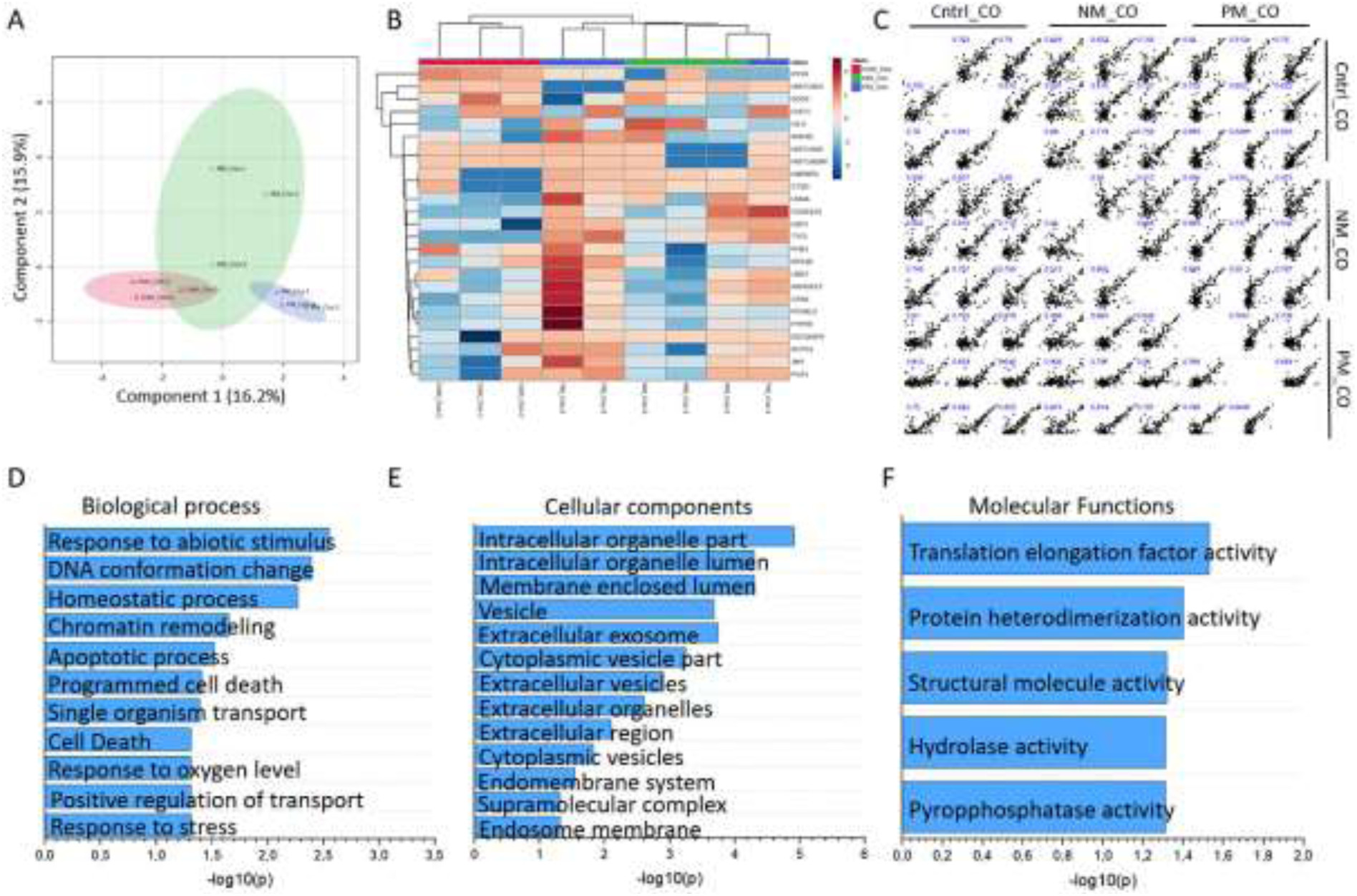
Post-microbial interaction protein profiles in COs (biological replicate=3). (A) PLS-DA score plot highlighting the discrimination in protein profiles between Cntrl_COs, NM_COs, and PM_COs. (B) Heatmap diagram with hierarchical clustering analysis depicting the proteomic signature in COs with the 25 most influential proteins. (C) A multi-scatter plot with Pearson correlation shows the variance between each sample. (D-F) Biological processes, cellular compartments, and molecular functions that are mostly affected by the most significant proteins in COs.

Among the identified proteins that exhibited up- or down-regulation between the groups (**Table S4**), several key proteins that significantly contributed to the discrimination in protein profiles were ANXA1, ANXA5, LMNA, PHB1, UBR7, UBR4, NDUFS2, ARHGEF2, CPA6, PCNXL2, PTPRE, JMY, CHIT1, APPL2, RYR2, RMDN3, EEF2, and MUC19. The biological processes, cellular components, and molecular functions affected in COs by these proteins are presented in **Figure 3D-F**. Strikingly, stress-induced processes such as response to stress, response to oxygen level, apoptotic process, and response to abiotic stimulus were highlighted (**Figure 3D**). These processes were mainly induced by extracellular cell components, such as extracellular vesicles and outer membrane vesicles (**Figure 3E**). Bacterial extracellular vesicles primarily triggered these stress-induced responses in COs through enzymatic activities (**Figure 3F**).

### 3.5 Metabolic alterations in cerebral organoids after co-cultured with microbial populations

To investigate how distinct microbial populations modulate metabolite profiles in COs, a GC-MS-based metabolomics analysis was conducted. The variance in the metabolome of COs co-cultured with either NM or PM was investigated first. The PLS-DA score plot highlighted discrimination across groups, with component 1 (17.4%) and component 2 (32.1%) (**Figure 4A**). The hierarchical clustering of metabolites that significantly changed in COs after co-culturing with NM and PM is shown in **Figure 4B**. The heatmap generated by the identified 157 metabolites based on the MS data revealed that co-culturing caused global changes in the metabolomes of COs. Importantly, 78 metabolites were found to be significantly (*p<0.05*) altered across groups (**Table S5**). Approximately 40% of the statistically significant metabolites were associated with amino acids and peptides, while approximately 20% were related to carbohydrates and carbohydrate conjugates (**Figure 4C**). To gain a clear understanding of up- or down-regulated metabolites in COs in response to NM or PM, volcano plots comparing the significantly (*p<0.05, fold change>2*) altered metabolites between Cntrl_COs, NM_COs, and PM_COs were constructed. Amino acids (e.g., 4-hydroxyproline, hypotaurine, putrescine, valine, 5-hydroxy tryptophan, beta-alanine, acetyl aspartate) and sugar metabolism-related metabolites (e.g., pyruvic acid, orotic acid, glycolic acid) were dominant in NM_COs compared to Cntrl_COs (**Figure 4D**). These significant metabolites in NM_COs only affected amino acid anabolic pathways, which mainly include arginine biosynthesis (*p=1.66×10^-4^*), tryptophan metabolism (*p=1.60×10^-3^*), glutathione metabolism (*p=2.70×10^-3^*), alanine, aspartate, and glutamate metabolism (*p=2.70×10^-3^*), and aminoacyl-tRNA biosynthesis (*p=3.20×10^-3^*) (**Figure 4E**). In the case of PM_COs, amino acid derivatives (e.g., 4-hydroxyproline, hypotaurine, putrescine, ornithine, tryptophan, and melatonin) and energy metabolism-related metabolites (e.g., creatinine and pyruvic acid) were dominant across metabolites (**Figure 4F and Table S5**). The significant metabolites affected amino acid metabolism (alanine, aspartate, glutamate metabolism (*p=1.62×10^-6^*) and arginine biosynthesis (*p=9.35 ×10^-4^*)), energy metabolism (TCA cycle (*p=1.53 ×10^-4^*) and pyruvate metabolism (*p=0.0037*)), and lipid metabolism (glycerolipids (*p=0.0223*) and pantothenate and CoA biosynthesis (*p=0.3090*)), as illustrated by the pathway enrichment analysis (**Figure 4G**).

**Figure 4.**
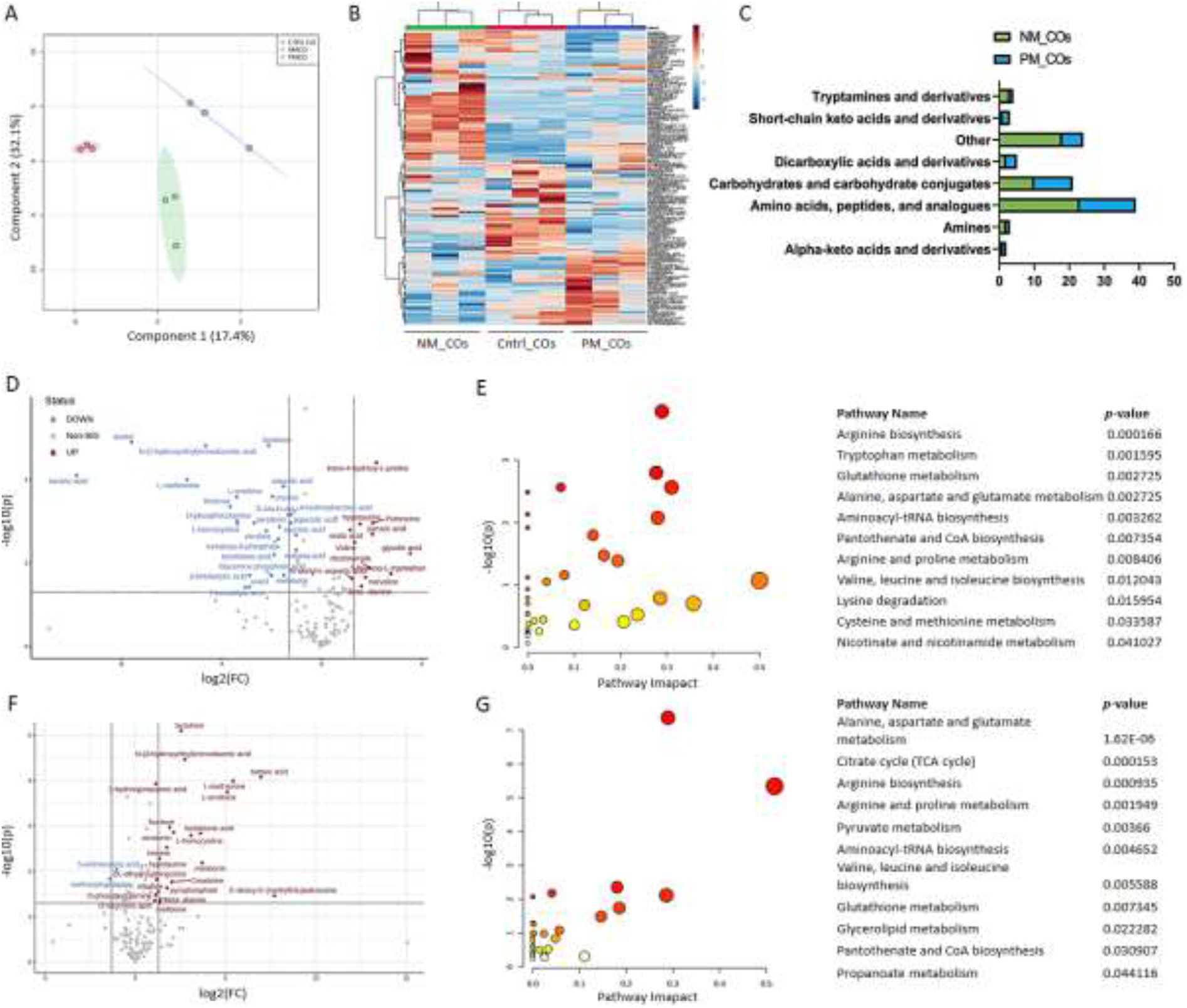
Metabolomic alterations in COs co-cultured with NM or PM (biological replicate=3). (A) PLS-DA diagram showing the discrimination between Cntrl_COs, NM_COs, and PM_COs. (B) Heatmap with hierarchical clustering analysis obtained for Cntrl_COs, NM_COs, and PM_COs depicted the discrimination in the metabolome across groups. (C) Bar chart showing the distributions of the proportions of different species in the metabolome in NM_COs and PM_COs. (D) Volcano plot showing the significant metabolites differentiate in NM_COs compared to Cntrl_COs (*p<0.05*, *fold change>2*). (E) Pathway impact graph representing the most significantly affected pathways in NM_COs compared to Cntrl_COs. (F) Volcano plot showing the significant metabolites differentiate in PM_COs compared to Cntrl_COs (*p<0.05, fold change>2*). (G) Pathway impact graph representing the most significantly affected pathways in PM_COs compared to Cntrl_COs. Statistical and pathway analyses have been conducted by MetaboAnalyst (ver. 5.0).

### 3.6 Metabolic flux analysis between microbial populations and cerebral organoids

In ^18^O-labeling studies, the rates of sequential enzymatic reactions exhibit exponential kinetics with saturation in a few minutes^21,25^. To observe the kinetic differences in phenotypes, the labeling time of metabolites should be performed within the initial linear phase of the ^18^O-labeling curve^26^. Therefore, we optimized the labeling time (30 minutes, 4 hours, and 24 hours) to determine the optimal co-culture duration in the presence of H_2_[^18^O] before saturation. The ^18^O labeling ratios of inorganic phosphate (Pi) and succinic acid, which have higher turnover rates and are involved in various cellular activities such as energy metabolism and signaling, were calculated to get an idea of the labeling saturation time. The labeling ratios increased with increasing incubation time, and Pi and succinate were observed almost to reach saturation at 24 hours. Therefore, we selected 4 hours as the optimal labeling time since the ^18^O-labeling was unsaturated and higher than that at 30 min (**Figure S17**). It should also be noted that bacteria maintained their positive growth kinetics in the presence of H_2_[^18^O] (**Figure S18**).

To quantify the level of metabolic flux from bacterial populations to COs and understand the microbial metabolite-driven metabolic changes in COs, we inoculated bacterial populations in 30% H_2_[^18^O] to the bottom chamber of the transwell system and placed unlabeled COs in the top chamber (30 organoids per chamber), and co-culture was run for 4 hours. The ^18^O-labeling ratios of 49 metabolites involved in central carbon metabolism, amino acid metabolism, and lipid metabolism, were identified and calculated by the use of fragment ions (m/z) (**Table S6**). The results showed that intermediates of central carbon metabolism (glycolysis, pentose phosphate, Krebs cycle), lipid, and amino acid metabolism pathways were communicated between NM and PM, and COs (**Figure 5**). Consistent with these data, alterations in ^18^O-labeling ratios of numerous intermediates (L-Leucine, *p=0.018*; L-isoleucine, *p=0.000*; malonic acid, *p=0.004*; L-serine, *p=0.000<*, L-valine, *p=0.000<*; L-threonine, *p=0.000<*; succinic acid, *p=0.005*; uracil, *p=0.022*; L-methionine, *p=0.012*; capric acid, *p=0.009*; malic acid, *p=0.006;* D-threitol*, p=0.000<*; phenylalanine, *p=0.003*; glycerol-1-phosphate, *p=<0.001*; palmitic acid, *p=0.000<*; stearic acid, *p=0.002*; glyceric acid, *p=0.008*; aspartic acid, *p=<0.001*; L-pyroglutamic acid, *p=<0.001*; hypotaurine, *p=<0.001*; citric acid, *p=0.033*<; allo-inositol, *p=<0.001*; methyl-beta-D-galactoside, *p=0.018*; D-ribulose 5 phosphate, *p=<0.001*) between NM and PM indicated that they play different roles and characteristics in microbiota-brain communication (**Figure S19-21**). Accordingly, the ^18^O-labeling of PM was higher for glucose-6-phosphate (G6P) and glycerol-1-phosphate (G1P) labeling ratios, indicating a strong activity of glycolytic pathway and hexokinase-catalyzed phosphotransferase pathways and glycogenolysis pathways in the PM_COs compared to NM_COs^21,25^ **(Figure S19)**. Contrarily, it was found that ^18^O-labeling ratios of Pi *(p=0.009)*, glycerol *(p=0.001)*, fumaric acid *(p=0.022)*, D-threitol *(p=0.017)*, xylitol *(p=0.045)*, stearic acid *(p=0.018)*, glyceric acid *(p=0.007)*, citric acid *(p=0.000<)*, glucose-6-phosphate *(p=0.030)*, dihydroxy acetone phosphate *(p=0.015)* and 3-phosphoglycerate *(p=0.049)* were statistically varied between NM_COs and PM_COs.

**Figure 5.**
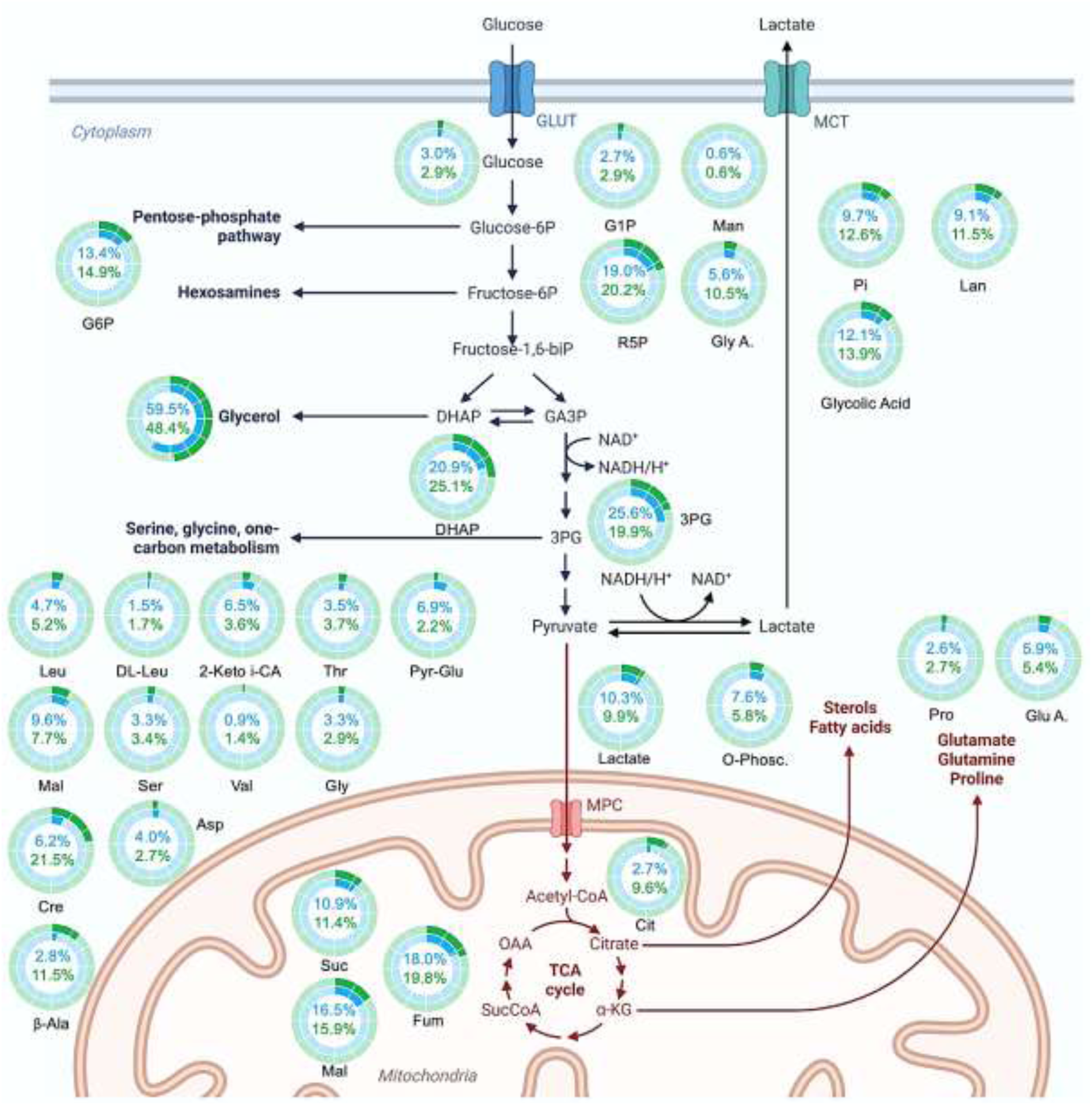
Altering metabolic flux rates in COs in response to NM and PM. Communication between the bacterial population and the COs was given as the ^18^O-labeling percentage. The blue shading represents NM_COs, whereas the green shading represents PM_COs. For instance, the percentage of ^18^O-labeled G6P transmitted from the bacterial population was 14.9 for NM and 10.3 for PM in the scenario of G6P as the sum of the labeling percentages of the pathogenic population. G1P, glycerol-1-phosphate; Man, mannitol; R5P, ribulose-5-phosphate; Gly A., glyceric acid; Pi, inorganic phosphate; Pro, proline; Glu A., gluconic acid; 3PG, 3-phosphoglycerate; o-Phosc, o-phosphocolamine; DHAP, dihydroxyacetone phosphate; Cit, citric acid; G6P, glucose-6-phosphate; Leu, leucine; DL-Leu, DL-isoleucine; 2-Keto i-CA, 2-Ketoisocaproic acid; Thr, threonine; Pyr-Glu, pyroglutamic acid; Mal, malonic acid; Ser, serine; Val, valine; Gly, glycine; Cre, creatinine; Asp, aspartic acid; β-Ala, beta-alanine.

### 3.7 Establishment and testing the hypothesis: PM disrupts the intact structure of COs by inhibiting glycolysis

According to the transcriptome and fluxome analyses, it has been concluded that PM inhibits hexokinase enzyme in COs, thereby halting glycolysis. It can be speculated that glycolytic bypass pathways might have been activated as a response, leading to the production of G3P and activating the second cascade of glycolysis. As a result, TCA has also been activated. Under normal conditions, neurons produce energy through glycolysis rather than TCA. This hypothesis is schematized in **Figure 6A**. To test this hypothesis, the expression of neuron-specific enolase, which is an isoenzyme of enolase catalyzing the conversion of 2-phosphoenolpyruvate to phosphoenolpyruvate, was analyzed using immunofluorescent imaging in response to the increase in glycolysis. It should also be noted that the level of NSE increases in the case of neurodegeneration. As expected, NSE expression was not observed in Cntrl_CO and NM_CO, while it was expressed in PM_CO (**Figure 6B**). Additionally, to test the hypothesis, the deficiency of G6P caused by hexokinase inactivation was eliminated by externally applying G6P (1 mM, every 3 days for 2 weeks), and a decreased level of NSE expression was observed.

**Figure 6.**
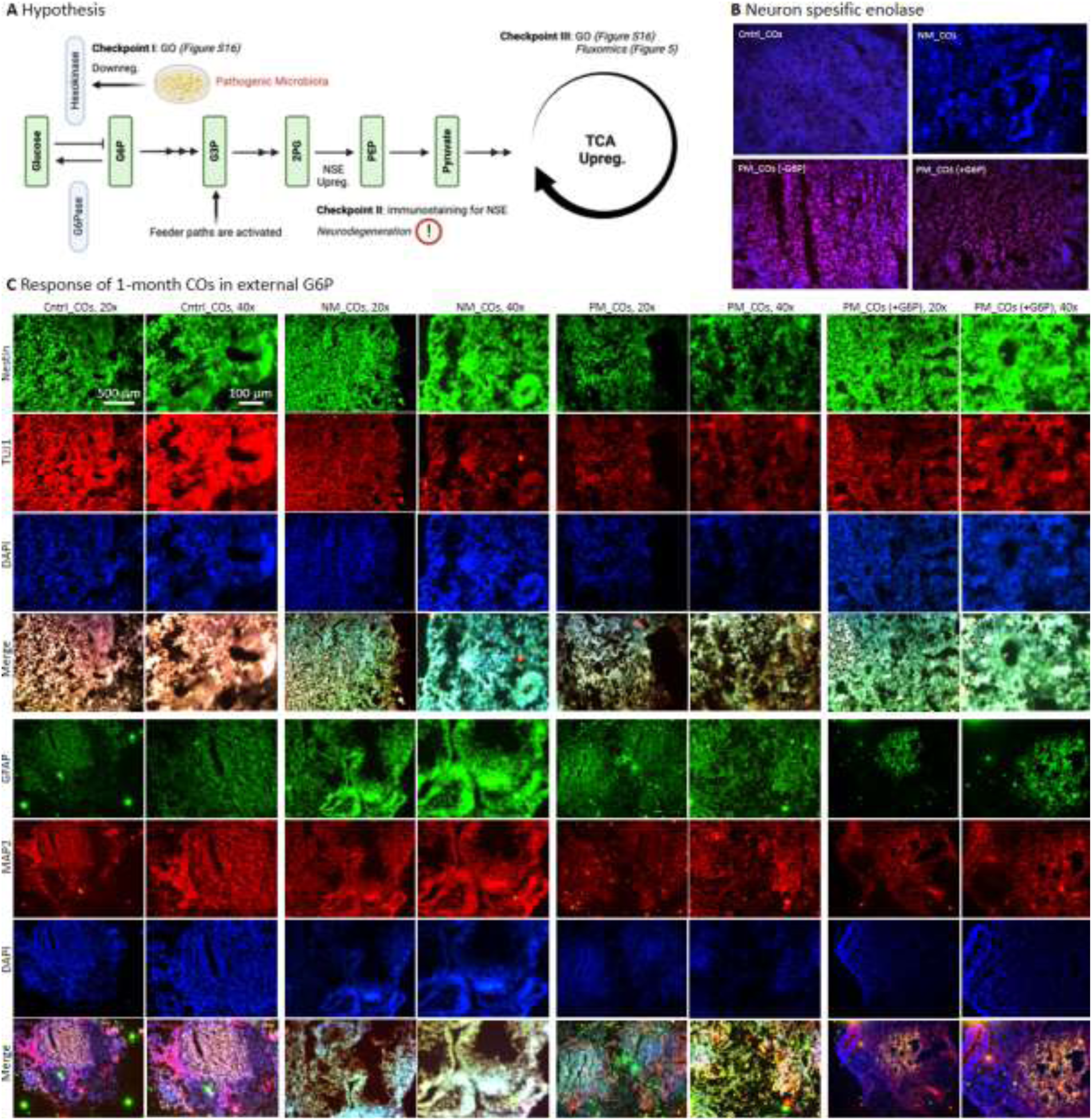
(A) Hypothesis: It is postulated that PM induces apoptosis in neurons through the inhibition of hexokinase, thereby impeding glycolysis. This hypothesis is supported by gene ontology analysis. Consequently, the TCA cycle is overactivated, as evidenced by the gene ontology analysis, due to the reactivation of glycolysis facilitated by the increased levels of G3P via alternative metabolic pathways such as lipolysis. Another crucial checkpoint for the activation of glycolysis is the expression of NSE, which is also indicative of neuronal damage. (B) Immunofluorescent imaging was employed to visualize the expression of NSE in Cntrl_COs, NM_COs, PM_COs, as well as PM_COs supplemented with G6P. (C) The assessment of neuronal damage was conducted in PM_COs with or without the provision of external G6P, in comparison to Cntrl_ COs, NM_COs, and PM_COs. One-month-old COs were examined for the expression of Nestin, TUJ1, MAP2, and GFAP, which are markers associated with neuronal development.

To initially test the hypothesis that PM inhibits glycolysis in COs and leads to neuronal cell death, COs were generated, matured for 1 month, and co-cultured with PM or NM. In addition, a part of COs was co-cultured with PM and supplemented with G6P externally. Expected CO phenotypes were observed in Cntrl_CO and NM_CO, while disrupted phenotype was observed in PM_CO (**Figure 6C)**. In the PM_CO/G6P group, the PM-mediated neurodegeneration process was halted, the phenotype was preserved, and neural apoptosis was prevented (**Figure 6C**).

## 4. Discussion

In this work, we aimed to discover how distinct bacterial populations affect brain organoids at phenotype, gene, protein, and metabolic flux levels. To reveal fundamental knowledge, we first developed COs (**Figure 1**) following a well-established method^22^, which displayed the expected forebrain identity, including ventricles with neural stem cells, in addition to differentiated neurons. As known, neurogenesis involves an increase in TUJ1+ neurons that reflect specific molecular and morphological changes that contribute to the early stages of neuronal differentiation and the maturation stage of COs^27^. The significant role of TUJ1 was confirmed by the decreased number of TUJ1+ neurons in COs exposed to metabolites from pathogenic bacterial population^28,29^. The significant increase in the number of TUJ1+ fluorescent neurons in NM_COs confirmed the development of neurogenesis and cortical region with TUJ1+ organoid maturation. The remarkable decrease in the number of TUJ1+ neurons in PM_COs revealed the inhibition of neurogenesis and brain development in organoids exposed to PM. During aging and neurogenic disorders, morphological characteristics of the SVZ region change. Consequently, the SVZ cavity widens and causes thinning of the existing neural and neuronal SVZ layers^30^. The increase in the SVZ cavity area observed in PM_COs showed that induction with PMs resulted in insufficient neural and neuronal growth in the inner SVZ region of the COs; while decreased SVZ cavity area with thick neuro-layers in NM_COs support the neurogenesis and brain development^31^.

We pooled various prebiotic and pathogenic bacteria known to play roles in neuropathogenesis to create a co-culture platform. This platform enables communication between the COs and NM or PM through the diffusion of the secretome. High-throughput tools, namely transcriptome, proteome, metabolome, and fluxome analyses, were employed to discover how bacterial populations affect brain tissue at biomolecular levels. The hypothesis that distinct bacterial populations alter transcript profiles in COs to induce pathogenesis was tested by RNA-seq. Although a large portion of the transcriptome was observed to be similar in NM_COs and PM_COs (**Figure 2A, B, E**), the modification in the transcript profile by PM_COs was found to affect several transcripts (**Figure 2C, D**) which is related to various neurodegenerative disease (**Figure 2F**). In response to co-culture with PM, we observed an upregulation of EC 3.1.3.9 (glucose-6-phosphatase) and a downregulation of EC 1.2.1.3 (glyceraldehyde-3-phosphate dehydrogenase) in the glycolysis/gluconeogenesis pathway (**Figure S16**), which catalyze the hydrolysis of glucose-6-phosphate to glucose and inorganic phosphate and the conversion of glyceraldehyde-3-phosphate to 1,3-bisphosphoglycerate, respectively. These changes suggest that COs exposed to PM experience a decrease in the activity of the glycolysis pathway, leading to the production of endogenous glucose^32^ instead of consuming G6P in the energy production pathways. This is crucial for cell survival, given the glycolysis pathway is a central pathway that produces phosphorylated precursor metabolites such as G6P. In addition, this downregulation in G6Pase might be attributed to the observed necrosis in PM_COs. Several studies have suggested that disruptions in glucose metabolism, including decreased levels of G6P, may contribute to the pathogenesis of neurodegenerative diseases such as Alzheimer’s disease and Parkinson’s disease^33^. Our transcriptome data aligns with these reports.

In the pentose phosphate pathway, we observed an upregulation of EC2.7.1.11 (phosphofructokinase) and EC2.2.1.1 (transketolase), and a downregulation of EC5.1.3.1 (ribulose-phosphate 3-epimerase). These enzymes catalyze the conversion of fructose-6-phosphate to fructose-1,6-bisphosphate in glycolysis, the transfer of a two-carbon unit from xylulose-5-phosphate to ribose-5-phosphate in the non-oxidative branch of the pentose phosphate pathway, and the interconversion of ribulose-5-phosphate and xylulose-5-phosphate in the non-oxidative branch of the pentose phosphate pathway, respectively. These changes suggest an increase in glucose utilization and a decrease in ribulose-5-phosphate production. The pentose phosphate pathway generates NADPH and ribose 5-phosphate, thereby governing anabolic biosynthesis. The changes in these pathways suggest deterioration in cellular metabolism and may linked with neurodegenerative diseases^34^.

To explain the putative neurodegenerative effects of PM at the protein level, we acquired proteomics data of COs treated with NM and PM. Proteomic analysis revealed that the proteins that contributed to proteomic differentiation across groups were ANXA1, ANXA5, LMNA, PHB1, UBR7, UBR4, NDUFS2, ARHGEF2, CPA6, PCNXL2, PTPRE, JMY, CHIT1, APPL2, RYR2, RMDN3, EEF2, and MUC19 (**Figure 3**). Among these proteins, ANXA1 has been reported in the literature to increase with increased neurodegenerative damage and neuropathological conditions^35^. In addition, ANAX5 has been previously reported to be one of the proteins most frequently observed in extracellular vesicles in Alzheimer’s^36,37^. This miR124-mediated process proceeds via the ANXA5/ERK pathway^38^. On the other hand, it is known that hippocampal LMNA is upregulated in the case of late-stage Alzheimer’s^39^. Similarly, MUC19 is known as an Alzheimer’s marker^40^. The proteins upregulated in COs after co-culturing with PM are related to neurodegenerative diseases, and overlap with the data obtained by RNA-seq.

To further investigate the systematic cascade, we utilized GC-MS-based metabolite profiling, revealing metabolic discrimination across the groups (**Figure 4**). The metabolites, which were upregulated in PM_COs, were mostly related to amino acid and lipid metabolisms. This is probably due to the adaptation of cell metabolism to the deprivation of G6P and glycolytic pathway, as well as the activation of gluconeogenesis to compensate for the lack of G6P. These results are in line with the pathway enrichment outputs obtained by RNA-seq analysis and suggest that PM suppresses glycolysis in COs, leading to the upregulation of the phosphate- pentose pathway. Furthermore, it is inferred that PM induces a shift in metabolism towards lipolysis to synthesize acetyl-CoA. In other words, PM promotes the formation of glucose and ketone bodies, which serve as immediate energy sources, by inducing cellular stress in COs. Consequently, the results obtained are consistent with previous findings, and it was determined that PM induced plasticity in organoid metabolism compared to the control group.

In light of the transwell co-culture concept, the transfer of ^18^O-labeled metabolites from NM or PM to COs suggests that metabolites play a role in the communication between bacteria and neural cells, in particular, mitochondria of neural cells, which is a novel concept^41^. Furthermore, the varying expression of ^18^O ratios moved to brain organoids via metabolites indicates that pathways linked to NM and PM have different turnover rates. In addition, the fact that COs had higher ^18^O enrichment of TCA cycle intermediates in response to co-culture with NM and PM (**Figure 5**) indicates that the activity of oxidative phosphorylation increased when COs met with bacterial populations. Furthermore, the fact that these intermediates have different ^18^O enrichments in NM_COs and PM_COs indicates that they communicate with COs through similar pathways but at different levels and rates. Thus, higher ^18^O enrichment of citric acid, fumaric acid, and Pi in COs after co-culture with PM points to PM increasing the activity of oxidative phosphorylation in COs compared to NM. In addition, the increased rate of Pi in COs after co-culturing with PM points to intracellular energy transmission occurring faster. Moreover, a higher ^18^O-labeling ratio of dihydroxyacetone phosphate, transferring from the PM to the COs, indicates a greater effect of redox balance and substrate shuttling to mitochondria in PM_COs^42^. The increased ^18^O enrichment, which is fully transferred from bacterial populations to COs, demonstrates that 3-phosphoglycerate, an intermediate metabolite in glycolysis charging in the conversion of glucose to pyruvate and ATP production, is as effective in neuronal function and neurotransmitter regulation as Pi, succinic acid, citric acid, and G6P. It is also worth noting that insignificant differences in numerous amino acids transferred from bacteria to COs suggest that amino acid transfer from microbiota to brain is permitted to some extent and that excess amino acid transmission is limited owing to regulatory and homeostasis mechanisms in the brain. To conclude, our methodology has the potential to provide a useful platform for monitoring metabolic signals and communications in future studies.

To consolidate our findings obtained with transcriptomics study (showed downregulation in hexokinase and inhibition in glycolysis in PM_COs) and fluxomics study (showed an upregulation in G6P, thus glycolysis, and TCA metabolites in PM_COs), meaning a G6P flux from PM to COs, we tested the activity of G6P in neurogenesis by providing external G6P into COs culture (**Figure 6**). The inhibition in hexokinase seen in GO analysis was expected to lead to an over-activation in alternative energy sources (TCA); i.e., upregulation in isoenzyme NSE, indicative of both the reactivated glycolysis and neuronal degeneration^43,44^. This was highly observed in PM_COs and slightly observed in PM_COs/G6P, confirming that PM_COs halts glycolysis, yet alternative paths elevate the G3P to reactivate glycolysis. When we introduced the external G6P, the cytosolic accumulation of NSE was observed at a slightly lower level, possibly due to the recovery of cell metabolism.

## 5. Conclusion

In conclusion, our study has demonstrated the impact of distinct bacterial populations on brain organoid phenotypic and biomolecular structures at the gene, protein, and metabolic flux levels. Through the use of high-throughput tools, we have identified changes in biomolecular profiles that suggest a link between bacterial populations and neurological diseases. We have also revealed the communication between bacteria and neural cells through the transfer of metabolites, indicating the potential for a new concept in neuroscience. Given that the metabolites transferred from bacterial populations mostly affect energy pathways, these interactions between bacterial populations and neural mitochondria may serve as a new focus, which could help to extend the vision of the gut-brain axis. Our methodology provides a useful platform for future studies investigating metabolic signals and communications between microbiota and the brain. This study highlights the importance of understanding the relationship between microbiota and brain health and the potential for microbiota-targeted therapies in treating neurodegenerative diseases.

## Acknowledgment

This work was financially funded by the Scientific and Technological Research Council of Turkey (TUBITAK, grant number: 219S661). B. Derkus also thanks the Turkish Academy of Science (TUBA) and the Science Academy (Istanbul) for their support.

## Contributions

B.D. developed the concept, received the grant, and supervised this study. B.D., M.I., and E.N. designed the experiments. M.I., C.C.E., K.E-G. performed the experiments. B.D., E.N., P.A-C., and E.B.M. cured and analyzed the data. Y.Y. provided cell lines and commented on the manuscript. B.D. wrote the manuscript. A.C., E.E., and A.R.M. edited and commented on the manuscript. All authors approved the final version of the manuscript.

## Data availability

The data that support the findings of this study are available from the authors on reasonable request, see author contributions for specific data sets.

