## Supplementary material for "Pathogenic microbiota disrupts the intact structure of cerebral organoids by altering energy metabolism": Supporting Information.pdf

**Content**

**Section 1** Rationale of the study

**Section 2** Characterisation of COs in gene expression and phenotype levels

**Section 3** Exometabolome analysis

**Section 4** Transcriptome analysis

**Section 5** Proteome analysis

**Section 6** Metabolome analysis

**Section 7** Metabolic flux analysis - Fluxomics

### **Section 1** Rationale of the study

The aim of this study was to establish an in vitro co-culture model of brain-microbiota and to decipher the possible molecular neuropathogenic mechanisms associated with pathological microbiota using multi-omics approaches. First, brain organoids were generated, and then microbiota pools were created by selecting four aerobic or facultative strains, which were reported as probiotic or pathological in the literature. After co-culturing the obtained cerebral organoids (COs) with non-pathogenic (NM) or pathogenic microbiota (PM) in a co-culture system, molecular changes occurring in the COs were examined. The literature studies that form the basis of the selection of the pathological strains are presented in **Table S1**.

*Table S1*. The rationale of the selection of strains

| | **Parkinson Disease** | **Alzheimer's Disease** | **Neuropathy** | **Autism Spectrum Disorder** | **Neurodegenerative Disease** |
|---|---|---|---|---|---|
| *Pseudomonas aeruginosa* | [1] | [3] | [4] | [5] | [6] |
| | [2] | | | | |
| *Staphylococcus aureus* | | | | | [7] |
| *Salmonella typhimurium* | [2] | [8] | | | [9] |
| | | | | | [10] |
| *Enterococcus faecalis* | | | | [11] | |
| | | | | [5] | |

**Section 2.** Characterisation of COs in gene expression and phenotype levels

Cntrl_COs, NM_COs, and PM_COs were characterized by RT-qPCR. To this end, RNA was isolated by a total RNA isolation kit (GeneDireX, USA), and 500 ng of RNA in each sample was converted into cDNA using a kit (Bio-Rad, USA). RT-qPCR was run on a thermal cycler (Bio-Rad CFX96 instrument, USA) using the primers (Oligomer Biotechnology Jsc., Turkey) listed below. Each quantitative experiment was performed with three replicates. The statistical difference between more than two groups was investigated by one-way ANOVA followed by Tukey's post-hoc test. Normalisation was done by using equal amount of RNA from each sample, and by comparing gene expressions with control groups. The gene expression results were presented in **Figure S1**.

**List of primers**

**Nestin:**
F-CAGCGTTGGAACAGAGGTTGG
R-TGGCACAGGTGTCTCAAGGGTAG
**PAX6:**
F-ACCCATTATCCAGATGTGTTTGCCCGAG
R-ATGGTGAAGCTGGGCATAGGCGGCAG
**MAP2:**
F-CTCAGCACCGCTAACAGAGG
R-CATTGGCGCTTCGGACAAG
**TUJ1:**
F-GGCCTTTGGACATCTCTTC
R-CTCCGTGTAGTGACCCTTG
**GFAP:**
F-CAAGATGAAACCAACCTGAGGCT
R-GGCTTGGCCACATCCATCT

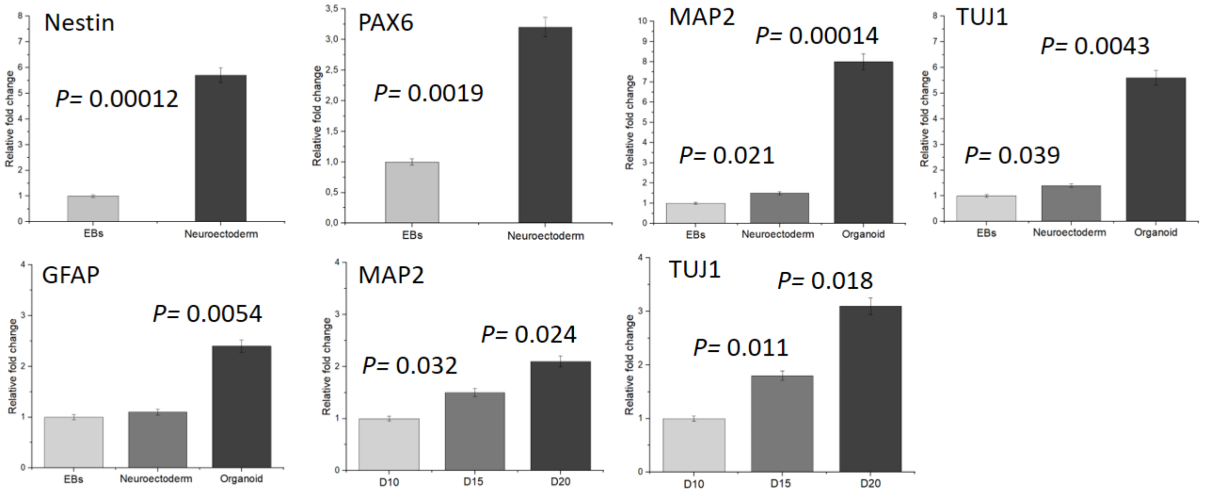

*Figure S1*. Characterisation of Cntrl_COs, NM_COs, and PM_COs in gene expression level (technical replicate=3, biological replicate=3, 10 organoids per sample). The expressions of Nestin and PAX6 were used for the assessment of neural stem cell identity, the expressions of MAP2 and TUJ1 were used for the assessment of neuronal identity, and GFAP was considered as neuroglial identity. Analyses were performed with COs after 5 days of maturation phase. Statistical significance was determined using a one-way ANOVA followed by Tukey's post hoc test (*p<0.05).

In addition to gene expression analysis, Cntrl_COs, NM_COs, and PM_COs were also characterized in phenotype level by immunofluorescent microscopy. Nestin, TUJ1, and DAPI were used to stain neural stem cells, neurons, and nuclei, respectively. The overlay images in each group were presented in the main text, additionally, we presented single staining images in **Figure S2-4** to provide a better understanding of the histological structures of COs.

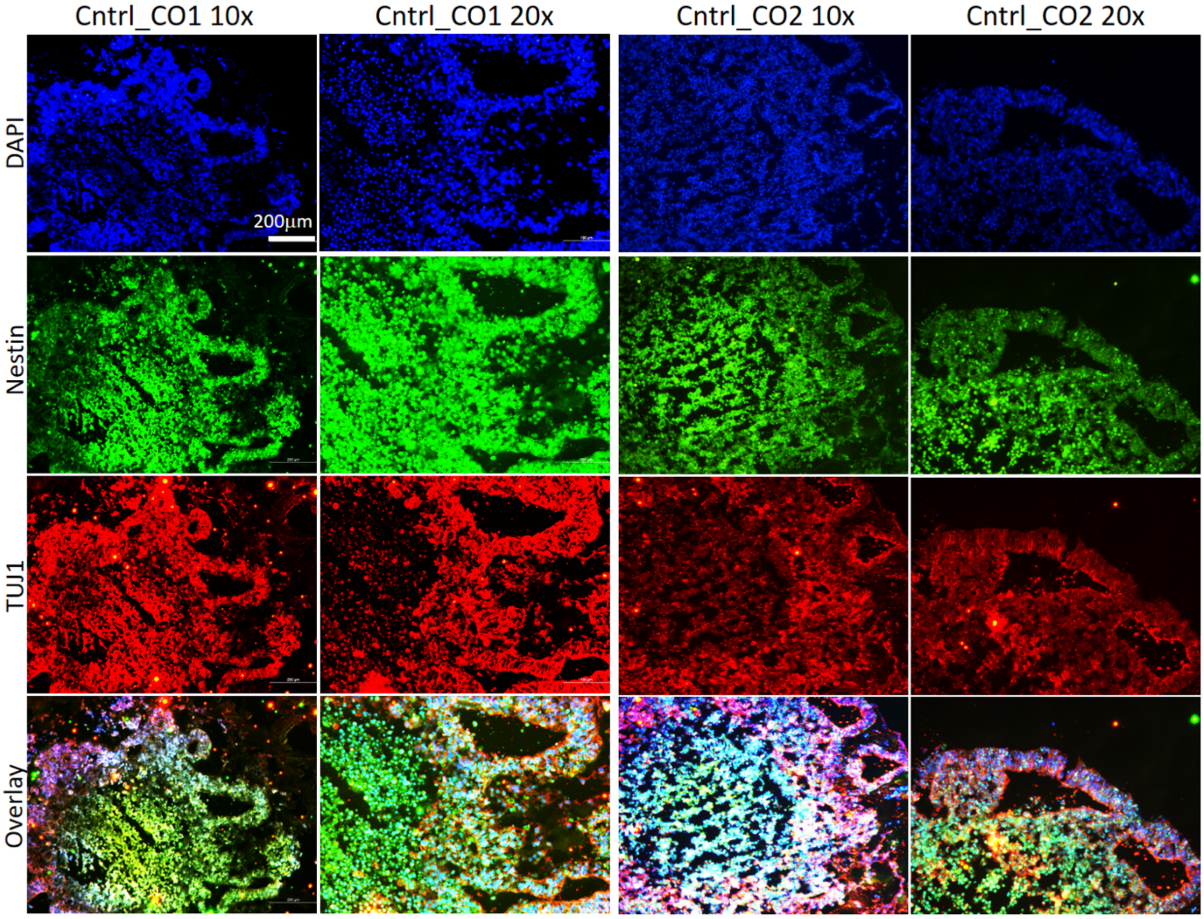

*Figure S2*. Immunophenotyping of Cntrl_COs, COs from two different batches constructed with two different iPSCs lines were stained for Nestin, TUJ1, and DAPI. Images were presented in two different magnifications.

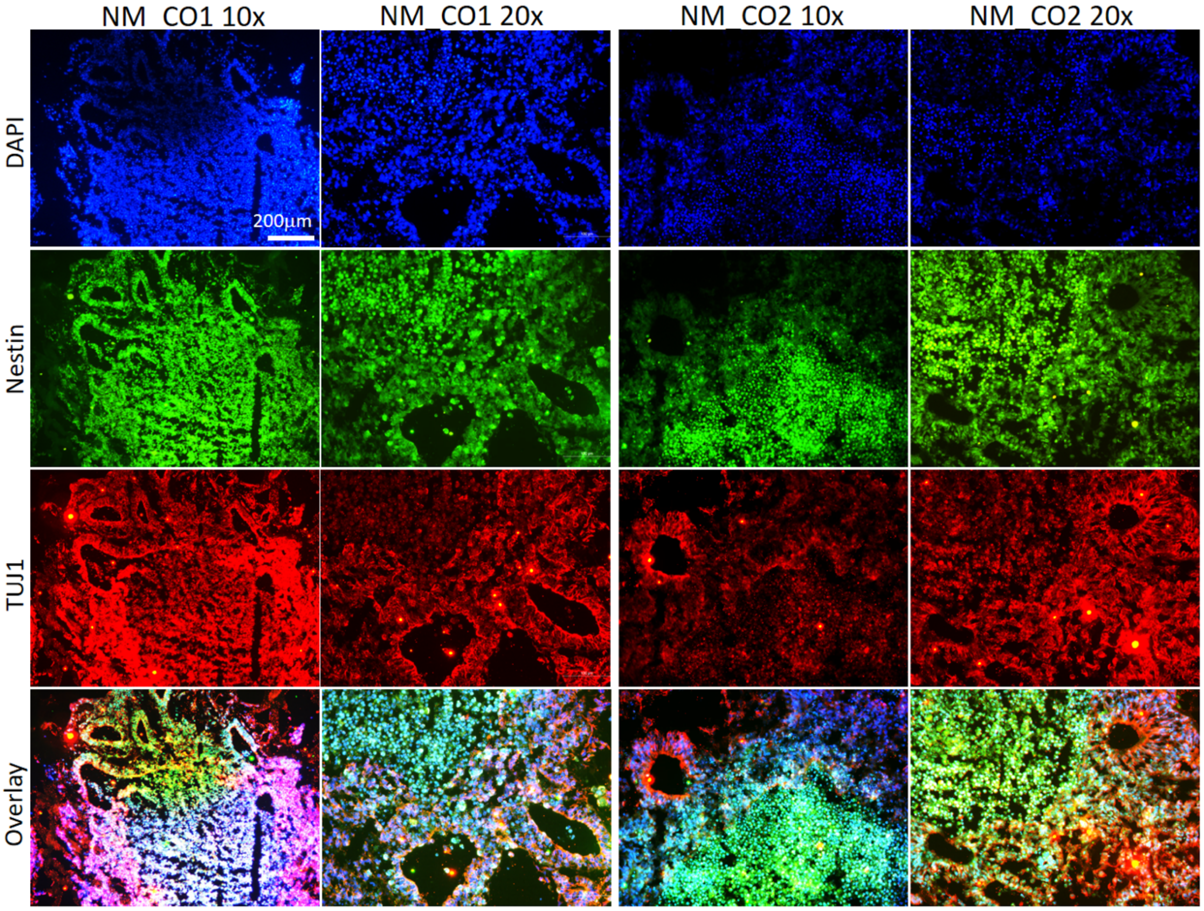

*Figure S3*. Immunophenotyping of NM_COs. COs from two different batches constructed with two different iPSCs lines were stained for Nestin, TUJ1, and DAPI. Images were presented in two different magnifications.

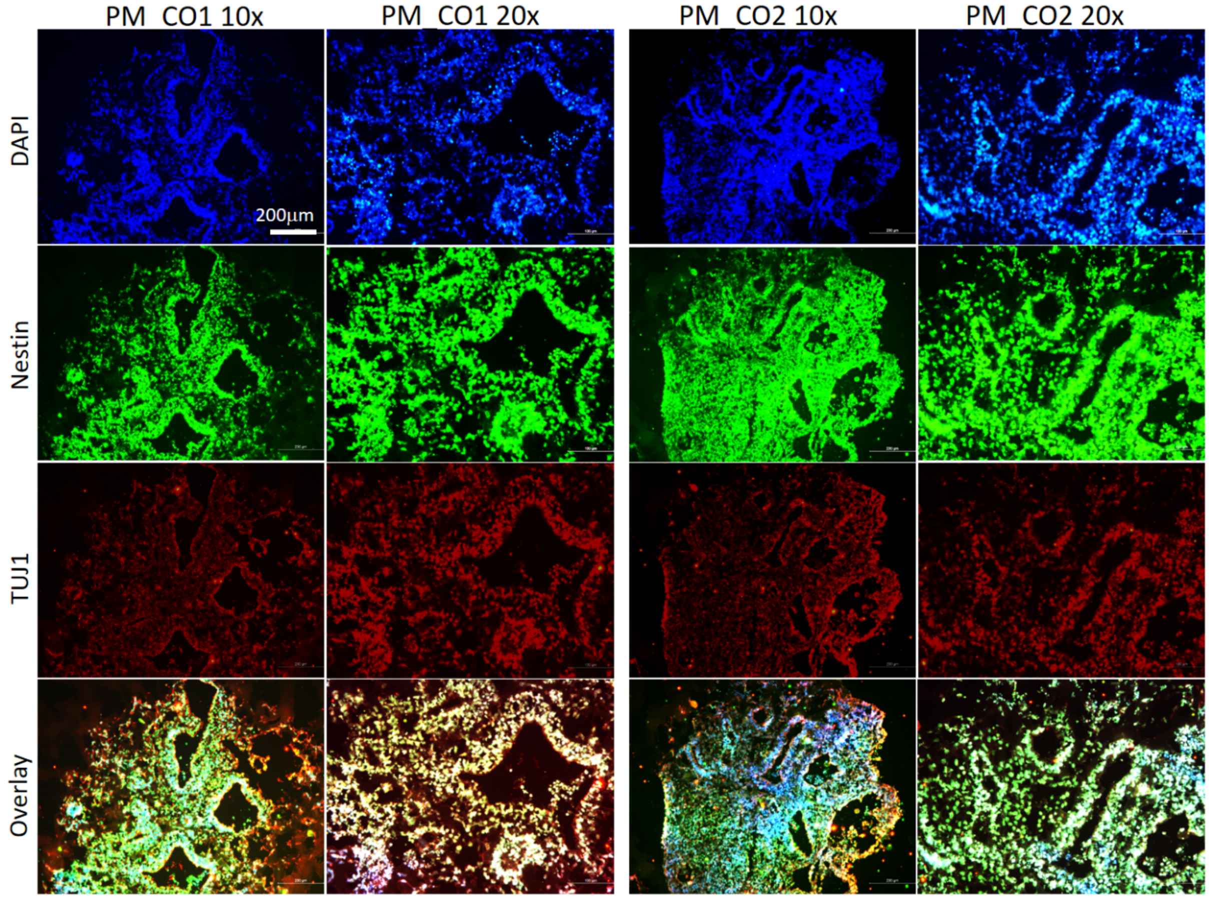

*Figure S4*. Immunophenotyping of PM_COs. COs from two different batches constructed with two different iPSCs lines were stained for Nestin, TUJ1, and DAPI. Images were presented in two different magnifications.

**Section 3.** Exometabolome analysis

Prior to delving into the co-culture experiments, we conducted an exometabolomics analysis by GC-MS to elucidate the potential impacts of microbiota secretion products on COs.

*GC-MS based metabolomics analyses*: The dried samples were methoxyaminated using 20 µL methoxyamine hydrochloride in pyridine (20 mg/mL) for 90 min at 30°C. Shortly after, the samples were derivatized using *N*-methyl-*N*-trimethylsilyl trifluoroacetamide (MSTFA) + trimethylchlorosilane (TMCS, 1%) for 30 min at 37°C. Samples were transferred to silylated vials and analyzed by GC-MS using a DB5-MS column (30 m +10 m duraguard × 0.25 mm i.d. and 0.25-µm film thickness). The run time and injection volume were set at 37.5 min and 1 µL (splitless), respectively. The MS scan data were collected in the range of 50–650 m/z using GC-

MS Solution (Shimadzu, ver. 4.20). Injection temperature and the flow rate were set at 290°C and 0.99 mL/min, respectively. The solvent delay time was adjusted to 5.90 min.

The list of exometabolites was provided in **Table S2**.

*Table S2*. Exometabolome obtained from NM and PM

| **Metabolite** | **NM** | **PM** | **p** |
|---|---|---|---|
| | **Mean ± SE** | **Mean ± SE** | |
| 1-hexadecanol | 0.12 ± 0.04 | 0.27 ± 0.09 | 0.21 |
| 1,3-diaminopropane | 0.61 ± 0.31 | 0.87 ± 0.08 | 0.46 |
| 2-hydroxybutyric acid | 4.06 ± 0.33 | 0.05 ± 0.01 | **0.00** |
| 2-ketoadipic acid | 5.56 ± 2.72 | 6.79 ± 0.1 | 0.68 |
| 2-ketocaproic acid | 2.99 ± 0.31 | 0.69 ± 0.18 | **0.00** |
| 2-ketoisocaproic acid | 0.09 ± 0.02 | 0.1 ± 0 | 0.77 |
| 2-phenylacetamide | 3.86 ± 0.61 | 0.5 ± 0.09 | **0.01** |
| 3-aminoisobutyric acid | 1.37 ± 0.69 | 11.03 ± 0.2 | **0.00** |
| 3-hexenedioic acid | 1.02 ± 0.5 | 1.27 ± 0.05 | 0.64 |
| 3-indolelactic acid | 8.25 ± 2.81 | 3.63 ± 0.06 | 0.18 |
| 3-phenyllactic acid | 9.29 ± 1.16 | 3.33 ± 0.04 | **0.01** |
| 3-phosphoglyceric acid | 0.01 ± 0.01 | 10.48 ± 0.34 | **0.00** |
| 4-guanidinobutyric acid | 1.6 ± 0.79 | 6.62 ± 0.26 | **0.00** |
| 4-hydroxybenzoic acid | 2.17 ± 0.87 | 7.58 ± 0.14 | **0.00** |
| 4-hydroxy-L-proline | 0.07 ± 0.03 | 5.41 ± 0.25 | **0.00** |
| 4-hydroxyphenylacetic acid | 0.7 ± 0.03 | 5.86 ± 0.06 | **0.00** |
| 4-hydroxyquinoline | 1.44 ± 0.03 | 0.22 ± 0.01 | **0.00** |
| 6-hydroxyhexanoic acid | 3.05 ± 0.79 | 5.91 ± 0.23 | **0.03** |
| 6-phosphogluconic acid | 0.13 ± 0.07 | 1.49 ± 0.04 | **0.00** |
| 10-hydroxydecanoic acid | 2.84 ± 2.76 | 1.28 ± 0.03 | 0.60 |
| Adenine | 0.56 ± 0.28 | 12.14 ± 0.09 | **0.00** |
| Adenosine | 1.78 ± 0.88 | 10.02 ± 0.91 | **0.00** |
| Adenosine-3-monophosphate | 2.95 ± 1.47 | 8.82 ± 0.45 | **0.02** |
| Adenosine-5-monophosphate | 1.31 ± 0.66 | 8.69 ± 0.86 | **0.00** |
| Alpha ketoglutaric acid | 1.95 ± 0.09 | 1.82 ± 0.05 | 0.28 |
| Alpha tocophereol | 0.01 ± 0 | 0.01 ± 0 | 0.41 |
| Alpha-D-glucosamine phosphate | 0.87 ± 0.3 | 0.13 ± 0.01 | 0.07 |
| Arabitol | 6.15 ± 3.32 | 0.64 ± 0.02 | 0.17 |
| Arachidic acid | 0.38 ± 0.08 | 0.86 ± 0.07 | **0.01** |
| Arbutin | 1.88 ± 0.91 | 0.47 ± 0.11 | 0.20 |
| Aspartic acid | 0.84 ± 0.33 | 1.19 ± 0.07 | 0.36 |
| Benzoic acid | 0.01 ± 0 | 0.83 ± 0.08 | **0.00** |
| Beta- alanine | 0.06 ± 0.03 | 1.01 ± 0.03 | **0.00** |
| Beta-cyano-L-alanine | 1.85 ± 0.26 | 0.39 ± 0.03 | **0.00** |
| Beta-glycerolphosphate | 1.29 ± 0.15 | 0.5 ± 0.01 | **0.01** |

| | | | |
|---|---|---|---|
| Capric acid | 0.01 ± 0 | 0.38 ± 0.01 | **0.00** |
| Cellobiose | 2.44 ± 0.35 | 0.77 ± 0.06 | **0.01** |
| Cholesterol | 0.02 ± 0.02 | 0.07 ± 0 | 0.05 |
| Citraconic acid | 0.58 ± 0.2 | 0.9 ± 0.06 | 0.20 |
| Citramalic acid | 3.24 ± 0.88 | 1.37 ± 0.03 | 0.10 |
| Citric acid | 2.85 ± 1.9 | 1.78 ± 0.08 | 0.60 |
| Citrulline | 1.39 ± 0.24 | 1.39 ± 0.07 | 0.99 |
| Creatinine | 0.01 ± 0 | 3.3 ± 0.08 | **0.00** |
| Cycloleucine | 1.15 ± 0.1 | 8.47 ± 0.98 | **0.00** |
| Cytidine | 1.02 ± 0.46 | 8.44 ± 2.51 | **0.04** |
| Cytindine-5-monophosphate | 0.18 ± 0.08 | 0.1 ± 0.01 | 0.32 |
| Cytosine | 6.52 ± 3.24 | 5.48 ± 5.37 | 0.88 |
| D (+) galactose | 0.64 ± 0.2 | 6.86 ± 0.28 | **0.00** |
| D-(+) trehalose | 9.16 ± 0.35 | 0.14 ± 0 | **0.00** |
| D-(+)-melezitose | 1.18 ± 0.55 | 9.82 ± 0.64 | **0.00** |
| D-ala-D-ala | 0.03 ± 0.01 | 12.06 ± 0.31 | **0.00** |
| Dehydroascorbic acid | 1.19 ± 0.2 | 0.47 ± 0.14 | **0.04** |
| D-glucosaminic acid | 1.44 ± 0.13 | 0.02 ± 0 | **0.00** |
| D-glucose | 0.71 ± 0.08 | 0 ± 0 | **0.00** |
| D-glucose-6-phosphate | 0.47 ± 0.13 | 0.03 ± 0.01 | **0.03** |
| Dl-threo-beta-hydroxyaspartic acid | 1.9 ± 0.97 | 7.16 ± 0.12 | **0.01** |
| D-lyxose | 0.41 ± 0.19 | 5.92 ± 0.06 | **0.00** |
| D-malic acid | 3.58 ± 0.56 | 0.2 ± 0.01 | **0.00** |
| D-mannitol | 1.56 ± 0.13 | 0.8 ± 0.05 | **0.01** |
| D-sorbitol | 3.61 ± 0.43 | 0.08 ± 0 | **0.00** |
| D-threitol | 0.39 ± 0.02 | 0.31 ± 0 | **0.01** |
| Estradiol | 1.54 ± 0.77 | 6.15 ± 0.31 | **0.01** |
| Fructose | 0.02 ± 0.01 | 1.18 ± 0.02 | **0.00** |
| Fumaric acid | 0.01 ± 0.01 | 8.35 ± 0.41 | **0.00** |
| Galactinol | 1.22 ± 0.14 | 10.96 ± 0.21 | **0.00** |
| Glucoheptonic acid | 1.16 ± 0.3 | 0.19 ± 0.01 | **0.03** |
| Gluconic acid | 1.7 ± 0.6 | 0.49 ± 0.01 | 0.11 |
| Glucuronic acid | 0.97 ± 0.08 | 0.38 ± 0.01 | **0.00** |
| Glyceric acid | 5.61 ± 0.85 | 0.99 ± 0.03 | **0.01** |
| Glycerol | 7.15 ± 2.59 | 1.67 ± 0.26 | 0.10 |
| Glycerol-6-phosphate | 0.35 ± 0.08 | 0.46 ± 0.01 | 0.24 |
| Glycine | 0.05 ± 0.03 | 0.22 ± 0.01 | **0.00** |
| Glycolic acid | 1.77 ± 0.2 | 0.44 ± 0.02 | **0.00** |
| Glycyl-L-tyrosine | 1.81 ± 0.76 | 4.91 ± 0.47 | **0.03** |
| Guanosine | 8.25 ± 3.98 | 4.39 ± 0.36 | 0.39 |
| Heptadecanoic acid | 1.94 ± 0.44 | 4.23 ± 0.26 | **0.01** |
| Hypotaurine | 0.04 ± 0 | 0.05 ± 0 | 0.05 |
| Inosine | 1.25 ± 1 | 1.47 ± 0.21 | 0.84 |
| Isocitric acid | 7.78 ± 2.86 | 2.38 ± 0.1 | 0.13 |
| Isomaltose | 9.71 ± 0.84 | 0.2 ± 0.01 | **0.00** |

| | | | |
|---|---|---|---|
| Itaconic acid | 0.63 ± 0.35 | 2.09 ± 0.27 | **0.03** |
| Lactamide | 0.34 ± 0.17 | 0.62 ± 0.03 | 0.17 |
| Lactic acid | 0.86 ± 0.45 | 0.04 ± 0 | 0.14 |
| Lactitol | 1.38 ± 0.55 | 10.7 ± 0.38 | **0.00** |
| Lactobionic acid | 3.07 ± 0.17 | 0.12 ± 0.01 | **0.00** |
| Lactose | 2.05 ± 1.01 | 1.88 ± 0.05 | 0.88 |
| Lactulose | 0.05 ± 0.01 | 1.12 ± 0.01 | **0.00** |
| L-alanine | 0.6 ± 0.3 | 0.72 ± 0.12 | 0.74 |
| Lanosterol | 0.03 ± 0 | 0.07 ± 0.01 | **0.02** |
| Lauric acid | 1.61 ± 0.18 | 2.65 ± 0.03 | **0.01** |
| L-canavanine | 1.36 ± 0.12 | 0 ± 0 | **0.00** |
| L-cysteine | 5.58 ± 1.99 | 2.62 ± 0.05 | 0.21 |
| L-glutamic acid | 0.18 ± 0.07 | 0.6 ± 0.02 | **0.01** |
| L-glutamine | 0.83 ± 0.51 | 0.12 ± 0.05 | 0.24 |
| L-homoserine | 1.36 ± 0.2 | 1.76 ± 0.04 | 0.12 |
| L-isoleucine | 0.13 ± 0.06 | 5.98 ± 0.09 | **0.00** |
| L-leucine | 1.12 ± 0.5 | 2.82 ± 0.07 | **0.03** |
| L-lysine | 0.58 ± 0.3 | 4.76 ± 0.28 | **0.00** |
| L-methionine | 0.2 ± 0.02 | 4.86 ± 0.22 | **0.00** |
| L-methionine sulfoxide | 0.7 ± 0.04 | 10.93 ± 0.6 | **0.00** |
| L-mimosine | 1.3 ± 0.3 | 1.87 ± 0.1 | 0.15 |
| L-norleucine | 3.68 ± 0.2 | 3.54 ± 0.14 | 0.59 |
| L-proline | 1.02 ± 0.51 | 1.79 ± 0.03 | 0.20 |
| L-pyroglutamic acid | 0.66 ± 0.31 | 0.86 ± 0.03 | 0.55 |
| L-serine | 0.65 ± 0.38 | 0.9 ± 0.06 | 0.55 |
| L-threonine | 0.82 ± 0.1 | 0.46 ± 0.04 | **0.03** |
| L-tryptophan | 1.73 ± 1.1 | 6.49 ± 1.07 | **0.04** |
| L-tyrosine | 0.39 ± 0.2 | 2.54 ± 0.2 | **0.00** |
| L-valine | 0.6 ± 0.2 | 1.3 ± 0.35 | 0.16 |
| Maleamic acid | 2.42 ± 1.21 | 9.38 ± 0.57 | **0.01** |
| Malonic acid | 2.21 ± 0.57 | 0.34 ± 0.02 | **0.03** |
| Maltose | 0.49 ± 0.08 | 2.84 ± 1.65 | 0.23 |
| Maltotriitol | 11.1 ± 2.27 | 0.3 ± 0.03 | **0.01** |
| Maltotriose | 0.49 ± 0.25 | 12.08 ± 0.88 | **0.00** |
| Melezitose | 1.09 ± 0.54 | 9.95 ± 0.32 | **0.00** |
| Melibiose | 1.54 ± 0.42 | 0.76 ± 0.04 | 0.14 |
| Methyl palmitate | 2.25 ± 0.33 | 0.14 ± 0.03 | **0.00** |
| Methyl-beta-D-galactopyranoside | 2 ± 0.42 | 6.02 ± 0.12 | **0.00** |
| Methylmalonic acid | 6.26 ± 2.91 | 0.09 ± 0.08 | 0.10 |
| Myo-inositol | 0.07 ± 0 | 0.41 ± 0.01 | **0.00** |
| Myristic acid | 0.29 ± 0.07 | 0.46 ± 0.06 | 0.14 |
| N-(2-hydroxyethyl)iminodiacetic acid | 0.89 ± 0.1 | 0.42 ± 0.03 | **0.01** |
| N-acetyl-D-glucosamine | 0.11 ± 0.01 | 0.71 ± 0.32 | 0.13 |
| N-acetyl-D-mannosamine | 2.03 ± 0.12 | 3.4 ± 0.38 | **0.03** |
| N-acetyl-L-aspartic acid | 0.93 ± 0.21 | 0.6 ± 0.02 | 0.19 |

| | | | |
|---|---|---|---|
| N-acetyl-L-glutamic acid | 0.29 ± 0.29 | 0.53 ± 0.05 | 0.45 |
| N-acetyl-L-leucine | 2.94 ± 2.61 | 5 ± 0.13 | 0.48 |
| N-carbamyl-L-glutamic acid | 1.25 ± 1.23 | 9 ± 0.62 | **0.00** |
| Neohesperidin | 1.13 ± 0.54 | 2.12 ± 0.08 | 0.15 |
| N-ethylglycine | 0 ± 0 | 12.7 ± 0.16 | **0.00** |
| Nicotinamide | 2.55 ± 0.38 | 2.4 ± 0.04 | 0.71 |
| Nicotinic acid | 0 ± 0 | 11.62 ± 0.16 | **0.00** |
| N-methylglutamic acid | 0.43 ± 0.04 | 1.1 ± 0.03 | **0.00** |
| Norepinephrine | 7.73 ± 3.86 | 3.57 ± 0.33 | 0.34 |
| Norvaline | 0.07 ± 0.03 | 0.89 ± 0.06 | **0.00** |
| Oleic acid | 0.95 ± 0.03 | 1.13 ± 0.11 | 0.21 |
| O-phosphocolamine | 1.73 ± 0.39 | 0.35 ± 0.01 | **0.02** |
| O-phospho-L-threonine | 0.71 ± 0.45 | 0.79 ± 0.02 | 0.86 |
| Orotic acid | 1.1 ± 0.53 | 10.78 ± 0.4 | **0.00** |
| Oxalic acid | 8.73 ± 0.96 | 1.55 ± 0.7 | **0.00** |
| Palatinitol | 0.63 ± 0.31 | 0.46 ± 0.02 | 0.60 |
| Palatinose | 1.54 ± 1.14 | 0.38 ± 0.04 | 0.37 |
| Palmitic acid | 0.31 ± 0.08 | 0.74 ± 0.06 | **0.01** |
| Pantothenic acid | 1.86 ± 0.21 | 1.01 ± 0.05 | **0.02** |
| Phenylalanine | 1.83 ± 0.75 | 3.9 ± 0.17 | 0.06 |
| Phenyl-beta-glucopyranoside | 1.52 ± 0.5 | 0.84 ± 0.03 | 0.24 |
| Phosphoric acid | 0 ± 0 | 0.75 ± 0.7 | 0.35 |
| p-hydroxyphenyllactic acid | 3.04 ± 0.94 | 3.06 ± 0.09 | 0.98 |
| Picolonic acid | 11.84 ± 1.9 | 0.11 ± 0.05 | **0.00** |
| Porphine | 0.01 ± 0 | 0.41 ± 0.04 | **0.00** |
| Purine riboside | 0.69 ± 0.05 | 0.43 ± 0.03 | **0.01** |
| Putrescine | 0.02 ± 0 | 0.82 ± 0.04 | **0.00** |
| Pyridoxine | 0.04 ± 0 | 0.35 ± 0.01 | **0.00** |
| Pyrophosphate | 0.02 ± 0.01 | 0.44 ± 0.12 | **0.02** |
| Raffinose | 1.9 ± 0.95 | 10.84 ± 0.55 | **0.00** |
| Ribitol | 6.82 ± 1.17 | 0.24 ± 0 | **0.00** |
| Ribonic acid-gamma-lactone | 0.86 ± 0.43 | 3.81 ± 0.11 | **0.00** |
| Ribose | 10.49 ± 1.68 | 0.46 ± 0.05 | **0.00** |
| Ribulose-5-phosphate | 1.85 ± 0.53 | 0.13 ± 0.01 | **0.03** |
| Sarcosine | 2.4 ± 0.26 | 0.02 ± 0.02 | **0.00** |
| Sedoheptulose | 0.57 ± 0.12 | 0.32 ± 0.02 | 0.10 |
| Sophorose | 0.59 ± 0.03 | 3.13 ± 0.39 | **0.00** |
| Stearic acid | 0.33 ± 0.14 | 0.87 ± 0.04 | **0.02** |
| Stigmasterol | 0.43 ± 0.11 | 0.64 ± 0.03 | 0.15 |
| Succinic acid | 0.11 ± 0.03 | 0.97 ± 0.23 | **0.02** |
| Sucrose | 0.31 ± 0.18 | 0.46 ± 0.07 | 0.47 |
| Tagatose | 0.05 ± 0.02 | 0.91 ± 0.01 | **0.00** |
| Talose | 0.32 ± 0.27 | 0.03 ± 0 | 0.35 |
| Threose | 3.34 ± 3.03 | 0.28 ± 0.01 | 0.37 |
| Thymine | 0.16 ± 0.15 | 9.62 ± 0.19 | **0.00** |

| | | | |
|---|---|---|---|
| Trans-aconitic acid | 8.62 ± 4.89 | 1.31 ± 0.12 | 0.21 |
| Trehalose-6-phosphate | 1.08 ± 0.6 | 10.45 ± 0.31 | **0.00** |
| Tyrosine | 0.51 ± 0.25 | 10.25 ± 0.67 | **0.00** |
| Uracil | 0 ± 0 | 10.5 ± 0.16 | **0.00** |
| Urea | 0.01 ± 0 | 0 ± 0 | **0.00** |
| Urethane | 0.3 ± 0.1 | 0.49 ± 0.22 | 0.48 |
| Uridine 5-monophosphate | 0.07 ± 0 | 0.79 ± 0.04 | **0.00** |
| **Xanthosine** | 2.07 ± 0.99 | 1.88 ± 0.06 | 0.86 |
| **Xylitol** | 3.14 ± 0.69 | 2.72 ± 0.1 | 0.58 |

The OPLS-DA diagram showing the discrimination between metabolites secreted by NM and PM was presented in **Figure S5A**. The metabolites that varied between the groups were depicted in the heatmap diagram with hierarchical cluster analysis (**Figure S5B**); the top 15 metabolites that contributed to the discrimination were presented in the VIP diagram (**Figure S5C**). Up- and down-regulated metabolites between NM and PM were shown on the Volcano plot (**Figure S5E**) and the data were verified by a t-test (**Figure S5E**).

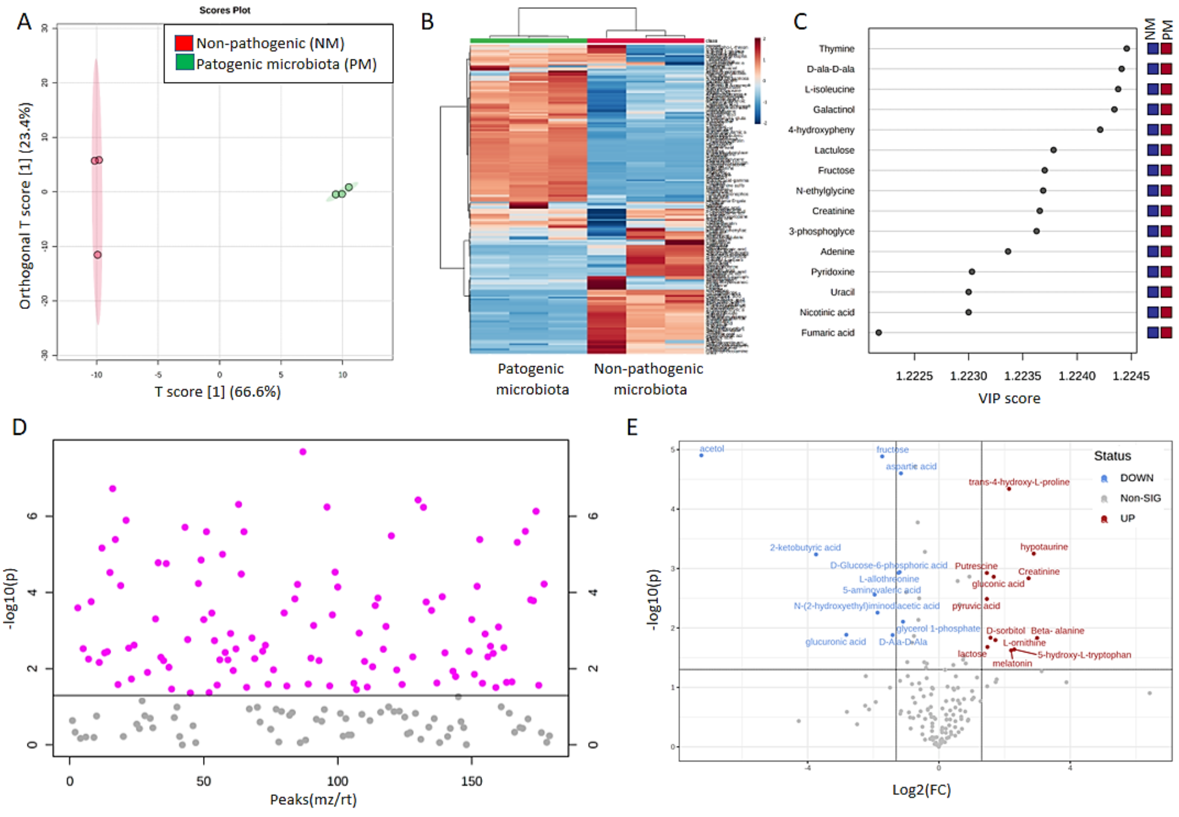

*Figure S5*. Exometabolomics of NM and PM. (A) OPLS-DA diagram showing the discrimination between NM and PM. (B) Heatmap with hierarchical clustering analysis obtained for NM and PM depicted the discrimination in the metabolome between the two groups. (C) VIP graph highlighting the top 15 metabolites that contributed to the metabolomics discrimination. (D) t-test applied for PM vs NM. (E) Volcano plot showing the significant metabolites differentiate in PM compared to NM ($p<0.05$, fold change>2).

**Section 4.** Transcriptome analysis

The growth kinetics of the bacteria needed to be re-evaluated before omics analyses due to the co-culture experiments conducted in the 24-well plates. The growth kinetics of non-pathogenic strains and pathogenic strains were presented in **Figure S6**.

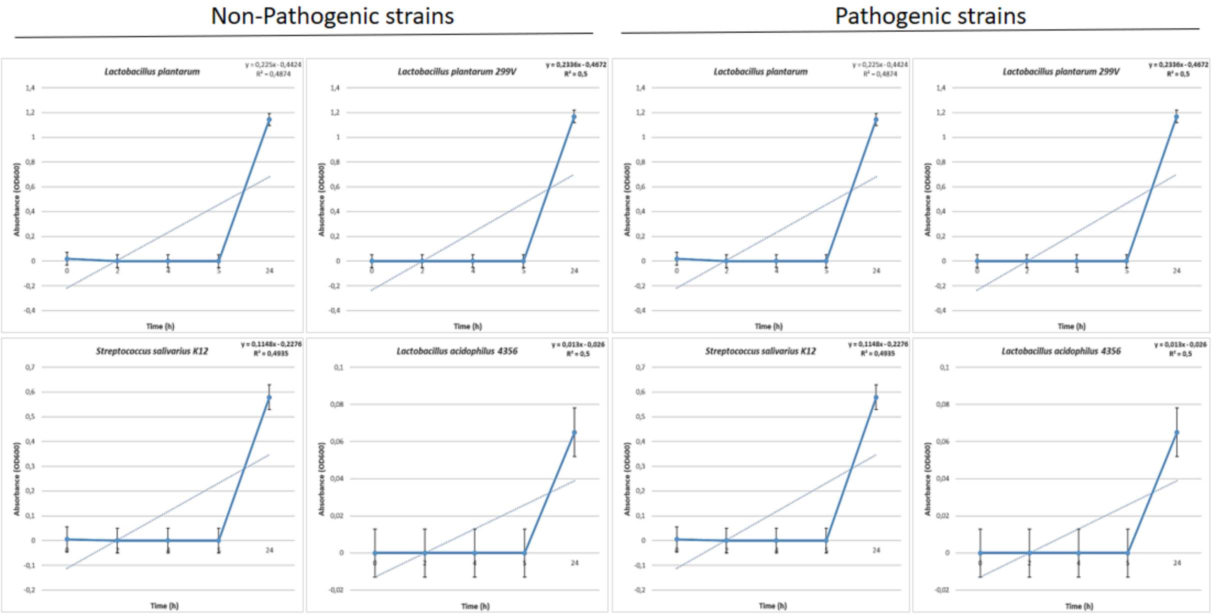

*Figure S6*. Expansion kinetics of non-pathogenic and pathogenic strains in 24-well plate

After COs were co-cultured with NM and PM, we examined the transcriptome data, and determined some key genes which are related to neurodegenerative diseases (**Table S3**).

*Table S3*. Genes which are significantly up-regulated in PM_COs, and are in relation with neurodegenerative diseases

| **Gene Name and p-value** | **Function** | **Reference** |
|---|---|---|
| CDKN1C<br>$p$=4.7x10$^{-185}$ | Cyclin-dependent kinase inhibitor 1C (CDKN1C) is a tight-binding inhibitor of certain G1 cyclin/Cdk complexes. It inhibits cell proliferation. | 12 |

| | | |
|---|---|---|
| Neuronatin (NNat) $p=2.7\times10^{-133}$ | Induces the differentiation of pluripotent stem cells into neural stem cells. At the same time, it has been reported that its overexpression prevents pituitary differentiation. | 13 |
| DAPK3 $p=8.3\times10^{-62}$ | Death-associated protein kinase 3 (DAPK3) plays roles in apoptosis. | 14 |
| BAG3 $p=2.85\times10^{-42}$ | It is effective in age-dependent diseases, such as Alzheimer's. | 15 |
| ODC1 $p=3.65\times10^{-40}$ | It plays a role in Bachmann-Bupp syndrome, which is a neuro-metabolic disorder. It is characterized by symptoms such as macrocephaly. | 16 |
| NOS3 $p=4.68\times10^{-24}$ | Its overexpression is related to neuronal stress and damage. | 17 |
| NRP2 $p=9.25\times10^{-16}$ | It is effective in axon guidance, tumorigenesis, and inflammation. | 18 |
| GAS1 $p=1.13\times10^{-15}$ | It is a pleiotropic protein gene that is effective in cellular arrest and apoptosis. The nervous system is significantly affected by the overexpression of this gene. This is due to its interaction with the signaling pathway inhibited by GDNF. | 19 |
| NRG1 $p=2\times10^{-7}$ | It is known that the interaction between Neuregulin 1 and ErbB4 is effective in the development mechanism of Schizophrenia. Additionally, Neuregulin 1 is known to be associated with anxiety. Endogenous Neuregulin 1 binds to the GABAergic neuronal receptor ErbB4 in the basolateral amygdala. | 20,21 |
| APBA2 $p=4.45\times10^{-6}$ APBA3 $p=4.66\times10^{-4}$ | It encodes a protein which interacts with Alzheimer amyloid precursor protein (APP). | 22 |
| PIDD1 $p=5.17\times10^{-4}$ | The protein encoded by this gene contains a leucine-rich death domain. It interacts with another death domain protein called FADD. It is an adaptor protein in the cell death pathway and is associated with neurotoxicity. | 23 |

The overexpression of the transcripts in PM_COs presented in **Table S3** were validated by RT-qPCR. The list of primers was presented below. The results were presented in **Figure S7**.

**List of primers**

**NNat**

F-CACCCACTTTCGGAACCAT

R-GCAGGGAGTACCTGAACACCT

**BAG3**

F-TGCCAGAAACCACTCAGCCAGA

R-TGAGGATGAGCAGTCAGAGGCA

**NOS3**

F-GAAGGCGACAATCCTGTATGGC

R-TGTTCGAGGGACACCACGTCAT

**NRG1**

F-GATTCCTACCGAGACTCTCCTC

R-TGGAAGGCATGGACACCGTCAT

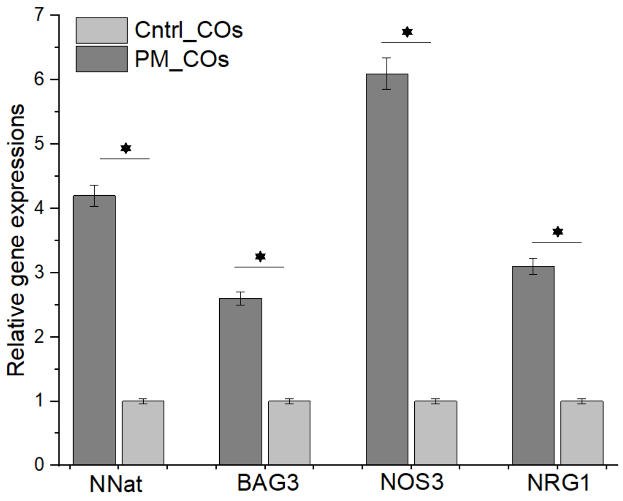

*Figure S7.* Validation of key transcripts by RT-qPCR

When examining the gene expression graph presented in **Figure S7**, it can be observed that there is a 4.2-fold increase in the expression of NNat in PM_CO compared to Cntrl_CO, a 2.5-fold increase in the expression of BAG3, a 6-fold increase in the expression of NOS3, and a 3.1-fold increase in the expression of NRG1.

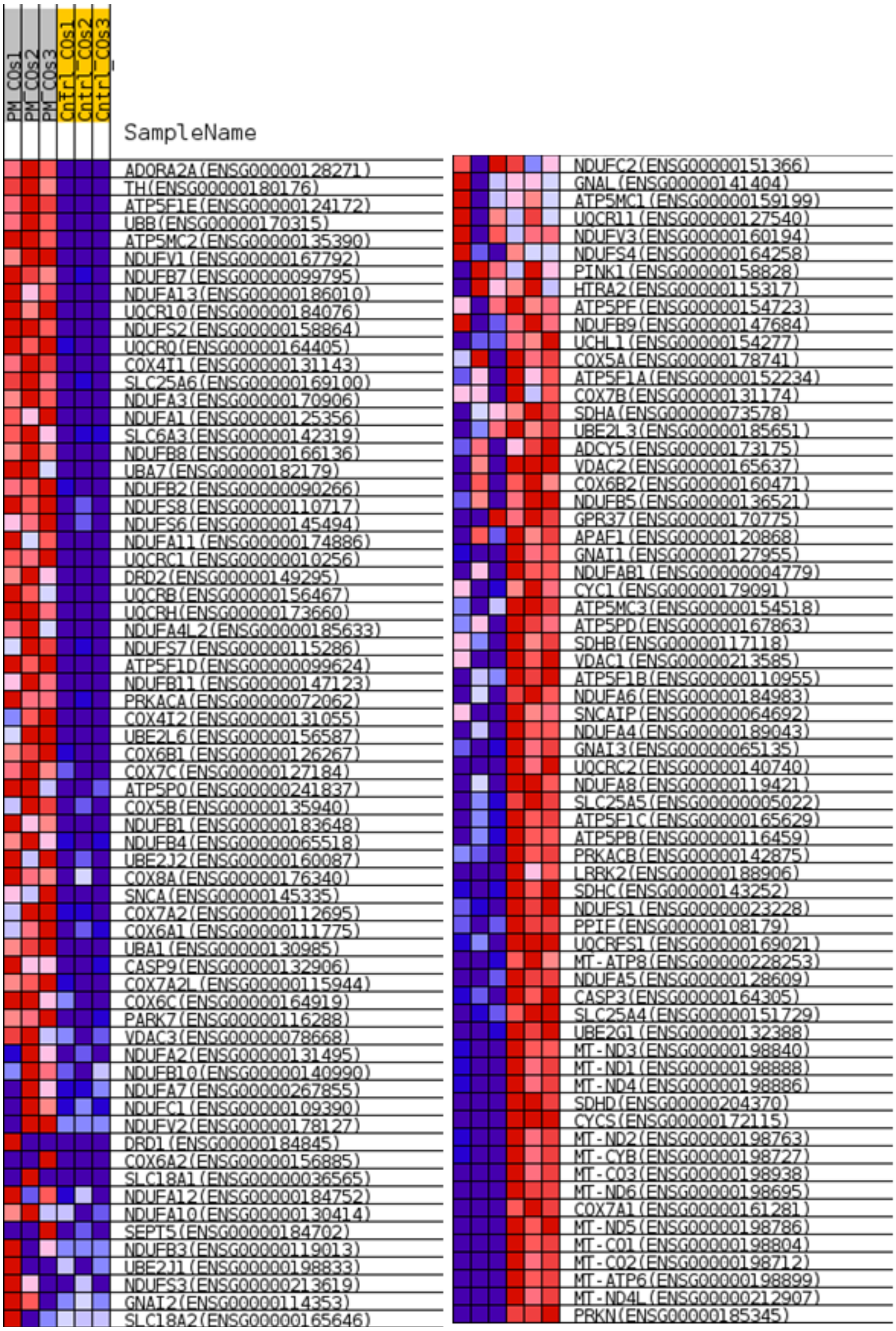

*Figure S8*. Heatmap diagram depicting the statistically significant transcripts in PM_COs compared to Cntrl_COs that are also related to Parkinson's Disease

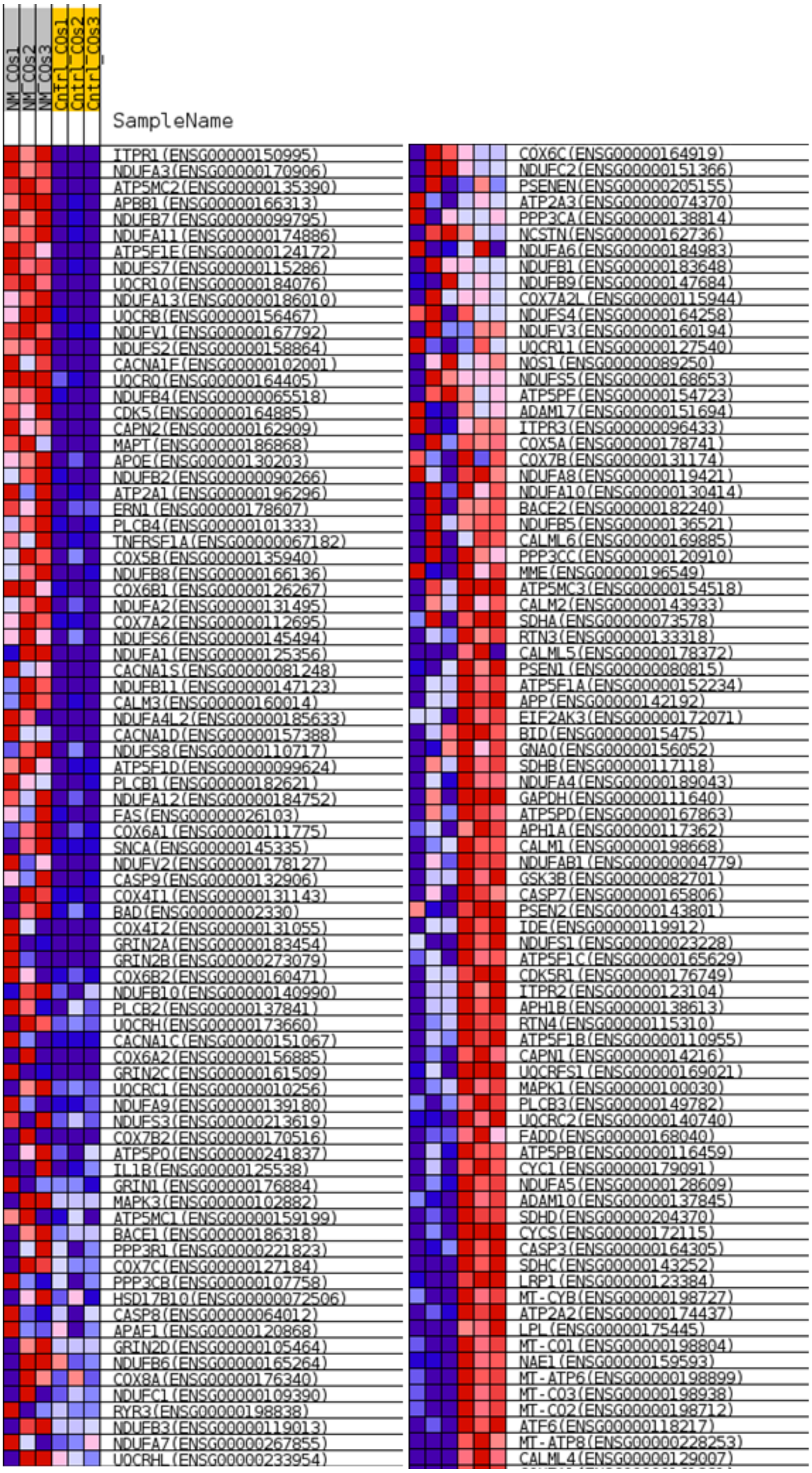

*Figure S9*. Heatmap diagram depicting the statistically significant transcripts in PM_COs compared to Cntrl_COs that are also related to Alzheimer's Disease

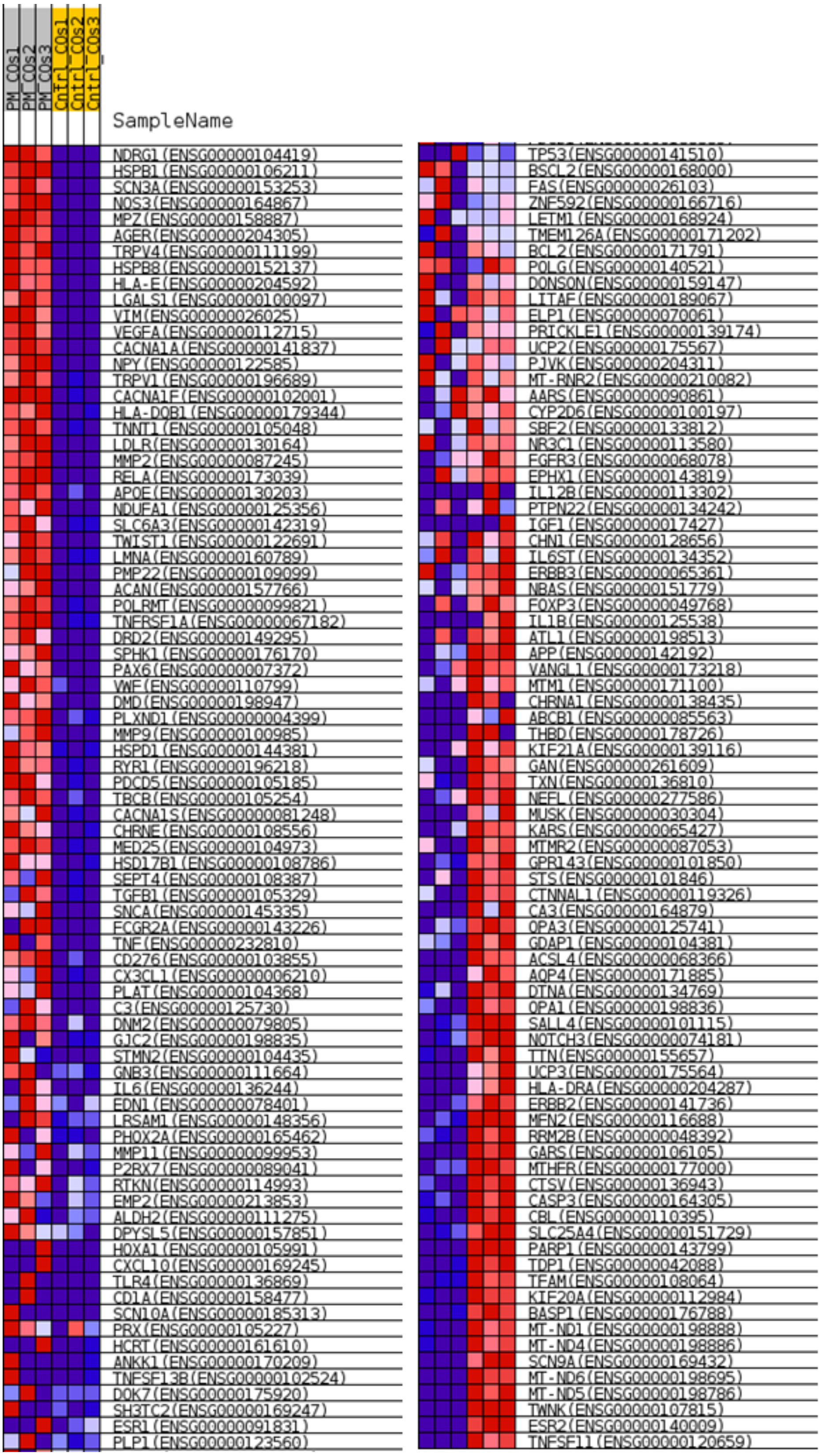

*Figure S10*. Heatmap diagram depicting the statistically significant transcripts in PM_COs compared to Cntrl_COs that are also related to Neuropathy

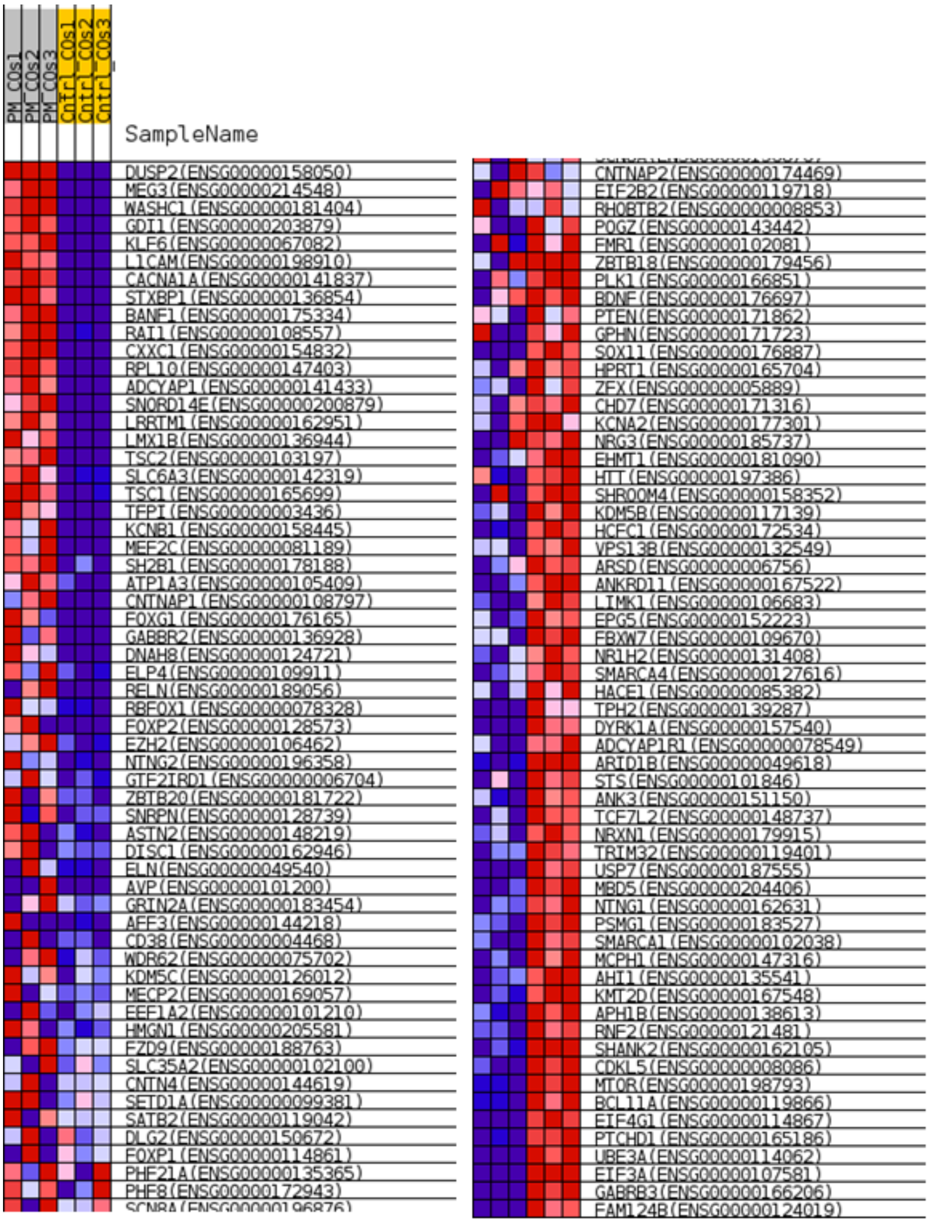

*Figure S11*. Heatmap diagram depicting the statistically significant transcripts in PM_COs compared to Cntrl_COs that are also related to neurodevelopmental diseases

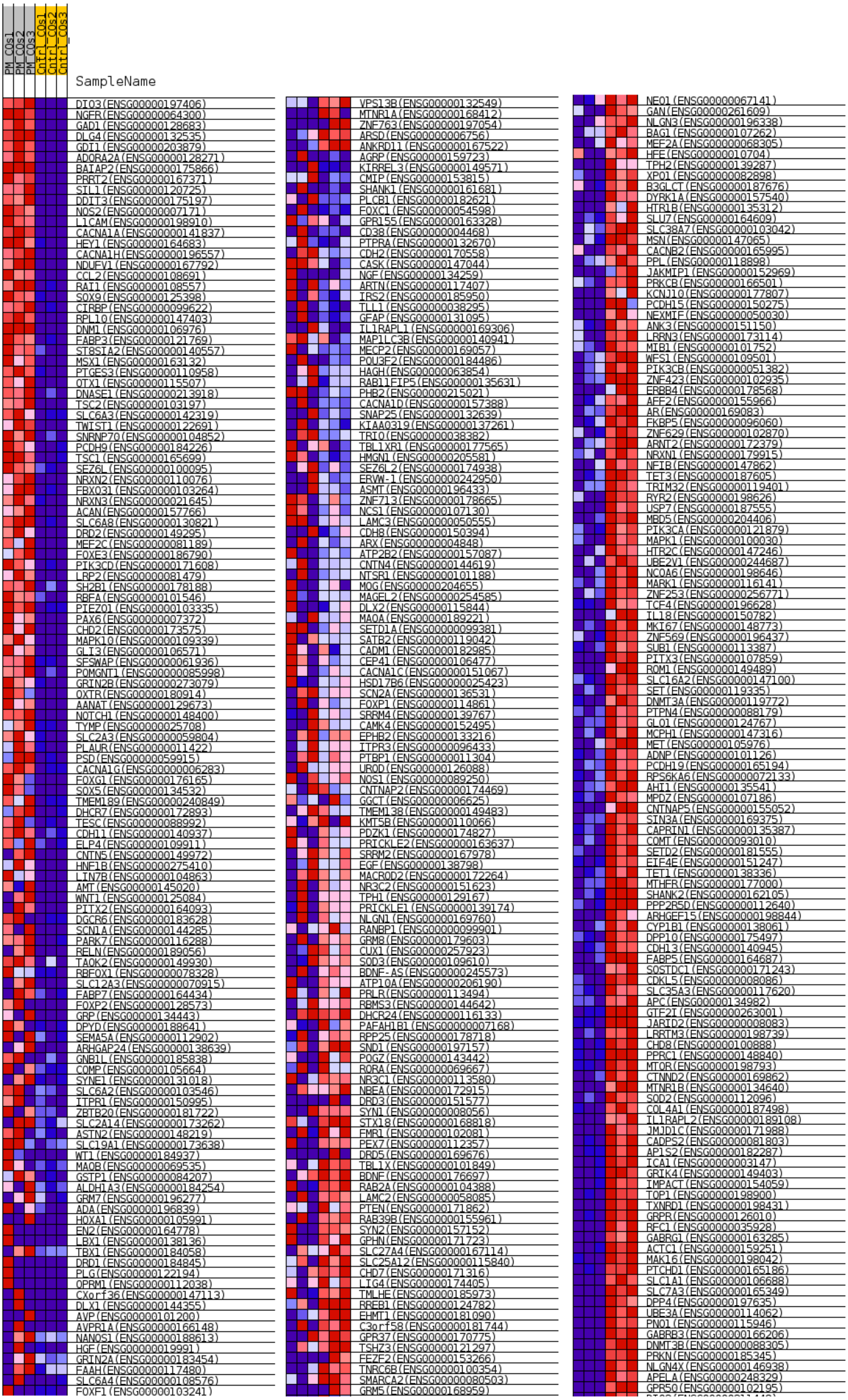

*Figure S12*. Heatmap diagram depicting the statistically significant transcripts in PM_COs compared to Cntrl_COs that are also related to Autism Spectrum Disorder

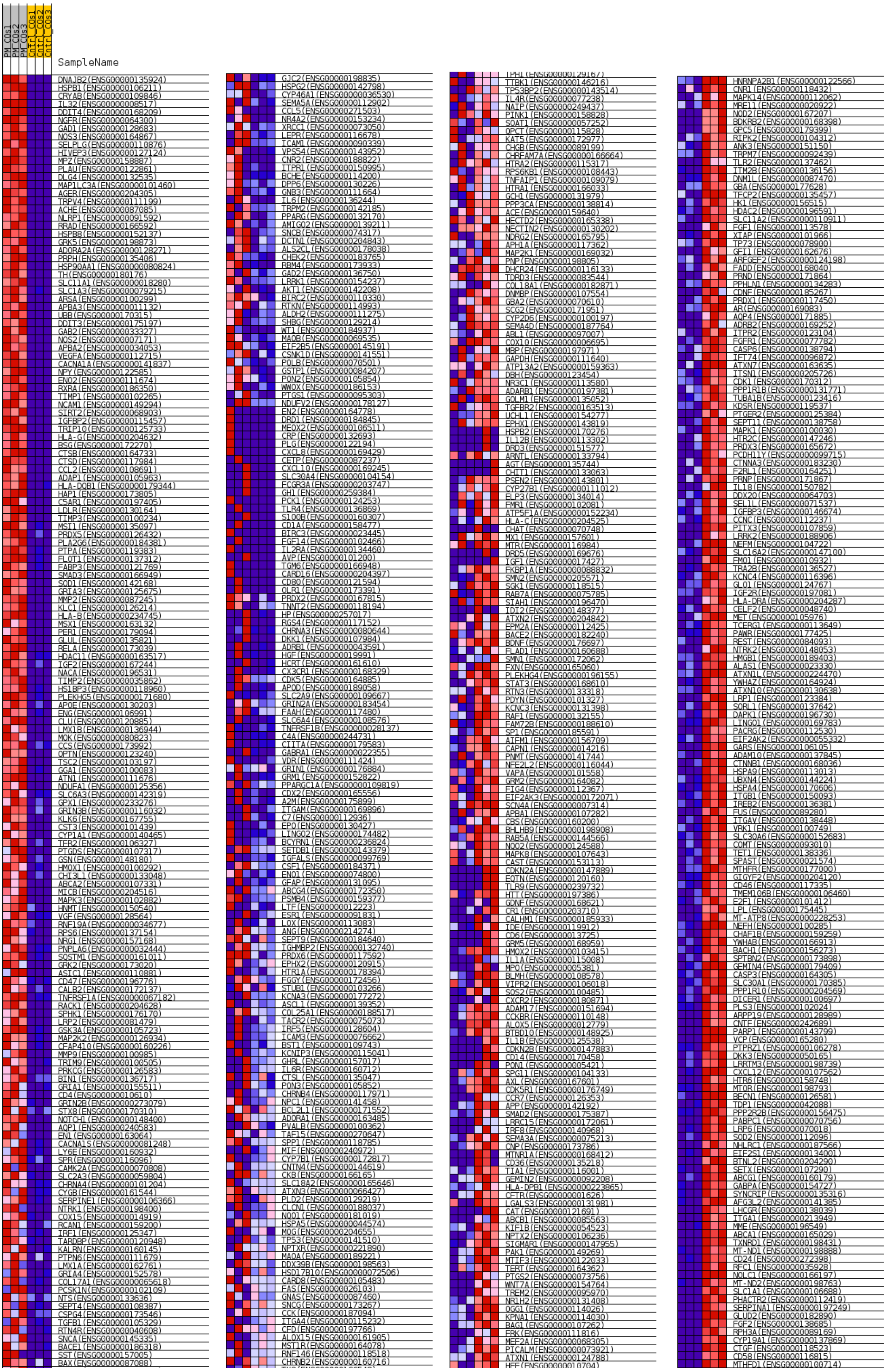

*Figure S13*. Heatmap diagram depicting the statistically significant transcripts in PM_COs compared to Cntrl_COs that are also related to neurodegenerative diseases

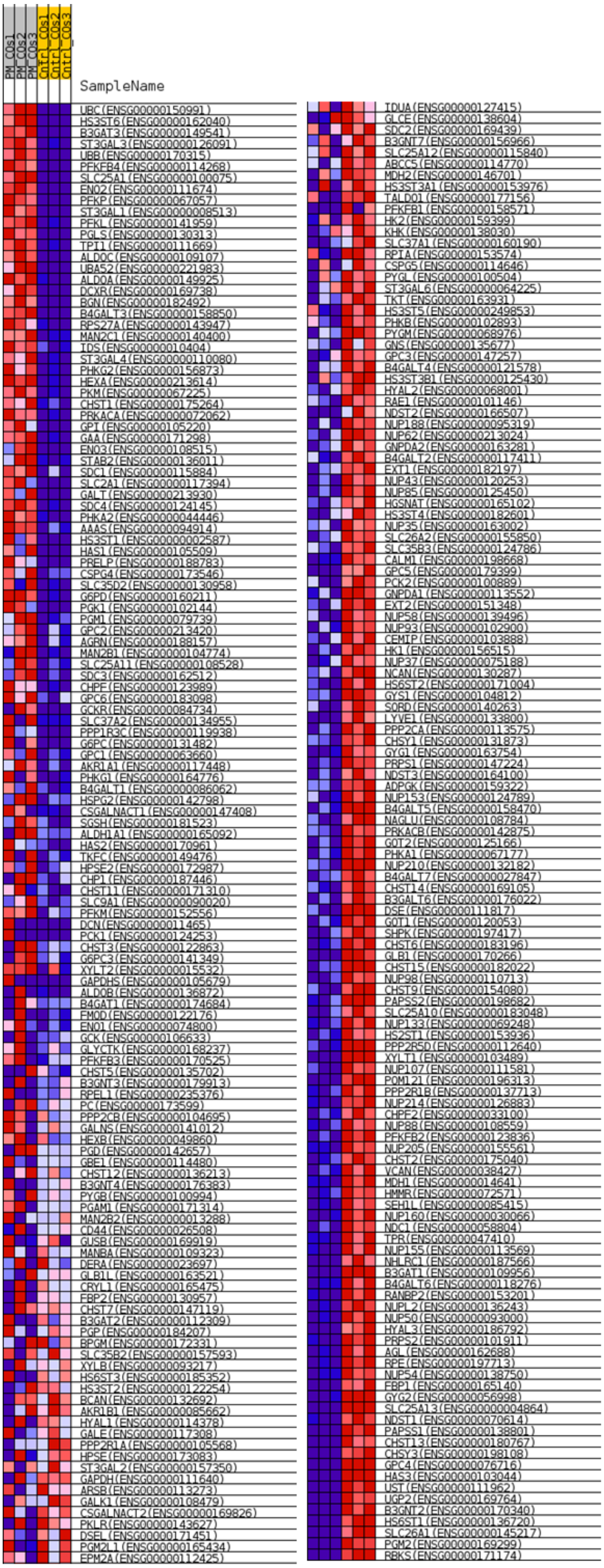

*Figure S14*. Heatmap diagram depicting the statistically significant transcripts in PM_COs compared to Cntrl_COs that are also related to carbohydrate metabolism

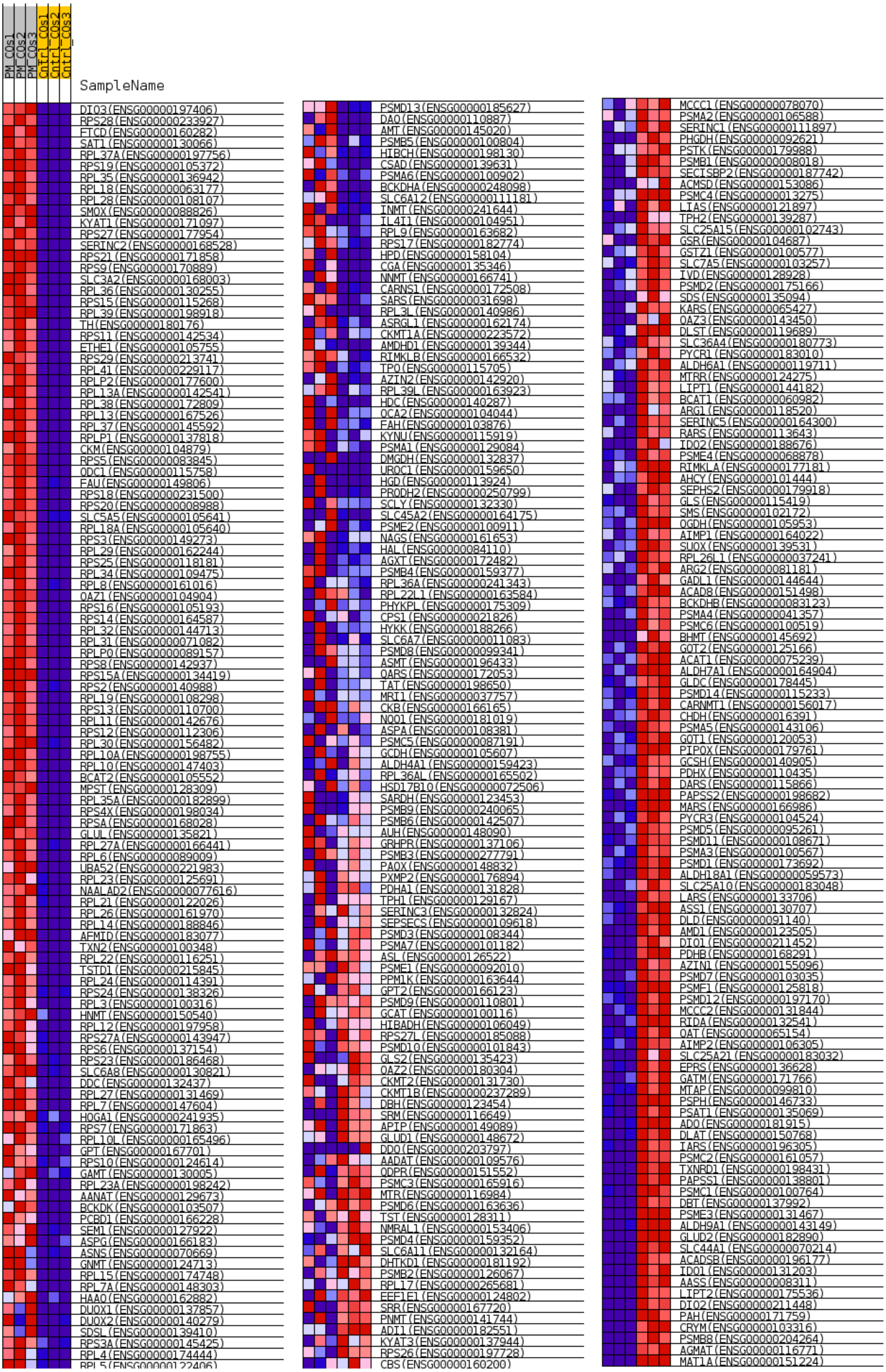

*Figure S15*. Heatmap diagram depicting the statistically significant transcripts in PM_COs compared to Cntrl_COs that are also related to amino acid metabolism

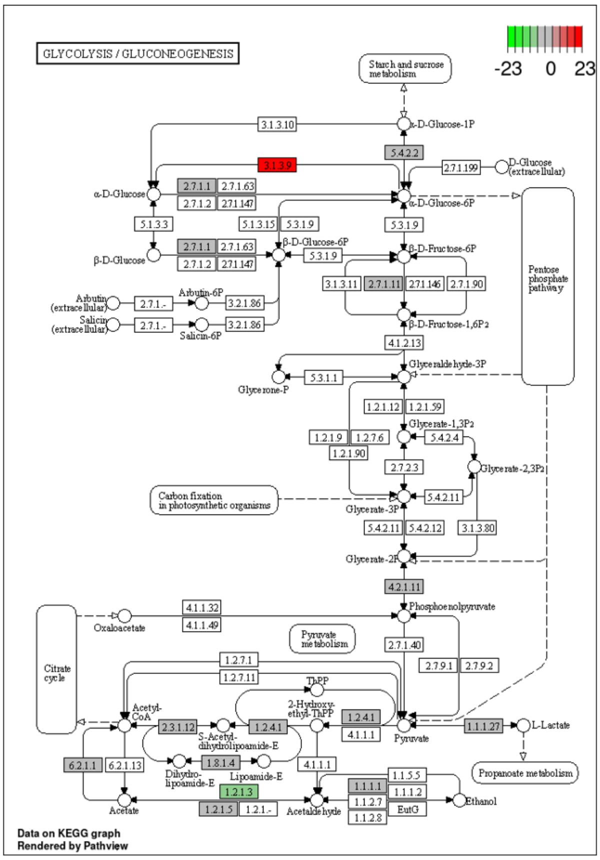

Pathway affected by PM in COs: 1 - Glicolysis/gluconeogenesis

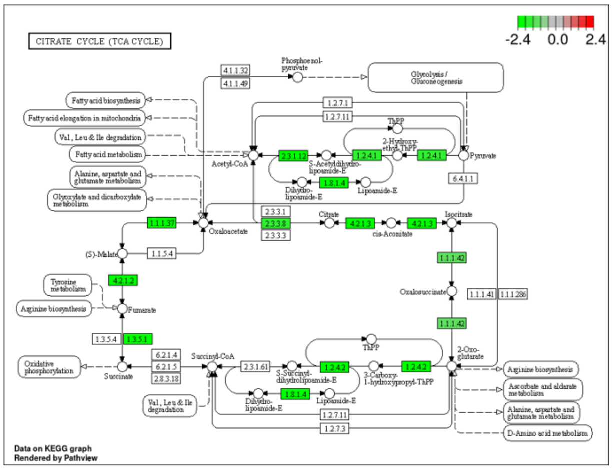

Pathway affected by PM in COs: 2 - TCA

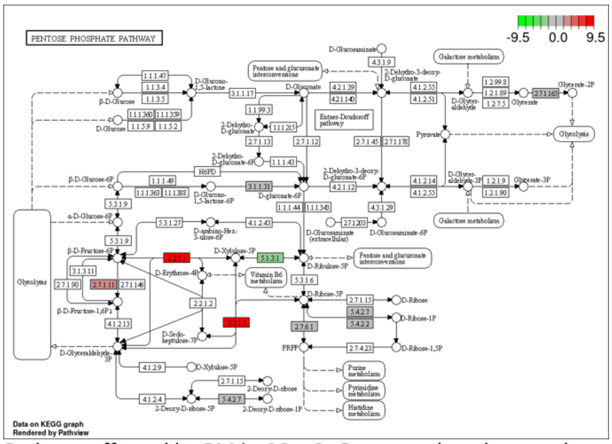

Pathway affected by PM in COs: 3 - Pentose-phosphate pathway

*Figure S16*. Pathways obtained with transcriptome data which were affected by PM in COs

**Section 5**. Proteome analysis

We performed a proteome analysis by LC-qTOF-MS to complement the transcriptome analysis.

The list of proteins, statistical analysis results and their corresponding genes were presented in **Table S4**.

*Table S4*. The list of identified proteins between the groups of PM_COs, NM_COs, and Cntrl_COs

| Proteins | Genes | PM_COs | Cntrl_COs | p | NM_COs | p |
|---|---|---|---|---|---|---|
| | | Mean ± SE | Mean ± SE | | Mean ± SE | |
| Dynein axonemal heavy chain 8 | DNAH8 | 11,78 ± 0,49 | 10,44 ± 0,36 | 0,093 | 11,2 ± 0,28 | 0,362 |
| Endoplasmin | HSP90B1 | 12,88 ± 0,63 | 12,55 ± 0,31 | 0,659 | 12,07 ± 0,28 | 0,304 |
| Rho GTPase-activating protein 23 | ARHGAP23 | 13,66 ± 1,22 | 12,61 ± 1,62 | 0,633 | 12,58 ± 1,86 | 0,654 |
| Putative elongation factor 1-alpha-like 3 | EEF1A1P5 | 14,23 ± 0,33 | 12,81 ± 0,92 | 0,222 | 13,78 ± 0,03 | 0,242 |

| | | | | | | |
|---|---|---|---|---|---|---|
| Ras-related protein Rab-13 | RAB13 | 11,84 ± 0,4 | 11,01 ± 0,48 | 0,253 | 11,38 ± 0,23 | 0,371 |
| Unconventional myosin-XV | MYO15A | 11,29 ± 0,34 | 10,97 ± 0,53 | 0,643 | 11,14 ± 0,33 | 0,773 |
| Diacylglycerol kinase eta | DGKH | 11,58 ± 0,56 | 10,31 ± 0,51 | 0,168 | 10,5 ± 0,21 | 0,143 |
| Centrosomal protein of 83 kDa | CEP83 | 13,57 ± 0,15 | 12,42 ± 1,13 | 0,373 | 11,56 ± 1,09 | 0,141 |
| Pecanex-like protein 2 | PCNXL2 | 13,45 ± 1,66 | 10,91 ± 0,51 | 0,218 | 11,03 ± 0,24 | 0,222 |
| Transient receptor potential cation channel subfamily M member 8 | TRPM8 | 13,69 ± 0,15 | 13,79 ± 0,19 | 0,702 | 13,57 ± 0,15 | 0,603 |
| E3 ubiquitin-protein ligase UBR4 | UBR4 | 11,59 ± 0,25 | 10,22 ± 0,35 | **0,033** | 11,14 ± 0,46 | 0,440 |
| Tubulin beta-3 chain | TUBB3 | 12,3 ± 0,38 | 10,95 ± 0,7 | 0,165 | 11,34 ± 0,48 | 0,192 |
| Zinc finger protein 280B | ZNF280B | 12,01 ± 0,31 | 11,62 ± 0,44 | 0,518 | 12,56 ± 0,7 | 0,515 |
| 40S ribosomal protein S18 | RPS18 | 12,31 ± 0,46 | 11,7 ± 0,3 | 0,332 | 11,14 ± 0,44 | 0,138 |
| Cathepsin D | CTSD | 11,63 ± 0,36 | 12,25 ± 0,21 | 0,209 | 11,13 ± 0,33 | 0,366 |
| Cation channel sperm-associated protein subunit beta | CATSPERB | 14,58 ± 0,11 | 12,88 ± 1,64 | 0,360 | 13,02 ± 1,49 | 0,356 |
| Alpha-enolase | ENO1 | 14,14 ± 0,44 | 14,38 ± 0,09 | 0,614 | 14,51 ± 0,1 | 0,454 |
| General transcription and DNA repair factor IIH helicase subunit XPB | ERCC3 | 11,89 ± 0,34 | 10,59 ± 0,52 | 0,105 | 11,5 ± 0,8 | 0,671 |
| Elongation factor 1-alpha 2 | EEF1A2 | 11,97 ± 0,65 | 12,68 ± 0,27 | 0,366 | 11,89 ± 0,51 | 0,928 |
| Rho guanine nucleotide exchange factor 2 | ARHGEF2 | 11,83 ± 0,43 | 10,49 ± 0,24 | 0,055 | 10,48 ± 0,44 | 0,095 |
| Ribosomal protein S6 kinase-related protein | RSKR | 11,51 ± 0,52 | 10,19 ± 0,25 | 0,085 | 10,84 ± 0,43 | 0,376 |
| Myosin-7B | MYH7B | 12,35 ± 0,59 | 11,83 ± 0,99 | 0,677 | 11,82 ± 0,6 | 0,563 |
| Centrosome and spindle pole-associated protein 1 | CSPP1 | 12,1 ± 0,28 | 10,65 ± 0,46 | 0,054 | 11,91 ± 1,14 | 0,882 |
| Prelamin-A/C | LMNA | 12,2 ± 0,47 | 10,54 ± 0,27 | **0,037** | 11,62 ± 0,59 | 0,486 |
| NADH dehydrogenase [ubiquinone] iron-sulfur protein 2, mitochondrial | NDUFS2 | 12,04 ± 0,11 | 10,57 ± 0,43 | **0,031** | 11,19 ± 0,31 | 0,064 |
| 60 kDa heat shock protein | HSPD1 | 12,92 ± 0,21 | 12,91 ± 0,39 | 0,986 | 12,61 ± 0,42 | 0,542 |
| Nucleophosmin | NPM1 | 13,2 ± 0,28 | 12,48 ± 0,87 | 0,473 | 13,26 ± 0,17 | 0,877 |
| Transient receptor potential cation channel subfamily M member 3 | TRPM3 | 11,81 ± 0,29 | 10,77 ± 0,53 | 0,159 | 11,4 ± 0,55 | 0,547 |
| 1-phosphatidylinositol 4,5-bisphosphate phosphodiesterase beta-4 | PLCB4 | 11,58 ± 0,34 | 10,62 ± 0,46 | 0,171 | 10,38 ± 0,53 | 0,130 |
| Calnexin | CANX | 12,13 ± 0,2 | 11,41 ± 0,7 | 0,377 | 11,76 ± 0,51 | 0,537 |
| Protein disulfide-isomerase | P4HB | 12,98 ± 0,32 | 13,54 ± 0,19 | 0,211 | 13,24 ± 0,43 | 0,651 |
| Endoplasmic reticulum chaperone BiP | HSPA5 | 13,96 ± 0,37 | 14,04 ± 0,22 | 0,852 | 13,92 ± 0,36 | 0,939 |
| Vimentin | VIM | 13,5 ± 0,39 | 13,2 ± 0,22 | 0,541 | 14,35 ± 0,42 | 0,207 |

| | | | | | | |
|---|---|---|---|---|---|---|
| Heterogeneous nuclear ribonucleoproteins C1/C2 | HNRNPC | 13,27 ± 0,32 | 12,94 ± 0,2 | 0,423 | 12,65 ± 0,36 | 0,260 |
| Upstream stimulatory factor 2 | USF2 | 15,04 ± 0,29 | 10,71 ± 0,3 | **0,001** | 12,55 ± 1,37 | 0,150 |
| Pyruvate kinase PKM | PKM | 13,68 ± 0,33 | 13,58 ± 0,36 | 0,848 | 13,37 ± 0,2 | 0,464 |
| Polyubiquitin-C | UBC | 13,81 ± 0,33 | 13,9 ± 0,26 | 0,838 | 13,56 ± 0,37 | 0,634 |
| Synaptopodin-2 | SYNPO2 | 11,99 ± 0,38 | 11,89 ± 0,44 | 0,872 | 12,41 ± 0,78 | 0,656 |
| Ubiquitin carboxyl-terminal hydrolase 28 | USP28 | 11,67 ± 0,26 | 10,77 ± 0,17 | **0,045** | 11,35 ± 0,34 | 0,504 |
| 40S ribosomal protein S7 | RPS7 | 11,79 ± 0,56 | 10,98 ± 0,43 | 0,311 | 11,01 ± 0,33 | 0,295 |
| Doublecortin domain-containing protein 1 | DCDC1 | 11,85 ± 0,47 | 10,76 ± 0,07 | 0,086 | 11,8 ± 0,64 | 0,958 |
| Prothymosin alpha | PTMA | 11,65 ± 0,17 | 10,86 ± 0,3 | 0,082 | 10,6 ± 0,52 | 0,128 |
| 60S ribosomal protein L23 | RPL23 | 11,47 ± 0,3 | 11,25 ± 0,26 | 0,615 | 10,86 ± 0,45 | 0,322 |
| WD repeat-containing protein 81 | WDR81 | 11,6 ± 0,33 | 11,27 ± 0,33 | 0,507 | 11,12 ± 0,43 | 0,424 |
| Prohibitin | PHB1 | 11,71 ± 0,27 | 11,69 ± 0,23 | 0,955 | 10,84 ± 0,33 | 0,109 |
| SRSF protein kinase 1 | SRPK1 | 12,7 ± 0,68 | 11,8 ± 0,89 | 0,465 | 10,88 ± 0,48 | 0,094 |
| Chitotriosidase-1 | CHIT1 | 12,35 ± 0,21 | 10,63 ± 0,09 | **0,002** | 10,98 ± 0,34 | **0,026** |
| Multicilin | MCIDAS | 11,61 ± 0,38 | 11,16 ± 0,52 | 0,524 | 11,23 ± 1,03 | 0,745 |
| Membrane-associated transporter protein | SLC45A2 | 12,13 ± 0,33 | 10,84 ± 0,76 | 0,196 | 11,22 ± 0,42 | 0,161 |
| 39S ribosomal protein L15, mitochondrial | MRPL15 | 11,86 ± 0,78 | 10,64 ± 0,26 | 0,214 | 11,82 ± 0,7 | 0,975 |
| Eukaryotic initiation factor 4A-I | EIF4A1 | 11,94 ± 0,47 | 11,51 ± 0,99 | 0,718 | 12,17 ± 0,17 | 0,663 |
| 60S ribosomal protein L14 | RPL14 | 12,06 ± 0,44 | 11,19 ± 0,22 | 0,153 | 10,73 ± 0,28 | 0,065 |
| Cyclin-dependent kinase inhibitor 1B | CDKN1B | 11,49 ± 0,31 | 11,23 ± 0,17 | 0,499 | 11,29 ± 0,65 | 0,793 |
| Glyceraldehyde-3-phosphate dehydrogenase | GAPDH | 16,02 ± 0,26 | 16,03 ± 0,31 | 0,998 | 15,79 ± 0,05 | 0,426 |
| Dihydropyrimidinase-related protein 5 | DPYSL5 | 12,59 ± 0,49 | 13,25 ± 0,95 | 0,571 | 12,43 ± 0,38 | 0,815 |
| Myb-related protein A | MYBL1 | 11,94 ± 0,2 | 10,96 ± 0,3 | 0,054 | 10,87 ± 0,64 | 0,187 |
| Putative E3 ubiquitin-protein ligase UBR7 | UBR7 | 11,98 ± 0,3 | 11,03 ± 0,31 | 0,092 | 10,76 ± 0,35 | 0,056 |
| Heat shock cognate 71 kDa protein | HSPA8 | 12,78 ± 0,1 | 13,16 ± 0,24 | 0,211 | 12,73 ± 0,08 | 0,740 |
| Exonuclease mut-7 homolog | EXD3 | 11,9 ± 0,38 | 10,96 ± 0,36 | 0,150 | 12,16 ± 0,96 | 0,812 |
| Zinc transporter ZIP10 | SLC39A10 | 11,72 ± 0,44 | 10,74 ± 0,64 | 0,275 | 11,09 ± 0,48 | 0,388 |
| Phosphatidylinositol 4-phosphate 3-kinase C2 domain-containing subunit beta | PIK3C2B | 11,9 ± 0,46 | 10,5 ± 0,21 | 0,050 | 10,8 ± 0,26 | 0,106 |
| Tubulin alpha-1C chain | TUBA1C | 11,68 ± 0,64 | 11,1 ± 0,28 | 0,452 | 10,87 ± 0,35 | 0,326 |

| | | | | | | |
|---|---|---|---|---|---|---|
| Neutral alpha-glucosidase AB | GANAB | 12,39 ± 0,45 | 11,77 ± 0,61 | 0,454 | 11,66 ± 0,68 | 0,420 |
| Transcription termination factor 2 | TTF2 | 12,05 ± 0,08 | 10,49 ± 0,01 | **0,000** | 11,36 ± 0,16 | **0,017** |
| 39S ribosomal protein L42, mitochondrial | MRPL42 | 11,8 ± 0,5 | 10,93 ± 0,14 | 0,169 | 12,01 ± 0,23 | 0,714 |
| Galectin-1 | LGALS1 | 13,22 ± 0,2 | 12,71 ± 0,21 | 0,160 | 12,52 ± 0,96 | 0,513 |
| Ribosomal protein S6 kinase beta-2 | RPS6KB2 | 12,21 ± 0,6 | 11,29 ± 0,9 | 0,447 | 11,04 ± 0,43 | 0,193 |
| Nebulin | NEB | 11,91 ± 0,3 | 12,5 ± 0,32 | 0,250 | 11,21 ± 0,44 | 0,263 |
| NACHT, LRR and PYD domains-containing protein 2 | NLRP2 | 11,98 ± 0,43 | 10,51 ± 0,36 | 0,060 | 11,28 ± 0,11 | 0,190 |
| DCC-interacting protein 13-beta | APPL2 | 12,03 ± 0,26 | 11,17 ± 0,04 | **0,030** | 10,42 ± 0,3 | **0,015** |
| Regulator of microtubule dynamics protein 3 | RMDN3 | 11,89 ± 0,22 | 10,28 ± 0,1 | **0,003** | 10,42 ± 0,14 | **0,005** |
| 60S ribosomal protein L4 | RPL4 | 12,38 ± 0,48 | 11,4 ± 0,28 | 0,157 | 11,56 ± 0,28 | 0,217 |
| 40S ribosomal protein S17 | RPS17 | 11,72 ± 0,28 | 11,25 ± 0,28 | 0,290 | 11,05 ± 0,26 | 0,148 |
| Fructose-bisphosphate aldolase A | ALDOA | 12,47 ± 0,29 | 12,3 ± 0,11 | 0,600 | 11,97 ± 0,17 | 0,201 |
| GRAM domain-containing protein 2A | GRAMD2 | 11,84 ± 0,16 | 10,69 ± 0,49 | 0,088 | 11,71 ± 1,15 | 0,912 |
| Ryanodine receptor 2 | RYR2 | 11,92 ± 0,39 | 13,12 ± 0,12 | **0,044** | 11,48 ± 0,69 | 0,610 |
| Complement component 1 Q subcomponent-binding protein, mitochondrial | C1QBP | 12,17 ± 0,35 | 11,86 ± 0,26 | 0,522 | 11,92 ± 0,38 | 0,663 |
| Uncharacterized protein C19orf47 | C19orf47 | 11,56 ± 0,36 | 10,71 ± 0,17 | 0,099 | 11,82 ± 0,91 | 0,797 |
| Histone H3.3 | H3-3A | 11,52 ± 0,31 | 12,81 ± 0,28 | **0,036** | 11,69 ± 0,69 | 0,836 |
| 40S ribosomal protein S5 | RPS5 | 11,55 ± 0,41 | 11,54 ± 0,18 | 0,979 | 11,1 ± 0,24 | 0,405 |
| 40S ribosomal protein S11 | RPS11 | 12,44 ± 0,09 | 11,46 ± 0,14 | **0,004** | 10,91 ± 0,6 | 0,067 |
| ADAM DEC1 | ADAMDEC1 | 11,69 ± 0,46 | 10,7 ± 0,23 | 0,124 | 12,26 ± 1,15 | 0,664 |
| Histone H2B type 1-N | HIST1H2BN | 16,16 ± 0,18 | 16,34 ± 0,09 | 0,401 | 16,06 ± 0,28 | 0,786 |
| L-lactate dehydrogenase A chain | LDHA | 14,88 ± 0,37 | 13,85 ± 0,8 | 0,307 | 14,19 ± 0,49 | 0,322 |
| Phosphoglycerate kinase 1 | PGK1 | 14,28 ± 0,06 | 13,44 ± 0,46 | 0,144 | 13,67 ± 0,28 | 0,098 |
| Annexin A1 | ANXA1 | 11,9 ± 0,24 | 10,94 ± 0,2 | **0,037** | 10,65 ± 0,55 | 0,106 |
| Superoxide dismutase [Mn], mitochondrial | SOD2 | 13,4 ± 0,44 | 14,34 ± 0,23 | 0,134 | 14,02 ± 0,31 | 0,321 |
| Tubulin beta-4A chain | TUBB4A | 13,76 ± 0,37 | 13,82 ± 0,12 | 0,880 | 11,25 ± 1,31 | 0,140 |
| ADP/ATP translocase 2 | SLC25A5 | 11,88 ± 0,65 | 11,05 ± 0,36 | 0,324 | 11,06 ± 0,34 | 0,325 |
| ATP synthase subunit beta, mitochondrial | ATP5F1B | 13,34 ± 0,32 | 13,47 ± 0,23 | 0,772 | 12,86 ± 0,51 | 0,463 |
| Histone H1.0 | H1-0 | 11,34 ± 0,32 | 10,85 ± 0,22 | 0,282 | 11,9 ± 0,36 | 0,312 |
| Annexin A2 | ANXA2 | 14,38 ± 0,2 | 14,71 ± 0,13 | 0,241 | 14,34 ± 0,12 | 0,880 |
| Tubulin beta chain | TUBB | 14,68 ± 0,67 | 13,14 ± 0,95 | 0,255 | 13,2 ± 1,46 | 0,409 |

| | | | | | | |
|---|---|---|---|---|---|---|
| Profilin-1 | PFN1 | 13,32 ± 0,43 | 13,76 ± 0,36 | 0,473 | 13,2 ± 0,73 | 0,889 |
| Annexin A5 | ANXA5 | 12,96 ± 0,26 | 12,05 ± 0,34 | 0,101 | 12,35 ± 0,41 | 0,280 |
| Annexin A4 | ANXA4 | 11,89 ± 0,08 | 12,32 ± 0,4 | 0,358 | 11,39 ± 0,7 | 0,517 |
| Histone H2A type 1-J | HIST1H2AJ | 16,69 ± 0,61 | 16,8 ± 0,32 | 0,877 | 16,27 ± 0,47 | 0,618 |
| Mast/stem cell growth factor receptor Kit | KIT | 11,86 ± 0,33 | 10,43 ± 0,49 | 0,072 | 11,83 ± 0,6 | 0,967 |
| Elongation factor 2 | EEF2 | 12,29 ± 0,43 | 11,33 ± 0,45 | 0,202 | 10,6 ± 0,39 | **0,044** |
| Macrophage migration inhibitory factor | MIF | 13,28 ± 0,18 | 13,16 ± 0,52 | 0,842 | 12,87 ± 0,42 | 0,413 |
| Histone H2AX | H2AFX | 12,6 ± 0,89 | 13,66 ± 0,51 | 0,362 | 12,65 ± 1,13 | 0,975 |
| Histone H1.5 | HIST1H1B | 11,61 ± 0,44 | 11,63 ± 0,13 | 0,969 | 11,81 ± 0,21 | 0,714 |
| Histone H1.3 | HIST1H1D | 13,72 ± 0,33 | 13,78 ± 0,13 | 0,869 | 13,44 ± 0,27 | 0,539 |
| Phosphoglycerate mutase 1 | PGAM1 | 12,34 ± 0,61 | 12,73 ± 0,37 | 0,616 | 11,55 ± 0,41 | 0,350 |
| Ubiquitin-like modifier-activating enzyme 1 | UBA1 | 11,68 ± 0,35 | 10,95 ± 0,34 | 0,204 | 10,7 ± 0,4 | 0,138 |
| Receptor-type tyrosine-protein phosphatase epsilon | PTPRE | 11,77 ± 0,76 | 10,69 ± 0,12 | 0,230 | 10,66 ± 0,26 | 0,237 |
| Cofilin-1 | CFL1 | 12,85 ± 0,22 | 13,27 ± 0,19 | 0,228 | 12,98 ± 0,12 | 0,639 |
| ATP synthase subunit alpha, mitochondrial | ATP5F1A | 11,97 ± 0,21 | 12 ± 0,81 | 0,972 | 11,28 ± 0,96 | 0,523 |
| Protein disulfide-isomerase A3 | PDIA3 | 12,9 ± 0,27 | 13,26 ± 0,33 | 0,451 | 12,85 ± 0,29 | 0,899 |
| Malate dehydrogenase, mitochondrial | MDH2 | 12,02 ± 0,45 | 12,34 ± 0,33 | 0,588 | 11,61 ± 0,52 | 0,581 |
| Antigen KI-67 | MKI67 | 11,98 ± 0,39 | 10,91 ± 0,4 | 0,129 | 10,73 ± 0,27 | 0,057 |
| Elongation factor Tu, mitochondrial | TUFM | 12,36 ± 0,63 | 10,28 ± 0,3 | **0,040** | 12,63 ± 0,4 | 0,744 |
| Receptor-interacting serine/threonine-protein kinase 4 | RIPK4 | 11,68 ± 0,48 | 10,65 ± 0,19 | 0,116 | 10,74 ± 0,62 | 0,297 |
| Triosephosphate isomerase | TPI1 | 12,35 ± 0,1 | 12,53 ± 0,06 | 0,202 | 12,4 ± 0,17 | 0,816 |
| Actin, cytoplasmic 1 | ACTB | 16,25 ± 0,24 | 16,51 ± 0,05 | 0,353 | 16,11 ± 0,23 | 0,714 |
| Heterogeneous nuclear ribonucleoprotein K | HNRNPK | 12,26 ± 0,26 | 11,46 ± 0,1 | **0,045** | 11,2 ± 0,27 | **0,048** |
| Histone H4 | HIST1H4A | 16,04 ± 0,3 | 15,9 ± 0,44 | 0,801 | 15,84 ± 0,24 | 0,623 |
| Actin, cytoplasmic 2 | ACTG1 | 14,91 ± 0,12 | 14,29 ± 0,48 | 0,278 | 13,75 ± 0,44 | 0,066 |
| Tubulin alpha-1B chain | TUBA1B | 14,76 ± 0,15 | 14,57 ± 0,23 | 0,522 | 14,49 ± 0,23 | 0,373 |
| Tubulin beta-4B chain | TUBB4B | 14,92 ± 0,26 | 14,9 ± 0,44 | 0,979 | 14,73 ± 0,33 | 0,685 |
| Histone H3.1 | HIST1H3A | 15,56 ± 0,29 | 15,47 ± 0,16 | 0,796 | 15,59 ± 0,33 | 0,962 |
| Tubulin beta-2B chain | TUBB2B | 11,93 ± 0,46 | 12 ± 0,5 | 0,922 | 11,65 ± 0,29 | 0,632 |
| Phosphatidate phosphatase LPIN1 | LPIN1 | 11,32 ± 0,35 | 10,78 ± 0,18 | 0,245 | 10,86 ± 0,35 | 0,401 |
| Protein disulfide-isomerase A6 | PDIA6 | 12,3 ± 0,36 | 12,36 ± 0,36 | 0,922 | 12,55 ± 0,2 | 0,590 |

| | | | | | | |
|---|---|---|---|---|---|---|
| Histone H2A type 2-C | HIST2H2AC | 15,56 ± 0,31 | 14,77 ± 0,72 | 0,373 | 14,81 ± 0,77 | 0,417 |
| Coiled-coil domain-containing protein 153 | CCDC153 | 12,44 ± 0,45 | 10,85 ± 0,15 | **0,028** | 11,69 ± 0,63 | 0,383 |
| Probable cation-transporting ATPase 13A5 | ATP13A5 | 11,78 ± 0,24 | 10,88 ± 0,1 | **0,025** | 10,77 ± 0,43 | 0,110 |
| Beta-actin-like protein 2 | ACTBL2 | 13,19 ± 1,68 | 14,65 ± 1,79 | 0,584 | 12,86 ± 1,68 | 0,898 |
| Phosphoinositide 3-kinase regulatory subunit 6 | PIK3R6 | 11,83 ± 0,47 | 10,7 ± 0,08 | 0,080 | 11,06 ± 0,38 | 0,277 |
| Capping protein, Arp2/3 and myosin-I linker protein 2 | RLTPR | 13,69 ± 0,72 | 12,99 ± 1,55 | 0,706 | 12,04 ± 1,26 | 0,321 |
| Histone H3.2 | HIST2H3A | 12,32 ± 0,29 | 13,37 ± 0,05 | **0,025** | 12,33 ± 0,57 | 0,987 |
| Tubulin alpha-1A chain | TUBA1A | 12,24 ± 0,78 | 12,33 ± 0,05 | 0,914 | 11,26 ± 0,47 | 0,342 |
| Abhydrolase domain-containing protein 12B | ABHD12B | 11,93 ± 0,29 | 10,48 ± 0,48 | 0,062 | 11,2 ± 0,96 | 0,508 |
| Mucin-19 | MUC19 | 11,93 ± 0,31 | 10,81 ± 0,13 | **0,030** | 11,1 ± 0,19 | 0,087 |
| Sperm-tail PG-rich repeat-containing protein 2 | STPG2 | 11,67 ± 0,65 | 10,53 ± 0,19 | 0,165 | 10,89 ± 0,42 | 0,371 |
| Carboxypeptidase A6 | CPA6 | 11,94 ± 0,4 | 10,88 ± 0,35 | 0,116 | 10,72 ± 0,37 | 0,089 |
| Junction-mediating and -regulatory protein | JMY | 11,82 ± 0,32 | 10,39 ± 0,46 | 0,065 | 11,06 ± 0,19 | 0,113 |
| Pantothenate kinase 1 | PANK1 | 11,92 ± 0,39 | 11,36 ± 0,32 | 0,327 | 10,93 ± 0,06 | 0,065 |
| Putative uncharacterized protein encoded by LINC00479 | LINC00479 | 15,43 ± 1,52 | 16,88 ± 0,2 | 0,395 | 16,66 ± 0,36 | 0,473 |
| Ferroptosis suppressor protein 1 | AIFM2 | 11,97 ± 0,39 | 11,04 ± 0,14 | 0,087 | 11,04 ± 0,23 | 0,109 |
| Endoplasmic reticulum transmembrane helix translocase | ATP13A1 | 11,52 ± 0,61 | 10,59 ± 0,64 | 0,356 | 12,07 ± 1,14 | 0,693 |
| Uncharacterized protein C11orf16 | C11orf16 | 11,73 ± 0,6 | 10,42 ± 0,16 | 0,103 | 12,4 ± 1,68 | 0,729 |
| Myocardin-related transcription factor B | MKL2 | 11,65 ± 0,56 | 10,84 ± 0,36 | 0,295 | 12,25 ± 1,45 | 0,720 |
| Talin-1 | TLN1 | 11,99 ± 0,46 | 11,34 ± 0,24 | 0,278 | 12,04 ± 1,5 | 0,979 |
| Prostaglandin D2 receptor 2 | PTGDR2 | 11,64 ± 0,66 | 11,27 ± 0,63 | 0,705 | 10,71 ± 0,36 | 0,285 |

*SE: Standard error

**Section 6.** Metabolome analysis

To investigate how NM and PM affect COs in metabolite levels, we performed an untargeted metabolome analysis by GC-MS as described in Section 2. The list of metabolites identified in all three groups were presented in **Table S5.**

*Table S5*. Post-microbial interaction metabolomic changes in COs: The list of the identified metabolites between the groups of Cntrl_COs, NM_COs, and PM_COs

| **Sample** | **CTRL_COs** | **NM_COs** | | **PM_COs** | | **NM_COs vs PM_COs** |
|---|---|---|---|---|---|---|
| **Label** | Mean ± SE | Mean ± SE | p | Mean ± SE | p | p |
| 10-hydroxydecanoic acid | 0.85 ± 0.23 | 1.02 ± 0.14 | 0.57 | 1.14 ± 0.13 | 0.34 | 0.57 |
| 16-glucuronide-estriol | 0.67 ± 0.1 | 1.31 ± 0.08 | **0.01** | 1.02 ± 0.11 | 0.09 | 0.10 |
| 1-methylhydantoin | 1.12 ± 0.08 | 1.01 ± 0.05 | 0.28 | 0.87 ± 0.39 | 0.56 | 0.74 |
| 2-amino-1-phenylethanol | 1.02 ± 0.09 | 0.69 ± 0.08 | 0.05 | 1.29 ± 0.17 | 0.24 | **0.04** |
| 2-Aminobutyric acid | 1.06 ± 0.08 | 0.73 ± 0.36 | 0.42 | 1.22 ± 0.01 | 0.10 | 0.25 |
| 2-aminoethanethiol | 1.22 ± 0.19 | 0.82 ± 0.04 | 0.11 | 0.96 ± 0.02 | 0.24 | **0.04** |
| 2'-deoxyguanosine | 1.27 ± 0.12 | 0.81 ± 0.12 | 0.06 | 0.92 ± 0.12 | 0.11 | 0.54 |
| 2-furoic acid | 1.05 ± 0.12 | 0.80 ± 0.29 | 0.47 | 1.15 ± 0.19 | 0.70 | 0.38 |
| 2-ketoadipic acid | 0.21 ± 0.03 | 1.77 ± 0.51 | **0.04** | 1.02 ± 0.53 | 0.20 | 0.36 |
| 2-ketobutyric acid | 0.14 ± 0.12 | 1.01 ± 0.20 | **0.02** | 1.85 ± 0.12 | **0.00** | **0.02** |
| 2-ketocaproic acid | 1.33 ± 0.19 | 0.68 ± 0.11 | **0.04** | 0.99 ± 0.04 | 0.16 | 0.05 |
| 2-ketoisocaproic acid | 1.17 ± 0.22 | 1.10 ± 0.20 | 0.82 | 0.73 ± 0.31 | 0.31 | 0.38 |
| 3-hydroxypropanoic acid | 0.54 ± 0.41 | 1.70 ± 0.05 | 0.05 | 0.76 ± 0.03 | 0.62 | **0.00** |
| 3-phosphoglyceric acid | 1.00 ± 0.29 | 1.30 ± 0.28 | 0.50 | 0.71 ± 0.20 | 0.47 | 0.16 |
| 4-acetamidobutyric acid | 0.27 ± 0.27 | 1.69 ± 0.89 | 0.20 | 1.04 ± 0.38 | 0.18 | 0.54 |
| 4-guanidinobutyric acid | 1.03 ± 0.08 | 1.01 ± 0.02 | 0.79 | 0.96 ± 0.04 | 0.48 | 0.37 |
| 4-hydroxybenzoic acid | 0.62 ± 0.04 | 1.44 ± 0.08 | **0.00** | 0.95 ± 0.06 | **0.01** | **0.01** |
| 5-aminovaleric acid | 0.44 ± 0.08 | 0.85 ± 0.05 | **0.01** | 1.71 ± 0.18 | **0.00** | **0.01** |
| 5'-deoxy-5'-(methylthio)adenosine | 0.87 ± 0.44 | 2.12 ± 0.67 | 0.19 | 0.01 ± 0.00 | 0.12 | **0.03** |
| 5-hydroxy-L-tryptophan | 2.22 ± 0.49 | 0.33 ± 0.06 | **0.02** | 0.45 ± 0.08 | **0.02** | 0.26 |
| 6-deoxy-D-glucose | 1.21 ± 0.06 | 1.07 ± 0.50 | 0.80 | 0.71 ± 0.28 | 0.16 | 0.56 |
| 6-hydroxy caproic acid | 0.98 ± 0.04 | 0.87 ± 0.01 | 0.06 | 1.15 ± 0.08 | 0.13 | **0.03** |
| 6-phosphogluconic acid | 1.12 ± 0.25 | 0.98 ± 0.15 | 0.67 | 0.90 ± 0.03 | 0.44 | 0.63 |
| acetoacetate | 0.57 ± 0.07 | 0.83 ± 0.43 | 0.57 | 1.60 ± 0.48 | 0.10 | 0.30 |
| acetol | 0.01 ± 0.00 | 1.67 ± 0.06 | **0.00** | 1.32 ± 0.05 | **0.00** | **0.01** |
| acetyl-L-serine | 1.17 ± 0.08 | 0.84 ± 0.41 | 0.49 | 0.99 ± 0.15 | 0.35 | 0.76 |
| adenine | 1.38 ± 0.31 | 0.96 ± 0.18 | 0.30 | 0.66 ± 0.26 | 0.15 | 0.40 |
| adrenaline | 0.88 ± 0.69 | 0.94 ± 0.12 | 0.94 | 1.18 ± 0.57 | 0.76 | 0.69 |
| alpha ketoglutaric acid | 1.06 ± 0.06 | 1.05 ± 0.11 | 0.94 | 0.89 ± 0.05 | 0.10 | 0.24 |
| Asparagine | 0.87 ± 0.04 | 1.03 ± 0.15 | 0.35 | 1.10 ± 0.16 | 0.23 | 0.79 |
| aspartic acid | 0.49 ± 0.02 | 1.41 ± 0.06 | **0.00** | 1.10 ± 0.02 | **0.00** | **0.01** |
| Beta- alanine | 2.06 ± 0.43 | 0.68 ± 0.11 | **0.04** | 0.26 ± 0.09 | **0.01** | **0.04** |
| beta-glutamic acid | 1.39 ± 0.50 | 0.86 ± 0.33 | 0.43 | 0.75 ± 0.34 | 0.35 | 0.83 |
| capric acid | 1.10 ± 0.09 | 0.79 ± 0.01 | **0.02** | 1.11 ± 0.11 | 0.97 | **0.04** |
| cis-Aconitic acid | 0.80 ± 0.03 | 1.40 ± 0.76 | 0.47 | 0.80 ± 0.30 | 1.00 | 0.50 |
| citramalic acid | 0.49 ± 0.04 | 1.96 ± 1.55 | 0.40 | 0.55 ± 0.03 | 0.32 | 0.41 |
| Citric acid | 1.07 ± 0.31 | 0.95 ± 0.07 | 0.71 | 0.98 ± 0.18 | 0.81 | 0.87 |
| citrulline | 0.40 ± 0.06 | 1.85 ± 0.19 | **0.00** | 0.75 ± 0.27 | 0.26 | **0.03** |

| | | | | | | |
|---|---|---|---|---|---|---|
| Creatinine | 1.70 ± 0.11 | 1.05 ± 0.14 | **0.02** | 0.26 ± 0.15 | **0.00** | **0.02** |
| cysteinylglycine | 1.04 ± 0.09 | 0.77 ± 0.03 | **0.04** | 1.19 ± 0.14 | 0.40 | **0.04** |
| cytosine | 0.42 ± 0.05 | 1.88 ± 0.59 | 0.07 | 0.70 ± 0.16 | 0.18 | 0.12 |
| D-(+)-melezitose | 1.16 ± 0.12 | 1.16 ± 0.13 | 0.99 | 0.68 ± 0.07 | **0.03** | **0.03** |
| D-Ala-D-Ala | 0.49 ± 0.02 | 1.22 ± 0.07 | **0.00** | 1.29 ± 0.19 | **0.01** | 0.74 |
| D-Glucose-6-phosphoric acid | 0.59 ± 0.01 | 1.05 ± 0.05 | **0.00** | 1.36 ± 0.09 | **0.00** | **0.05** |
| DL-3-aminoisobutyric acid | 1.10 ± 0.02 | 0.97 ± 0.08 | 0.21 | 0.93 ± 0.15 | 0.35 | 0.83 |
| DL-dihydrosphingosine | 0.91 ± 0.02 | 1.46 ± 0.17 | **0.03** | 0.63 ± 0.10 | 0.07 | **0.02** |
| DL-glyceraldehyde | 1.16 ± 0.54 | 0.57 ± 0.03 | 0.33 | 1.27 ± 0.59 | 0.90 | 0.30 |
| DL-isoleucine | 0.96 ± 0.06 | 0.94 ± 0.04 | 0.83 | 1.09 ± 0.01 | 0.11 | **0.03** |
| Malic acid | 0.86 ± 0.04 | 1.35 ± 0.01 | **0.00** | 0.80 ± 0.02 | 0.23 | **0.00** |
| D-saccharic acid | 0.98 ± 0.48 | 1.38 ± 0.1 | 0.46 | 0.64 ± 0.23 | 0.57 | **0.04** |
| D-sorbitol | 1.59 ± 0.25 | 0.88 ± 0.04 | **0.05** | 0.53 ± 0.06 | **0.01** | **0.01** |
| D-sphingosine | 1.36 ± 0.06 | 0.97 ± 0.11 | **0.04** | 0.67 ± 0.34 | 0.12 | 0.45 |
| fructose | 0.18 ± 0.01 | 2.23 ± 0.19 | **0.00** | 0.59 ± 0.01 | **0.00** | **0.00** |
| fumaric acid | 1.12 ± 0.06 | 1.15 ± 0.04 | 0.76 | 0.73 ± 0.03 | **0.00** | **0.00** |
| galactinol | 0.45 ± 0.08 | 2.07 ± 1.30 | 0.28 | 0.47 ± 0.18 | 0.93 | 0.29 |
| gluconic acid | 1.55 ± 0.01 | 0.96 ± 0.29 | 0.11 | 0.49 ± 0.13 | **0.00** | 0.22 |
| glucuronic acid | 0.11 ± 0.04 | 2.10 ± 0.84 | 0.08 | 0.79 ± 0.15 | **0.01** | 0.20 |
| Glutamic acid | 1.02 ± 0.23 | 1.34 ± 0.36 | 0.50 | 0.64 ± 0.27 | 0.34 | 0.19 |
| glyceric acid | 0.20 ± 0.11 | 1.65 ± 1.04 | 0.24 | 1.15 ± 1.04 | 0.42 | 0.75 |
| glycerol | 0.73 ± 0.00 | 1.14 ± 0.01 | **0.00** | 1.14 ± 0.03 | **0.00** | 0.97 |
| glycerol 1-phosphate | 0.26 ± 0.05 | 2.20 ± 1.88 | 0.36 | 0.55 ± 0.03 | **0.01** | 0.43 |
| Glycerone phosphoric acid | 0.41 ± 0.03 | 1.65 ± 0.33 | **0.02** | 0.94 ± 0.22 | 0.08 | 0.15 |
| Glycine 2TMS | 0.91 ± 0.16 | 1.21 ± 0.44 | 0.56 | 0.88 ± 0.15 | 0.87 | 0.51 |
| glycolic acid | 1.38 ± 0.23 | 0.12 ± 0.07 | **0.01** | 1.50 ± 0.78 | 0.89 | 0.15 |
| hyoscyamine | 1.08 ± 0.20 | 0.78 ± 0.01 | 0.21 | 1.15 ± 0.28 | 0.85 | 0.27 |
| hypotaurine | 2.03 ± 0.16 | 0.70 ± 0.02 | **0.00** | 0.27 ± 0.07 | **0.00** | **0.01** |
| hypoxanthine | 1.21 ± 0.20 | 0.89 ± 0.06 | 0.20 | 0.90 ± 0.16 | 0.29 | 0.97 |
| Indole-3-acetaldehyde | 1.05 ± 0.04 | 0.82 ± 0.07 | **0.05** | 1.13 ± 0.07 | 0.34 | **0.03** |
| inosine | 0.50 ± 0.01 | 1.93 ± 0.12 | **0.00** | 0.57 ± 0.18 | 0.69 | **0.00** |
| Isoleucine | 1.05 ± 0.14 | 1.11 ± 0.24 | 0.84 | 0.83 ± 0.15 | 0.35 | 0.38 |
| isomaltose | 1.14 ± 0.48 | 0.98 ± 0.05 | 0.75 | 0.88 ± 0.04 | 0.61 | 0.19 |
| itaconic acid | 0.86 ± 0.11 | 1.27 ± 0.3 | 0.26 | 0.87 ± 0.12 | 0.94 | 0.28 |
| L- sorbose | 1.26 ± 0.41 | 1.59 ± 0.67 | 0.70 | 0.14 ± 0.07 | 0.05 | 0.10 |
| L-(+) lactic acid | 0.92 ± 0.12 | 0.82 ± 0.40 | 0.83 | 1.27 ± 0.14 | 0.14 | 0.35 |
| lactamide | 0.81 ± 0.31 | 0.70 ± 0.49 | 0.86 | 1.49 ± 0.51 | 0.32 | 0.33 |
| Lactic acid | 1.04 ± 0.12 | 1.06 ± 0.20 | 0.94 | 0.90 ± 0.20 | 0.58 | 0.60 |
| lactobionic acid | 0.60 ± 0.17 | 2.22 ± 0.26 | **0.01** | 0.18 ± 0.05 | 0.08 | **0.00** |
| lactose | 1.54 ± 0.22 | 0.90 ± 0.09 | 0.05 | 0.55 ± 0.15 | **0.02** | 0.12 |
| lactulose | 0.50 ± 0.04 | 2.14 ± 0.05 | **0.00** | 0.37 ± 0.02 | **0.04** | **0.00** |
| L-alanine | 0.75 ± 0.13 | 0.93 ± 0.23 | 0.53 | 1.32 ± 0.07 | **0.02** | 0.18 |
| L-allothreonine | 0.60 ± 0.02 | 0.99 ± 0.06 | **0.00** | 1.41 ± 0.10 | **0.00** | **0.02** |

| | | | | | | |
|---|---|---|---|---|---|---|
| L-ascorbic acid | 1.04 ± 0.06 | 1.23 ± 0.14 | 0.28 | 0.73 ± 0.20 | 0.22 | 0.11 |
| lauric acid | 0.77 ± 0.05 | 0.90 ± 0.11 | 0.36 | 1.33 ± 0.31 | 0.15 | 0.25 |
| L-cysteine | 1.20 ± 0.52 | 0.83 ± 0.17 | 0.53 | 0.97 ± 0.09 | 0.69 | 0.49 |
| L-cystine | 1.02 ± 0.13 | 1.26 ± 0.26 | 0.45 | 0.72 ± 0.26 | 0.37 | 0.22 |
| L-dithiothreitol | 1.10 ± 0.11 | 0.83 ± 0.06 | 0.09 | 1.07 ± 0.11 | 0.82 | 0.12 |
| Leucine | 1.12 ± 0.08 | 1.05 ± 0.42 | 0.87 | 0.83 ± 0.12 | 0.10 | 0.64 |
| L-glutamine | 0.92 ± 0.11 | 0.89 ± 0.02 | 0.80 | 1.19 ± 0.02 | 0.07 | **0.00** |
| L-homocystine | 0.24 ± 0.06 | 2.47 ± 0.26 | **0.00** | 0.29 ± 0.12 | 0.73 | **0.00** |
| L-homoserine | 1.29 ± 0.05 | 1.03 ± 0.08 | **0.05** | 0.68 ± 0.06 | **0.00** | **0.02** |
| L-methionine | 0.07 ± 0.00 | 2.86 ± 0.18 | **0.00** | 0.07 ± 0.01 | 0.78 | **0.00** |
| L-ornithine | 0.26 ± 0.04 | 2.67 ± 0.19 | **0.00** | 0.08 ± 0.01 | **0.02** | **0.00** |
| L-proline | 0.14 ± 0.03 | 0.11 ± 0.08 | 0.77 | 2.74 ± 2.56 | 0.37 | 0.36 |
| L-pyroglutamic acid | 1.27 ± 0.02 | 0.87 ± 0.04 | **0.00** | 0.86 ± 0.05 | **0.00** | 0.86 |
| L-threonine | 1.46 ± 0.28 | 1.01 ± 0.37 | 0.39 | 0.53 ± 0.33 | 0.10 | 0.39 |
| L-tyrosine | 1.12 ± 0.12 | 1.17 ± 0.14 | 0.77 | 0.71 ± 0.27 | 0.24 | 0.20 |
| Lysine | 0.92 ± 0.05 | 1.17 ± 0.04 | **0.02** | 0.91 ± 0.03 | 0.94 | **0.01** |
| malonic acid | 0.66 ± 0.01 | 1.33 ± 0.12 | **0.00** | 1.00 ± 0.05 | **0.00** | 0.06 |
| maltotriose | 1.16 ± 0.13 | 1.14 ± 0.13 | 0.91 | 0.7 ± 0.07 | **0.03** | **0.04** |
| melatonin | 0.74 ± 0.09 | 2.10 ± 0.35 | **0.02** | 0.16 ± 0.13 | **0.02** | **0.01** |
| melibiose | 1.15 ± 0.18 | 1.33 ± 0.23 | 0.56 | 0.52 ± 0.19 | 0.07 | **0.05** |
| methoxytryptamine | 0.99 ± 0.11 | 0.55 ± 0.09 | **0.04** | 1.46 ± 0.20 | 0.11 | **0.01** |
| methyl-beta-D-galactopyranoside | 0.50 ± 0.03 | 1.61 ± 0.14 | **0.00** | 0.89 ± 0.17 | 0.09 | **0.03** |
| methylmalonic acid | 1.24 ± 0.14 | 0.93 ± 0.10 | 0.14 | 0.83 ± 0.20 | 0.17 | 0.68 |
| mucic acid | 0.89 ± 0.21 | 1.60 ± 0.59 | 0.32 | 0.51 ± 0.19 | 0.25 | 0.15 |
| N-(2-hydroxyethyl)iminodiacetic acid | 0.10 ± 0.02 | 2.52 ± 0.1 | **0.00** | 0.38 ± 0.05 | **0.01** | **0.00** |
| N-acetyl-5-hydroxytryptamine | 1.48 ± 0.65 | 0.71 ± 0.13 | 0.31 | 0.81 ± 0.11 | 0.36 | 0.60 |
| N-acetyl-D-mannosamine | 0.77 ± 0.38 | 1.14 ± 0.05 | 0.39 | 1.09 ± 0.10 | 0.46 | 0.68 |
| N-acetyl-L-aspartic acid | 1.31 ± 0.18 | 0.57 ± 0.11 | **0.02** | 1.13 ± 0.15 | 0.49 | **0.04** |
| N-acetyl-L-glutamic acid | 0.57 ± 0.10 | 1.15 ± 0.23 | 0.08 | 1.28 ± 0.51 | 0.25 | 0.83 |
| N-acetyl-L-leucine | 0.74 ± 0.17 | 0.99 ± 0.33 | 0.53 | 1.27 ± 0.44 | 0.33 | 0.64 |
| N-Acetyl-L-serine | 1.00 ± 0.04 | 0.80 ± 0.06 | **0.05** | 1.20 ± 0.15 | 0.28 | 0.07 |
| N-carbamyl-L-glutamic acid | 1.03 ± 0.08 | 1.01 ± 0.02 | 0.86 | 0.96 ± 0.04 | 0.52 | 0.33 |
| nicotinamide | 1.42 ± 0.24 | 0.38 ± 0.06 | **0.01** | 1.20 ± 0.50 | 0.71 | 0.18 |
| nicotinic acid | 1.61 ± 0.51 | 0.72 ± 0.02 | 0.16 | 0.67 ± 0.24 | 0.17 | 0.85 |
| N-methylalanine | 0.95 ± 0.07 | 0.95 ± 0.04 | 0.96 | 1.1 ± 0.01 | 0.10 | **0.02** |
| Normetadrenaline | 0.67 ± 0.04 | 1.31 ± 0.34 | 0.14 | 1.03 ± 0.3 | 0.30 | 0.57 |
| norvaline | 1.67 ± 0.21 | 0.50 ± 0.25 | **0.02** | 0.83 ± 0.44 | 0.16 | 0.55 |
| O-benzyl-L-tyrosine | 0.54 ± 0.53 | 1.24 ± 0.62 | 0.44 | 1.22 ± 0.13 | 0.28 | 0.97 |
| O-phosphocolamine | 0.19 ± 0.05 | 1.93 ± 0.20 | **0.00** | 0.88 ± 0.27 | 0.06 | **0.03** |
| O-phospho-L-threonine | 1.25 ± 0.18 | 0.91 ± 0.04 | 0.14 | 0.84 ± 0.07 | 0.10 | 0.41 |
| orotic acid | 1.30 ± 0.03 | 0.59 ± 0.09 | **0.00** | 1.11 ± 0.15 | 0.29 | **0.04** |
| oxalacetic acid | 0.94 ± 0.02 | 0.83 ± 0.10 | 0.33 | 1.23 ± 0.1 | **0.05** | **0.05** |

| | | | | | | |
|---|---|---|---|---|---|---|
| oxalic acid | 1.08 ± 0.03 | 0.63 ± 0.23 | 0.12 | 1.28 ± 0.13 | 0.21 | 0.07 |
| palmitic acid | 0.96 ± 0.13 | 0.92 ± 0.05 | 0.79 | 1.11 ± 0.19 | 0.54 | 0.37 |
| pantothenic acid | 1.10 ± 0.08 | 0.87 ± 0.07 | 0.10 | 1.03 ± 0.08 | 0.57 | 0.22 |
| Phenylalanine | 0.81 ± 0.04 | 1.14 ± 0.05 | **0.01** | 1.05 ± 0.07 | **0.04** | 0.30 |
| phenylpyruvate | 1.20 ± 0.52 | 0.74 ± 0.20 | 0.45 | 1.06 ± 0.35 | 0.83 | 0.47 |
| Pi | 0.93 ± 0.12 | 0.49 ± 0.47 | 0.41 | 1.58 ± 0.10 | **0.01** | 0.08 |
| pipecolic acid | 0.56 ± 0.05 | 1.32 ± 0.08 | **0.00** | 1.12 ± 0.07 | **0.00** | 0.14 |
| purine riboside | 1.36 ± 0.30 | 0.75 ± 0.32 | 0.24 | 0.88 ± 0.04 | 0.19 | 0.72 |
| Putrescine | 1.87 ± 0.12 | 0.46 ± 0.11 | **0.00** | 0.68 ± 0.08 | **0.00** | 0.18 |
| pyrophosphate | 0.73 ± 0.38 | 1.76 ± 0.29 | 0.10 | 0.52 ± 0.2 | 0.65 | **0.02** |
| pyruvic acid | 1.86 ± 0.18 | 0.46 ± 0.08 | **0.00** | 0.68 ± 0.05 | **0.00** | 0.07 |
| ribitol | 0.83 ± 0.13 | 1.39 ± 0.04 | **0.01** | 0.78 ± 0.03 | 0.74 | **0.00** |
| ribulose-5-phosphate 3 | 0.66 ± 0.14 | 1.04 ± 0.12 | 0.11 | 1.29 ± 0.15 | **0.04** | 0.27 |
| sarcosine | 0.00 ± 0.00 | 3.00 ± 3.00 | 0.37 | 0.00 ± 0.00 | 0.09 | 0.37 |
| Serine | 0.88 ± 0.01 | 0.94 ± 0.02 | **0.04** | 1.18 ± 0.03 | **0.00** | **0.00** |
| serotonin | 0.34 ± 0.09 | 2.17 ± 0.20 | **0.00** | 0.49 ± 0.07 | 0.25 | **0.00** |
| Shikimic acid | 1.01 ± 0.05 | 0.82 ± 0.07 | 0.08 | 1.18 ± 0.14 | 0.31 | 0.08 |
| spermidine | 0.23 ± 0.03 | 2.34 ± 1.15 | 0.14 | 0.42 ± 0.25 | 0.49 | 0.18 |
| spermine | 0.94 ± 0.21 | 1.78 ± 0.91 | 0.42 | 0.28 ± 0.18 | 0.07 | 0.18 |
| stearic acid | 0.96 ± 0.11 | 0.94 ± 0.05 | 0.90 | 1.10 ± 0.13 | 0.46 | 0.32 |
| succinic acid | 0.68 ± 0.05 | 1.39 ± 0.09 | **0.00** | 0.93 ± 0.07 | **0.04** | **0.02** |
| tagatose | 1.04 ± 0.07 | 1.06 ± 0.04 | 0.81 | 0.90 ± 0.05 | 0.17 | 0.05 |
| tartaric acid | 0.00 ± 0.00 | 2.97 ± 0.18 | **0.00** | 0.02 ± 0.02 | 0.26 | **0.00** |
| tetratriacontane | 0.96 ± 0.26 | 0.95 ± 0.04 | 0.95 | 1.09 ± 0.31 | 0.76 | 0.66 |
| threose | 0.78 ± 0.02 | 0.93 ± 0.45 | 0.75 | 1.29 ± 0.01 | **0.00** | 0.47 |
| thymine | 0.90 ± 0.21 | 1.54 ± 1.14 | 0.61 | 0.56 ± 0.42 | 0.50 | 0.46 |
| trans-4-hydroxy-L-proline | 2.07 ± 0.08 | 0.45 ± 0.02 | **0.00** | 0.47 ± 0.03 | **0.00** | 0.65 |
| trehalose-6-phosphate | 0.48 ± 0.11 | 1.59 ± 0.13 | **0.00** | 0.93 ± 0.14 | 0.06 | **0.02** |
| uracil | 0.24 ± 0.06 | 1.74 ± 0.49 | **0.04** | 1.03 ± 0.56 | 0.23 | 0.40 |
| urea | 1.08 ± 0.04 | 0.79 ± 0.04 | **0.01** | 1.13 ± 0.12 | 0.69 | **0.05** |
| uridine 5'-monophosphate | 1.12 ± 0.04 | 0.90 ± 0.15 | 0.23 | 0.98 ± 0.03 | 0.05 | 0.61 |
| Valine | 1.52 ± 0.14 | 0.61 ± 0.02 | **0.00** | 0.86 ± 0.17 | **0.04** | 0.21 |

**Section 7**. Metabolic flux analysis - Fluxomics

We performed $^{18}$O-SIL strategy for the first time in this study to track the metabolites passing from NM or PM to COs, which might cause cell stress or degeneration, and thus, help us understand the mechanism of pathogenic microbiota-driven neurodegeneration. Notes describing the sample preparation, derivatization, and instrumental analysis were provided below.

**Sample Preparation:** To determine the optimal labelling time, bacteria were incubated in 30% $H_2[^{18}O]$ for 30 min, 4 h, and 24 h, and were analysed under optimized GC-MS conditions. The metabolite extracts were completely dried in a vacuum dryer concentrator (Labconco Refrigerated CentriVap Vacuum Concentrator, Kansas City, USA) at 4 °C to avoid degradation.

**Derivatization:** The derivatization protocol was performed as described in our previous study [24]. Briefly, the dried samples were methoxyaminated by 20 µL MeOX (20 mg/mL in pyridine) for 90 min at 30 °C and silylated by 80 µL MSTFA + 1% TMCS for 120 minutes at 37 °C. Following, samples were transferred to silylated vials.

**GC-MS Analysis:** Derivatized samples were analyzed using DB-5MS (30 m + 10 m duraguard × 0.25 mm i.d. and 0.25-µm) capillary column by GC-MS (Shimadzu-QP2010 Ultra, Kyoto, Japan). In a splitless injection, the injection volume was set at 1 µL. The column temperature was initially maintained at 60 °C for 1 min, then increased from 60 to 325 °C at 5 °C/min, and maintained for 6 min. At a flow rate of 0.99 mL/min, the carrier gas was high-purity helium (>%99.999). The injector temperature was set to 250 °C. The MS parameters were as respectively; ion source and interface temperature, 230 °C and 290 °C; solvent delay time, 7.5 min. Data were scanned over the range of m/z of 50-650 at 0.30 sec event time.

**Labelling time optimization**: Before setting up the co-culture experiments of COs and NM or PM, we optimized the labelling time of $H_2[^{18}O]$ for NM and PM. The labelling efficiency of inorganic phosphate (Pi) and succinate at 0.5 h, 4 h, and 24 h was presented in **Figure S17**.

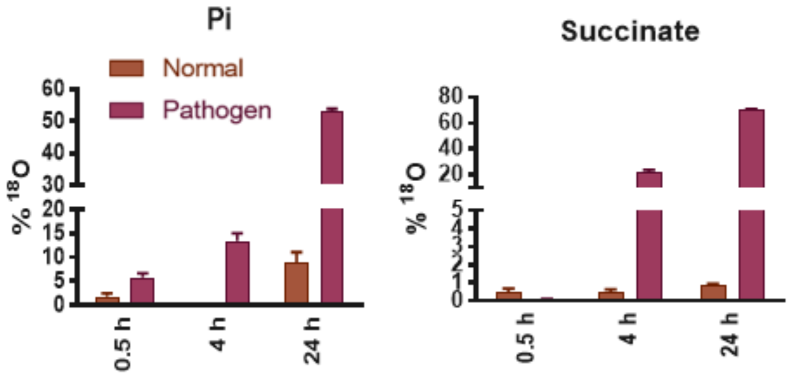

*Figure 17.* Labelling optimisation for $H_2[^{18}O]$ at different time points: The labelling efficiency of Pi and Succinate

**Calculation of $^{18}$O-labelled metabolites**: In essence, each oxygen ($^{16}O$) in the metabolite that was labelled with $H_2[^{18}O]$ induces a 2 AMU increase in mass, and thus in fragments. Consequently, the $^{18}O$ labelling ratios of metabolites are calculated by analysing the peak areas at each AMU (isotopologue distribution). We determined the $^{18}O$ labelling ratios using the following equation:

$$\text{Total } {}^{18}\text{O labeling percentage} = \left(\sum_{i=1}^{n} i \times {}^{18}\text{O}_i\%\right) / \, n \times \text{H}_2[{}^{18}\text{O}]\,\%$$

$n$: Total labelled oxygen in the molecule

$i$: Number of the labelled oxygen with $H_2[^{18}O]$

Before proceeding with the co-culture experiments, we further re-examined the growth kinetics of strains before and after labelling with $H_2[^{18}O]$ to investigate the possible adverse effects of $H_2[^{18}O]$ on strains. The growth kinetics of NM and PM before and after labelling with $H_2[^{18}O]$ were presented in **Figure S18**.

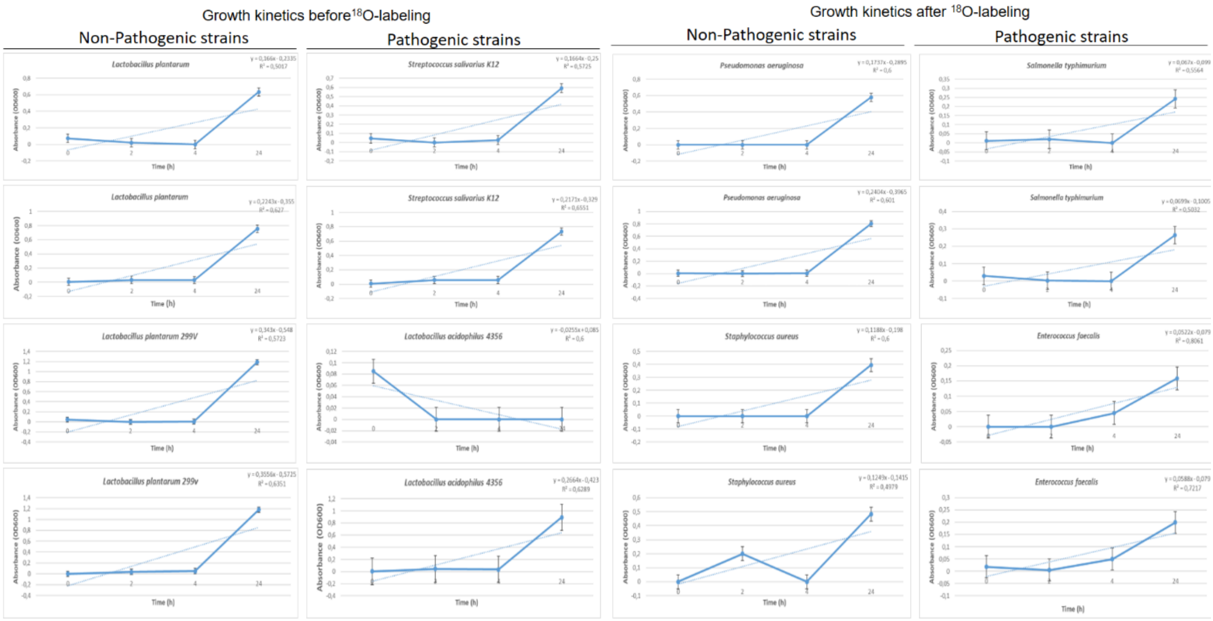

*Figure S18*. Growth kinetics of NM and PM before and after labelling with $H_2[^{18}O]$ in 24-well plates

The fragment ions (m/z) for mass monitoring and calculation of $^{18}O$ labelling ratios in COs after co-cultured with NM or PM were presented in **Table S6**.

*Table S6*. The fragment ions (m/z) monitored for the calculation of the $^{18}O$ labelling ratios

| Metabolite | $^{16}O$ | $^{18}O_1$ | $^{18}O_2$ | $^{18}O_3$ | $^{18}O_4$ | $^{18}O_5$ | $^{18}O_6$ | $^{18}O_7$ |
|---|---|---|---|---|---|---|---|---|
| 2-ketoisocaproic acid | 216 | 218 | 220 | 222 | | | | |
| 3-phosphoglycerate | 357 | 359 | 361 | 363 | 365 | 367 | 369 | 371 |
| allo-inositol | 318 | 320 | 322 | 324 | 326 | 328 | 330 | |
| Arachidic Acid | 369 | 371 | 373 | | | | | |
| Aspartic Acid | 245 | 247 | 249 | 251 | 253 | | | |
| Beta-Alanine | 248 | 250 | 252 | | | | | |
| Citric acid | 347 | 349 | 351 | 353 | 355 | 357 | 359 | 361 |
| Creatinine | 329 | 331 | | | | | | |
| D-glucose | 319 | 321 | 323 | 325 | 327 | 329 | 331 | |
| Glucose-6-phosphate | 387 | 389 | 391 | 393 | 395 | 397 | 399 | 401 |
| D-ribulose 5-phosphate | 357 | 359 | 361 | 363 | 365 | 367 | 369 | 371 |
| D-threitol | 217 | 219 | 221 | 223 | 225 | | | |
| Dihydroxyacetone phosphate | 400 | 402 | 404 | 406 | 408 | 410 | 412 | |
| L-isoleucine | 188 | 190 | 192 | | | | | |
| Fumaric acid | 245 | 247 | 249 | 251 | 253 | | | |
| Gluconic Acid | 333 | 335 | 337 | 339 | 341 | 343 | 345 | 347 |
| Glyceric Acid | 292 | 294 | 296 | 298 | 300 | | | |
| Glycerol | 205 | 207 | 209 | 211 | | | | |

| | | | | | | | |
|---|---|---|---|---|---|---|---|
| Glycerol-1-phosphate | 357 | 359 | 361 | 363 | 365 | 367 | 369 |
| Glycine | 248 | 250 | 252 | | | | |
| Glycolic acid | 177 | 179 | 181 | 183 | | | |
| Hypotaurine | 188 | 190 | 192 | | | | |
| Lactic acid | 219 | 221 | 223 | 225 | | | |
| L-glutamic acid | 230 | 232 | 234 | 236 | 238 | | |
| L-leucine | 188 | 190 | 192 | | | | |
| L-proline | 172 | 174 | 176 | | | | |
| L-pyroglutamic acid | 230 | 232 | 234 | 236 | | | |
| L-serine | 219 | 221 | 223 | 225 | | | |
| L-threonine | 219 | 221 | 223 | 225 | | | |
| L-tyrosine | 208 | 210 | 212 | 214 | | | |
| L-valine | 174 | 176 | 178 | | | | |
| Malic acid | 233 | 235 | 237 | 239 | 241 | 243 | |
| Malonic Acid | 233 | 235 | 237 | 239 | 241 | | |
| N-acetyl-D-mannosamine | 319 | 321 | 323 | 325 | 327 | 329 | 331 |
| Palmitic Acid | 313 | 315 | 317 | | | | |
| Phenylalanine | 222 | 224 | 226 | | | | |
| Pi | 299 | 301 | 303 | 305 | 307 | | |
| Stearic Acid | 341 | 343 | 345 | | | | |
| Succinic acid | 247 | 249 | 251 | 253 | 255 | | |
| Tartaric acid | 292 | 294 | 296 | 298 | 300 | 302 | 304 |
| Uracil | 241 | 243 | 245 | | | | |

The ratios of $^{18}O$-labelled metabolites were classified as carbohydrate, amino acid, and lipid metabolites, and presented in **Figure S19-21**. For each metabolite, the sum of the labelling ratio of the bacterial population and the labelled metabolite ratio in the organoids gives the total labelling percentage for that metabolite. For example, the sum of the labelling percentages of the pathogenic population (approx. 5%) and PM_COs (approx. 15%) for G6P is approximately 20%. After the co-culture period, the labeled G6P ratio remained at 5% in the pathogenic population, while it was around 15% in PM_COs. This result suggests that a 15% part of 20% of the $^{18}O$-labelled G6P was transported to COs.

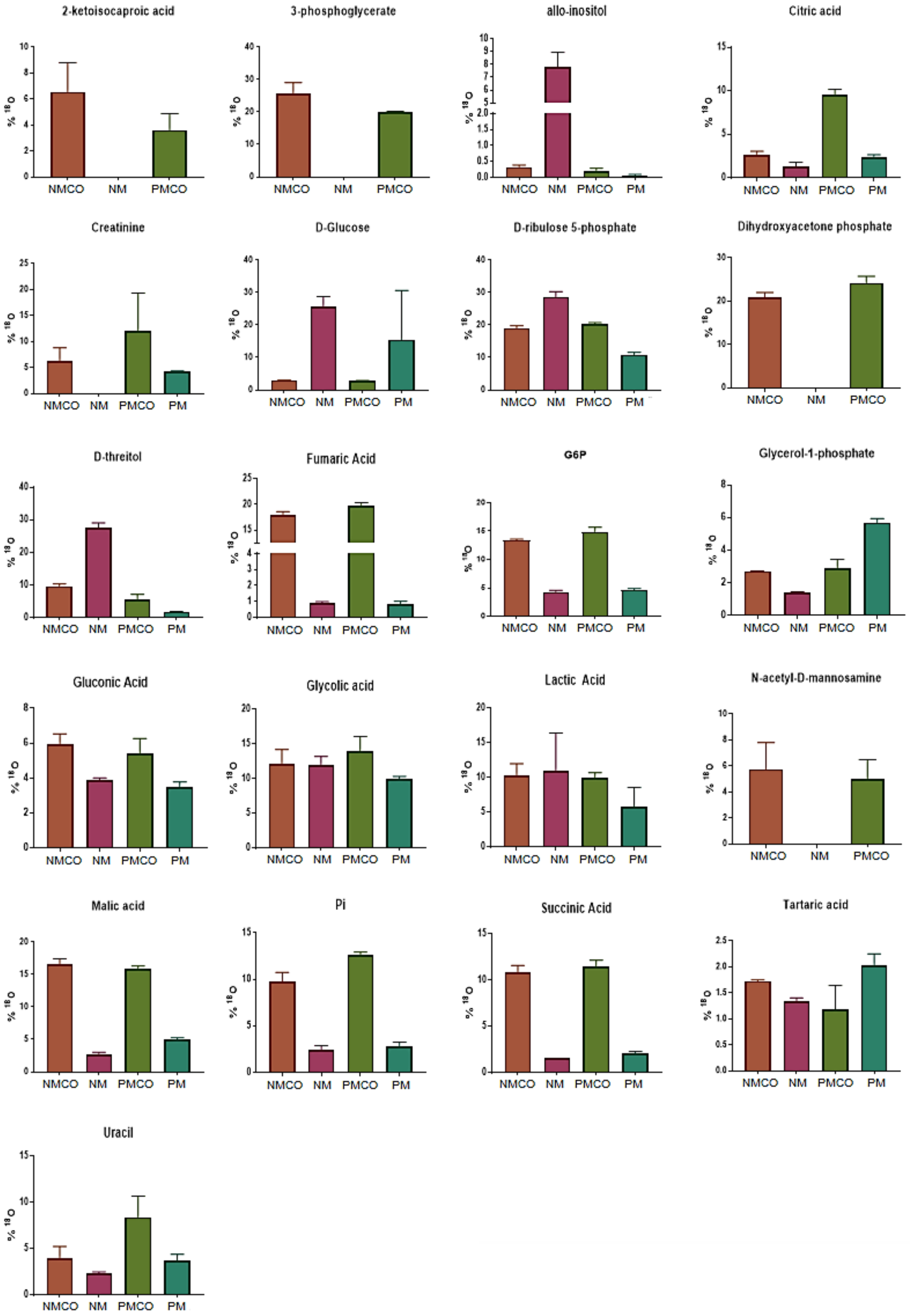

*Figure 19*. Central carbon metabolism related metabolites diffusing from NM or PM to COs, determined by $^{18}$O-SIL-based metabolic flux analysis. 30 COs were used in each sample, and three repeats were used in each group.

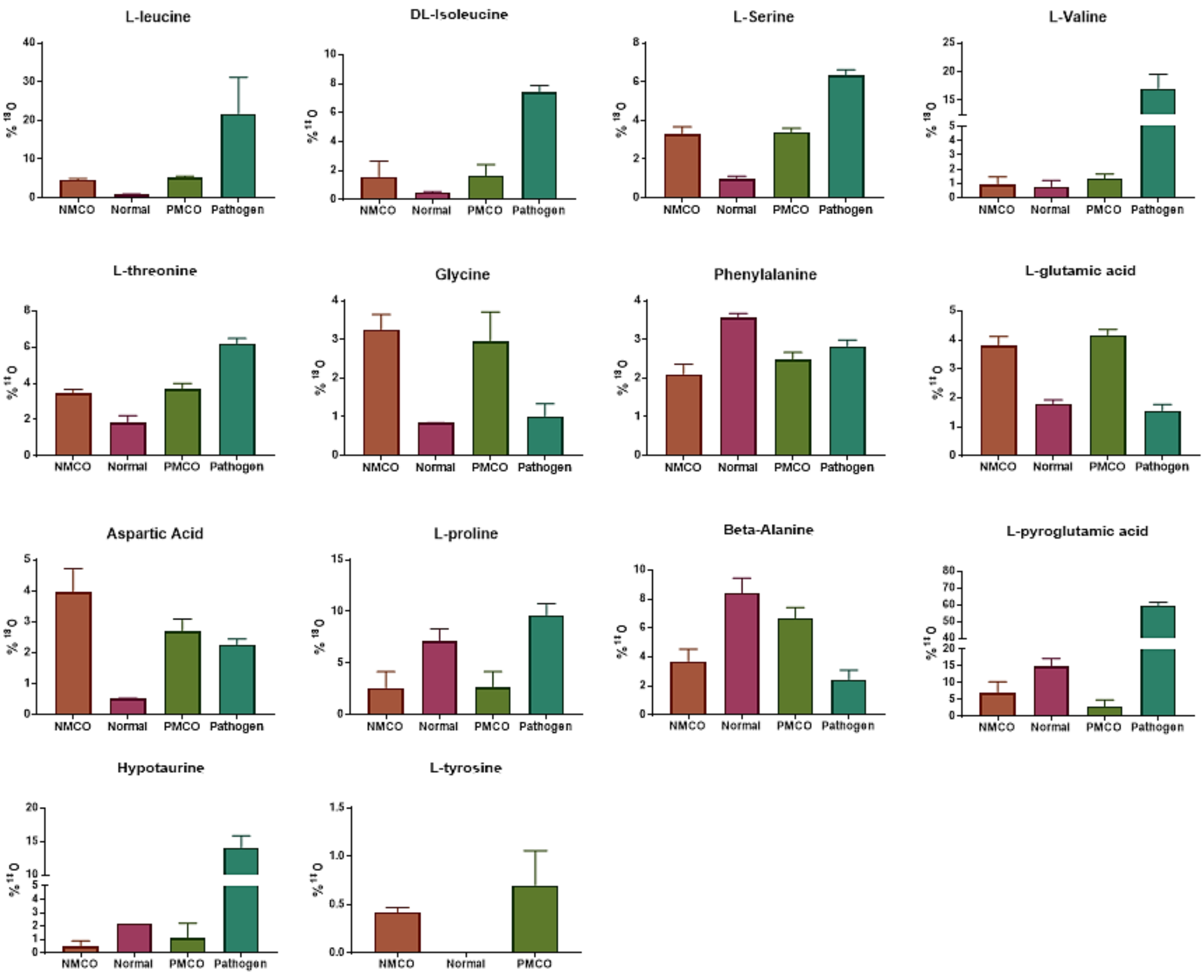

*Figure 20*. Amino acid metabolism related metabolites diffusing from NM or PM to COs, determined by $^{18}$O-SIL-based metabolic flux analysis. 30 COs were used in each sample, and three repeats were used in each group.

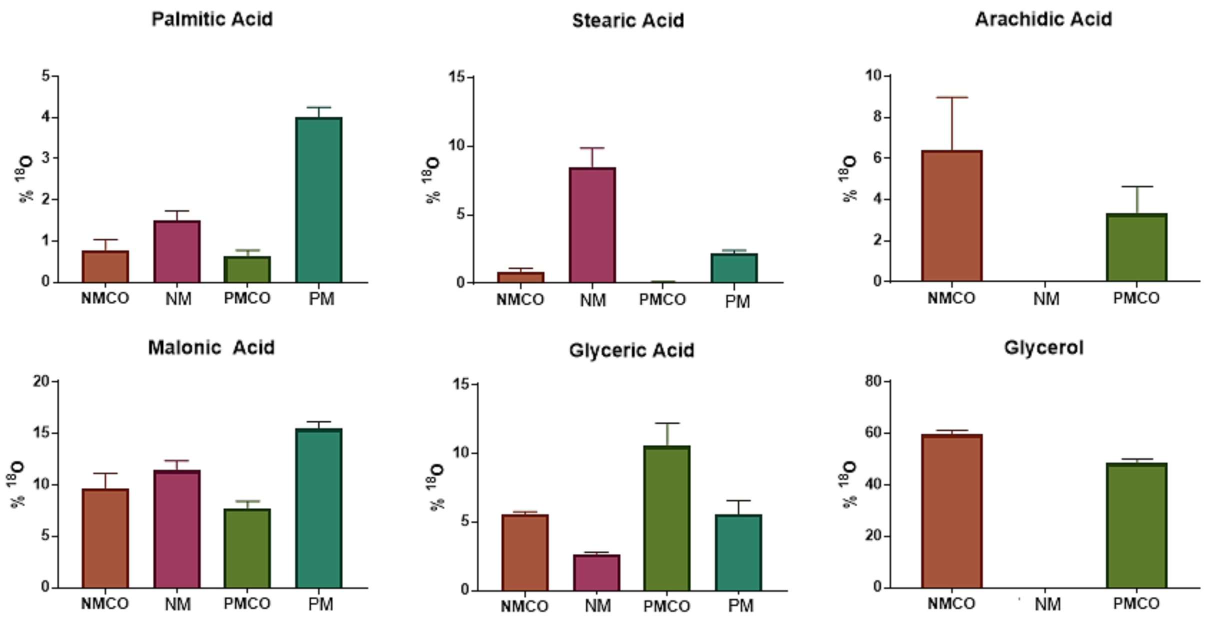

*Figure 21*. Lipid metabolism related metabolites diffusing from NM or PM to COs, determined by $^{18}O$-SIL-based metabolic flux analysis. 30 COs were used in each sample, and three repeats were used in each group.
